# Distinct development-associated roles of rice histone variant H2A.X in suppressing deposition of active H3K4me3 marks and in restricting H2A.W incorporation

**DOI:** 10.64898/2026.02.24.707635

**Authors:** Aravind Madhu, Vivek Hari-Sundar Gandhivel, Steffi Raju, Riju Dey, P.V. Shivaprasad

## Abstract

Histone variant H2A.X is a well-conserved histone that plays crucial roles in mediating DNA damage response across eukaryotes. Although H2A.X expresses even without any stress, and decorates gene bodies of actively expressed genes, it is not known if H2A.X has functions beyond DNA damage repair. Using genetic, high throughput genomics and molecular approaches, we identified a previously unappreciated role of H2A.X in regulating development-associated genes. Using custom-made antibodies specific to H2A.X variant, we show that it suppressed the deposition of active H3K4me3 marks over gene bodies and Transposable elements (TE)s, specifically regulating several root development, photosynthesis, and pigmentation-related genes as seen by the impairment of these processes in h2a.x ko (knockout) plants. H2A.X also suppressed global deposition of repressive mark H3K9me2 by restricting activity of H2A variant H2A.W. In agreement with this, there was a genome-wide re-localization of H2A.W to TEs and a few genes in h2a.x ko plants. H2A.X overexpressing plants exhibited stress phenotypes including increased anthocyanin levels, mimicking the transcriptome of DNA damaged wildtype plants. The transcriptome of kd lines of FACT complex, a known chaperone of H2A.X, was largely similar to that of h2a.x ko, suggesting that the development-associated functions of FACT are at least partially due to H2A.X. These results suggest a key role of H2A.X in regulating the competing histone marks and this function might be conserved across plants.

**Graphical Abstract:** 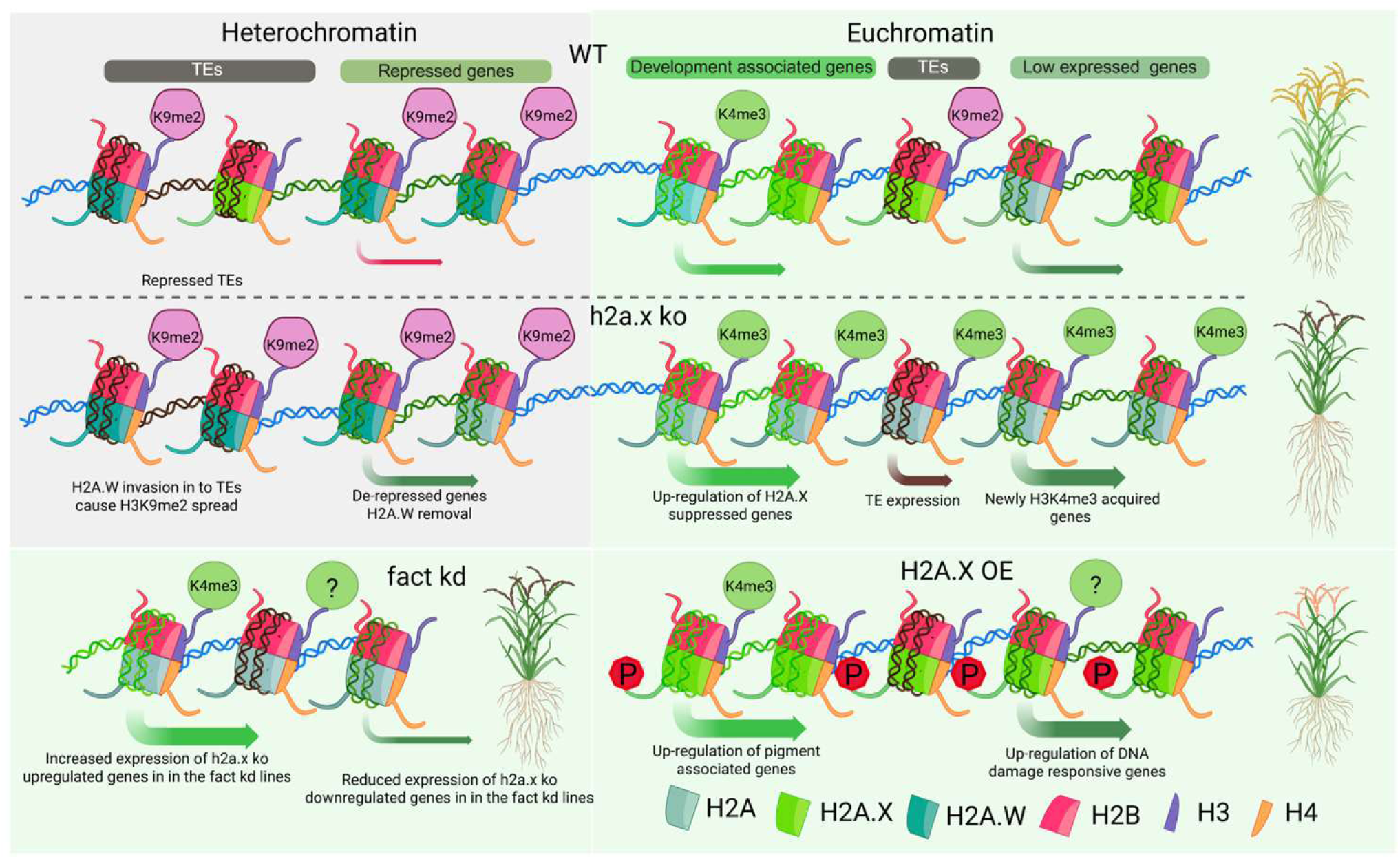

## Introduction

Histone octamers are made of a (H3-H4)_2_ tetramer and two dimers consisting of one each of H2A and H2B proteins (1, 2). Regulatory activity of histones largely comes from their ability to have often conserved one or more post-translational modifications (PTMs) predominantly in their N-terminal tails, or rarely elsewhere. Although histone sequences are well-conserved, especially where they undergo these PTMs, there are also histone paralogs called histone variants where PTM sites are often replaced with other amino acids (3). Similar to histones, these histone variants are also key players in regulating the chromatin structure and controlling diverse biological processes (4–7).

Plant genomes underwent several duplication and hybridisation events during their evolutionary timeline which contributed to the presence of diverse transposable elements (TEs) and repeats which are tightly regulated. Unlike the model plant *Arabidopsis*, such repeats and TEs are closely associated with genic regions in plants with larger genomes such as rice (8, 9). Histone variants play specific roles in transcriptional control of regions including repeats and TEs along with genic regions, and they mediate development of specific organs/tissues as well as stress responses (6). These histone variants are very diverse in their function; however conserved variants play specific roles across eukaryotes. The functions of these histone variants are based on their expression pattern, their unique PTMs that are different from the canonical forms, their structural properties, chaperone specificities, and genomic incorporation sites (7, 10–17).

Plants possess several histone variants including well conserved histone variants that are seen across eukaryotes as well as specialised family/clade-specific variants with either tissue-specific expression or stress-induced expression. In plants, well conserved histone variants of H3 include CENH3 and H3.3, while clade/genera-specific variants mainly consist of H3.15, H3.14 and H3.10. CENH3 is important for cell division and modifications in this variant led to haploid induction in different plants (18, 19). It is also known that parental CENH3 polymorphisms create epigenetically distinct centromeres to induce mating barrier in *Arabidopsis* (20). H3.3 variant is important in regulating gene expression by positively influencing active histone marks. Mutants of H3.3 display early flowering phenotype that is mostly attributed to its influence on the FLC locus (21). h3.3 knockout (ko) plants also show severe developmental defects, including smaller rosette leaves and siliques (22). The sperm cell specific H3.10 along with H2B.8 are important for compacting sperm nuclei, while H3.15 is important in regulating callus induction (23–25). Similarly, generally less-expressed but salt stress inducible H3.14 is important for mediating salt stress response in the roots of *Arabidopsis* (23–26). The role of *Oryza*-specific H4 variant in salt stress response is another example of the importance of genera-specific histone variants in plants (17). The mutants of all these variants have often compromised growth and development and in a few cases show abnormal phenotypes under specific stress conditions. Apart from these defects, mutants of histone variants such as H3.3 in *Arabidopsis* are lethal (27, 28).

H2A variants are conserved across eukaryotes, and among them are H2A.Z and H2A.X. Unlike H3 variants, the C terminal PTM modifications and sites of their incorporation in the genome, as determined by their specific chaperone proteins, dictate the functional outcome of these variants (29). H2A.Z variant is the most evolutionarily conserved H2A variant and it is seen to be mostly associated with the promoters and transcription start sites (TSS) of several stress responsive genes (30). Multiple key findings have systematically characterized functions of H2A.Z among plants. The minimal incorporation of H2A.Z observed in gene bodies is decisive enough to prevent or interfere with DNA methylation in gene bodies (31). It was also observed that there is a significant reduction in the levels of H2A.Z in the gene bodies of actively expressed genes that also have high DNA methylation (32, 33). It is not known if DNA methylation in promoters of genes is also influenced by H2A.Z. Dynamics of H2A.Z is mostly regulated by environmental stresses and this is mediated by the diverse chromatin remodelling complexes that act as its chaperones (34, 35). Evidence from *Arabidopsis* suggests that gene body occupancy of H2A.Z is repressive, while +1 nucleosome distribution of H2A.Z activates gene expression (36, 37). The repressive role of H2A.Z is mostly due to its property to create less accessible chromatin environment, reduced H3K4me3 and high levels of H3K27me3 marks (38). The incorporation of H2A.Z to nucleosome requires a SWI2/SNF2-RELATED1 (SWR1) complex containing PHOTOPERIOD-INDEPENDENT EARLY FLOWERING1 (PIE1) and ACTIN-RELATED PROTEIN 6 (ARP6) (39–41). Under heat stress, H2A.Z is removed from thermo-responsive and auxin-related genes to induce their expression and this is mediated by INOSITOL REQUIRING NUCLEOSOME REMODELING FACTOR (INO80) and PHYTOCHROME INTERACTING FACTOR 4 (PIF4) (42). In addition to this, specific PTMs in the H2A.Z, mostly monoubiquitination, contributes to repressive function of the variant (43). H2A.Z seems to be playing roles in development too, for example, it acts as an activator of floral repressor FLC and in agreement with this, knockdown of H2A.Z led to early flowering in *Arabidopsis* (31, 44). Among the other H2A variants, H2A.W is a plant-specific variant having a C-terminal SPKK motif which helps in compacting the chromatin and repressing TEs and repeats in heterochromatin regions (45, 46). H2A.W is incorporated by DECREASE IN DNA METHYLATION1 (DDM1) and is hence associated with repressive H3K9me2 marks (47). A modified H2A.W variant having SQEF motif, in addition to SPKK motif is known to regulate heterochromatin-specific DNA damage repair in *Arabidopsis* (29). H2A.W mutant in *Arabidopsis* did not show any developmental defects, however, these mutants had increased occupancy of H2A and H2A.X across TEs indicating a competition between these variants. The enrichment for H2A.X on centromeric repeats, replacing H2A.W in h2a.w ko was observed. In addition, the loss of H2A.W from centromeres led to higher levels of H1 to further reduce accessibility for DNA methyltransferases to induce DNA methylation (46, 48). Similar to H2A.W the linker histone H1 is also important for regulating the accessibility of heterochromatin regions (46, 49, 50).

Unlike plant-specific H2A.W, H2A.X is a well conserved H2A variant that is also a major regulator of DNA damage response and associated functions among eukaryotes (51, 52). H2A.X is distributed across the genome in both euchromatic regions, including gene bodies, and heterochromatic regions (45). It is known that FACILITATES CHROMATIN TRANSCRIPTION (FACT) complex is the chaperone for H2A.X and as expected, mutants of FACT complex reduced the levels of H2A.X incorporation (6, 53–55). FACT complex consists of SUPPRESSOR OF TY (SPT) 16 and STRUCTURE SPECIFIC RECOGNITION PROTEIN 1 (SSRP1) proteins and mutants of both these proteins exhibited severe developmental and reproductive defects in *Arabidopsis* (56–60). Upon DNA damage, the phosphorylation of the conserved serine residue in the SQEF motif of DNA-damage induced H2A.X by ATAXIA TELANGIECTASIA MUTATED (ATM) and ATAXIA TELANGIECTASIA AND RAD-3-RELATED (ATR) kinases are important for its function (61). Phospho-H2A.X signal is a very important mark for active DNA damage across organisms. MEDIATOR OF DNA DAMAGE CHECKPOINT 1 (MDC1) is a key DNA damage repair associated protein that interacts with phosphorylated tail of H2A.X using its BRCT domain, to recruit DNA damage repair (DDR) proteins to the DNA damage sites in animals (62, 63). Recent studies in *Arabidopsis* indicates that BRACT DOMAIN PROTEIN 4 (BCP4) is the homologue of MDC1 in plants and it interacts with phosphorylated H2A.X to recruit DDR proteins (64). Similarly, another BRCT domain containing protein gamma-H2A.X-INTERACTING PROTEIN (XIP) is also involved in regulating H2A.X dependent DNA damage repair in plants (65). Absence of H2A.X or the machinery to phosphorylate H2A.X disrupts DNA damage repair process in plants (66). In addition, the root length of *Arabidopsis* h2a.x ko plants are reduced upon DNA damage, indicating the influence of compromised DNA damage repair on development (55, 66).

Mutants of H2A.X in *Arabidopsis* show negligible root and silique development phenotypes (29, 55, 67). Probably due to the conserved role in DNA damage associated processes across eukaryotes, and since the mutants of H2A.X in *Arabidopsis* lack profound developmental defects, it has been generally assumed that function of H2A.X might be more specific to DNA damage response. Recent studies suggest that gamma-H2A.X is involved in regulating gene expression of DNA damage-associated genes in mammalian cell lines, where it has been identified that HIGH MOBILITY AT HOOK 2 (HMGA2) generated nicks in the DNA are the sites where gamma-H2A.X is incorporated by the action of the FACT complex to induce transcription (68). In plants, gamma-H2A.X interacts with the promoters of ABSCISIC ACID INSENSITIVE (ABI) genes to de-repress their expression (69). Probably connected to this, the *Arabidopsis* h2a.x ko plants show increased root growth and reduced cotyledon greening phenotype under ABA treatment (67, 69).

Here we show that, rice H2A.X contributes to rice development and growth, especially in specific stages and tissues, and this role of H2A.X is largely independent of its functions in DNA damage response. Using genetic, biochemical, and whole genome approaches, we show that H2A.X acts a suppressor of active H3K4me3 deposition across several development-associated genes. H2A.X also suppressed deposition of repressive H3K9me2 marks across genes and TEs by competing with another H2A variant, H2A.W. Unlike *Arabidopsis*, rice H2A.W deposition was not limited to peri-centromeric heterochromatin regions alone, largely because dispersed TEs and repeats in rice genome are often present near genic regions. Immunostaining using H2A.X and H2A.W specific antibodies and genomic analysis demonstrated competition between these two variants at specific genomic regions. Together, these results indicate that H2A.X plays crucial development-associated roles by maintaining specific chromatin states across hundreds of genes. It is likely that H2A.X plays similar roles across plants with larger genomes including crops.

## Results

### Tissue specifically expressed OsH2A.X is uniformly distributed in the nucleus

In order to understand the conservation of rice H2A.X, we aligned canonical H2A and its variants to identify amino acid variations that are observed in the variants (Fig. 1A). In *Arabidopsis*, H2A variants are encoded by 13 genes that include 4 copies of canonical H2A, 2 copies of H2A.X, 4 copies of H2A.Z and 3 copies of H2A.W respectively (Supplementary Fig. S1A). Among the 13 copies of H2A members in rice, 4 copies code for canonical H2A, 2 copies by H2A.X, H2A.Z with 3 copies and H2A.W is coded by 4 copies. Majority of the variations in H2A variants are either at the N-terminal tail or at the C-terminal, suggesting that these variants can alter the functions through amino acid variations alone. Both copies of H2A.X have several amino acid variations different from canonical as well as other H2A variants at both N and C terminal ends, while the conserved C terminal SQEF motif which is important for DNA damage signalling is conserved.

**Fig. 1:**
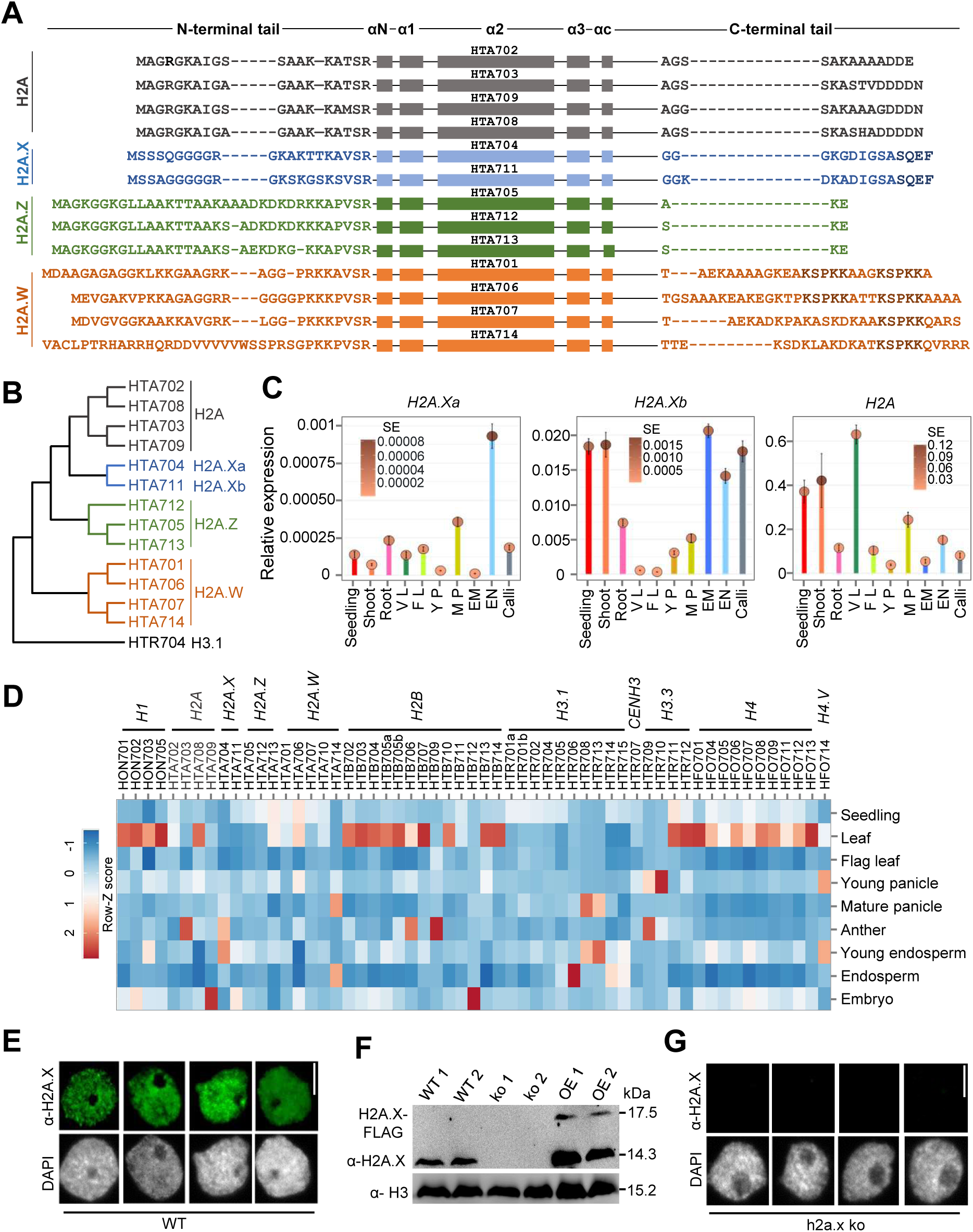
Tissue specifically expressed OsH2A.X is uniformly distributed in the nucleus. **A,** Aminoacid alignment of H2A variants from rice. Boxes depict different domains. Functionally conserved C-terminal residues are highlighted in bold. **B,** Phylogenetic tree of H2A variants from rice. Tree was generated using maximum likelihood method with 1000 bootstraps. **C,** Relative expression of H2A variants across tissues. *OsGAPDH* served as the internal control. Data represents means ± SE, n = 3. qRT-PCR experiments are performed twice with consistent results. V L – Vegetative Leaves, F L – Flag Leaves, Y P - Young Panicle, M P - Mature Panicle, EM – Embryo, EN - Endosperm. **D,** Heatmap depicting tissue-specific expression of histone variants from rice. Datasets used are-seedling (GSE229604), leaf (GSE229604), flag leaf (GSE111472), young panicle (GSE180457), mature panicle (GSE107903), anther (GSE180457), young endosperm (15 days after pollination-DAP), mature endosperm (25 DAP) and embryo (GSE229961). Row Z-score was plotted. **E,** Immunofluorescence (IFL) images of PB-1 nuclei probed with α-H2A.X antibody. **F,** Western blot with α-H2A.X antibody among rice lines. H3 is used as the loading control. **G,** IFL images of rice nuclei in h2a.x ko. Scale bar in **E** and **F** is 5 µm.

Surprisingly, the well conserved SPKK motif of H2A.W, important for chromatin compaction in *Arabidopsis*, has duplicated in two of the 4 copies of rice H2A.W variants (Fig. 1A). The conserved lysine residues in the N-terminal end of canonical H2A and H2A.Z, that are important for transcription control are absent in rice H2A.X variants (70, 71). H2A.X is phylogenetically closer to canonical H2A compared to other H2A variants such as H2A.W. (Fig. 1B). H2A variants in rice show distinct tissue specific expression pattern similar to *Arabidopsis*. (Fig. 1C and Supplementary Fig. S1A). Among the 2 copies of H2A.X (named a and b), *H2A.Xa* expresses highly in endosperm, whereas, *H2A.Xb* expresses highly in embryo, young endosperm, and seedling tissues (Fig. 1C, D). Such a tissue specific expression was also observed among H2A.Z and H2A.W variants in rice (Supplementary Fig. S1B). Tissue specific expression of most histone variants is also observed in rice (Fig. 1D). We generated specific antibodies for rice H2A.X to understand native functions of the variant without a transgenic approach. This antibody effectively detected both the H2A.X variants in rice and also did not cross-react with rice H2A or other H2A variants that were expressed in *E. coli* (Supplementary Fig. S1C). Further we checked the H2A.X protein levels and found that it is consistently high in seedling and reproductive tissues that were not subjected to any stress (Supplementary Fig. S1D). Immunostaining of nuclei derived from seedling tissues indicated uniform distribution of H2A.X throughout the nuclei, excluding nucleoli, that indicates genome-wide distribution of H2A.X in the absence of DNA damage (Fig. 1E and Supplementary Fig. S1E). This genome-wide distribution of H2A.X observed here is different from the nuclear distribution pattern of *Arabidopsis* gamma H2A.X in DNA damaged tissues, where multiple studies report clear nuclear speckles correlating with the sites of DNA damage (64, 65). All these indicate that unmodified plant H2A.X might have functions beyond DNA damage response as proposed previously (5). Among animals and plants, such a possibility is not fully explored, even though some tissue-specific or condition/treatment-specific phenotypes were documented in h2a.x mutants (55, 67, 69).

### Mis-expression of H2A.X led to development-associated phenotypes

In order to understand the function of H2A.X beyond DNA damage response, we generated transgenic lines mis-expressing H2A.X in *indica* rice (*Oryza sativa indica* Pusa basmati-1 (henceforth PB-1)) lines. The complete ko of H2A.X was achieved by targeting both members of H2A.X (*H2A.Xa* and *H2A.Xb*) using two different gRNAs in a single gRNA cassette (Supplementary Fig. S2A, Methods). Nine plants were obtained using *Agrobacterium*-mediated transformation (Supplementary Fig. S2B-G). The transgene integration was verified using junction fragment Southern analysis that indicated single copy insertions of T-DNA in two independent ko lines named ko 1 and ko 2 (Supplementary Fig. S2C). Further, T-DNA free edited, homozygous lines were selected by segregation and verified by PCR (Supplementary Fig. S2D). Among the two ko lines selected, ko 1 had a 1-nt insertion in exon2 of *H2A.Xa*, while *H2A.Xb* targeting resulted in a 1-nt deletion in exon1, both mutants leading to missense mutations. The ko 2 plant also induced a 1-nt insertion at *H2A.Xa* and had a 1-nt deletion in exon1 of *H2A.Xb*, again leading to missense mutation (Supplementary Fig. S2E). The ko of H2A.X at the protein levels was verified through western analysis using H2A.X-specific antibody (Fig. 1F and Supplementary Fig. S2F). In agreement with the absence of H2A.X, ko 1 did not show positive signal for H2A.X upon nuclear immunostaining (Fig. 1E and Supplementary Fig. FS2E). Since H2A.X gets upregulated under DNA damage, we generated constitutively expressing (OE) *H2A.Xa* lines with 3xFLAG tag as a control line that somewhat mimics DNA damage response (Supplementary Fig. S3A and S3G). Among the 5 transgenic lines obtained, OE 1 and OE 2, plants having T-DNA insertions as assayed through junction fragment Southern analysis were selected for further analysis (Supplementary Fig. S3B, C). Both transgenic lines expressed H2A.X at very high levels, as observed though RT-qPCR as well as western analysis (Fig. 1F and Supplementary Fig. S3D-F).

Both the ko and OE lines had several developmental abnormalities. The ko line showed longer roots while the overall seedling length was not significantly altered (Fig. 2A-D). On the other hand, the OE lines displayed stressed phenotypes, having shorter roots, but with longer lateral roots (Fig. 2A-D and Supplementary Fig. S4C). The OE plants also had high levels of anthocyanin, particularly in the stems and awns of seeds (Fig. 2E-G). Apart from this, the ko seedlings had increased chlorophyll content (Fig. 2H). The ko plants had reduced height, increased tiller numbers, and panicles, as well as delayed heading date (Fig. 2J-L and Supplementary Fig. S4D). Both ko and OE plants had significant reproductive defects with partially sterile panicles having significantly lesser number of seeds in each panicle (Fig. 2I and 2M). The seeds were smaller and the weight of the seeds was also significantly reduced in ko lines (Fig. 2M and Supplementary Fig. S4E). Both in ko and OE lines, the percentage of viable pollens were significantly reduced when compared to WT (Fig. 2N, O). The OE plants also showed partial male sterility whereas pollens were much more deformed in ko lines (Fig. 2Q, Supplementary Fig. S4L). The ko plants showed dehydrated stigmas, while anthocyanin accumulation was observed in the stigmas of OE plants (Fig. 2P). All these phenotypes conclusively indicate that both ko or constitutive expression of H2A.X led to severe phenotypic abnormalities.

**Fig. 2:**
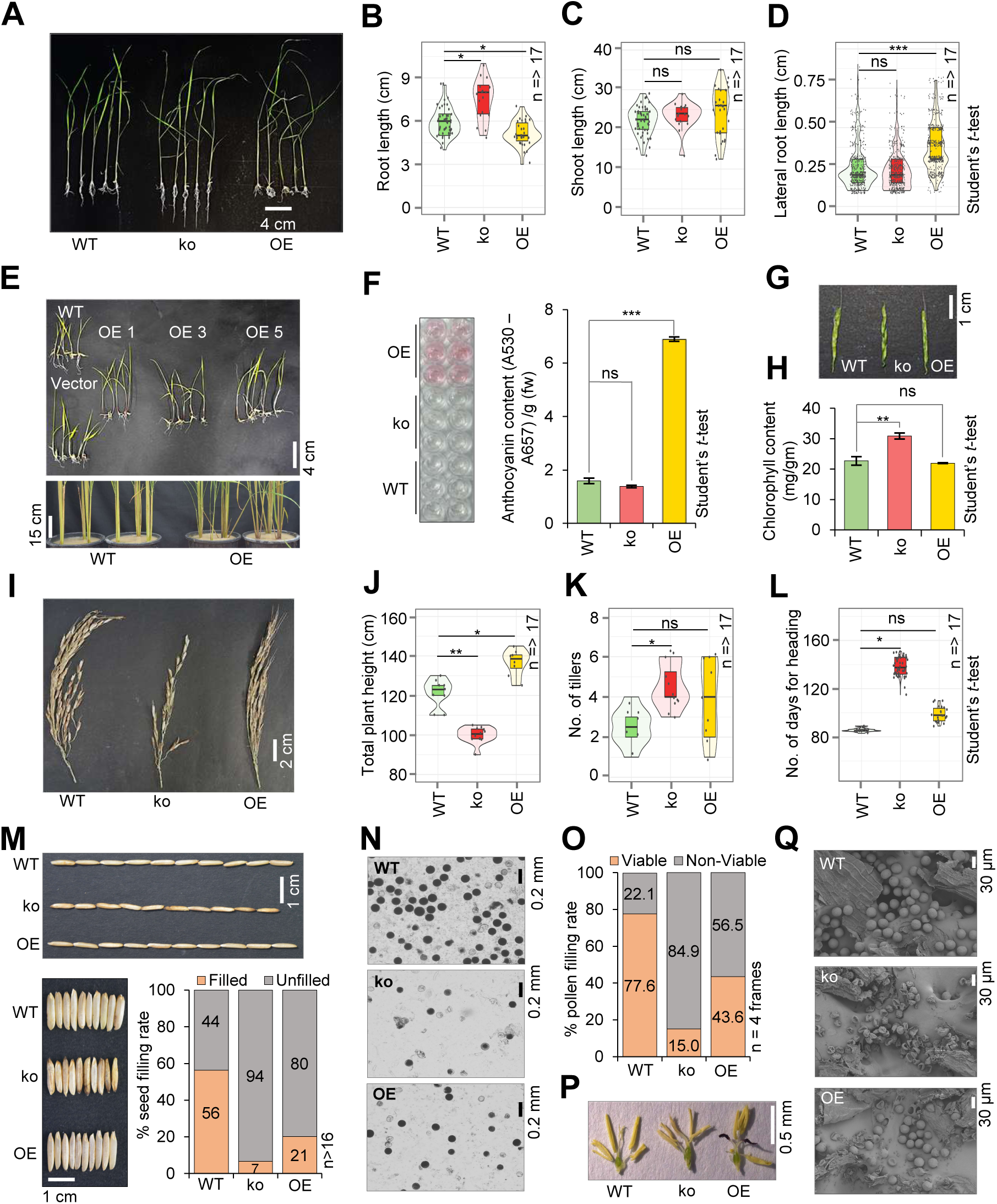
Phenotypes of H2A.X mis-expression lines. **A,** Morphology of 14-day old seedlings of H2A.X mis-expression lines. Box plots showing root length **B,** shoot length **C,** and lateral root length **D,** among lines. **E,** Images showing anthocyanin accumulation. **F,** Anthocyanin pigment obtained in the assay (left). Quantification of anthocyanin accumulation in ko and OE seedlings (right). **G,** Image showing accumulation of anthocyanin in seed awns. **H,** Box plot showing level of chlorophyll content in H2A.X mis-expression lines. **I,** Image showing panicle morphology of H2A.X mis-expression lines. Box plots displaying plant height **J,** tiller number **K,** and heading date **L,** of H2A.X transgenic lines. **M,** Image showing seed morphology in H2A.X transgenic lines. Barplot depicts the percentage of filled seeds. **N,** Representative images showing iodine-stained pollen grains of dehisced anther. **O,** Barplot showing percentage of viable and non-viable pollen after iodine staining. **P,** Phenotypes of reproductive structures. **Q,** Representative scanning electron microscope (SEM) images of pollen grains. Two-tailed Student’s *t*-test was used for statistical comparison in **B, C, D, F, H, J, K** and **L**. *p*-value (*) < 0.05, (**) < 0.01, (***) < 0.001, (ns) non-significant.

Since h2a.x ko plants set very few viable seeds, we also generated H2A.X knockdown (kd) using artificial microRNAs (amiR) designed to target both isoforms in a single construct (Supplementary Fig. S3H). Optimal amiRs were selected based on the previously identified position specificity (72, 73). We obtained 8 kd lines and the expression of amiR was confirmed using sRNA northern analysis (Supplementary Fig. S3I-K). Among these plants we selected kd 1, which showed around 50% reduction in the accumulation of both isoforms of H2A.X for further analysis (Supplementary Fig. S3J-L). Similar to ko lines, kd lines also had abnormal growth and development-associated phenotypes, such as increased seedling root length and delayed flowering (Supplementary Fig. S4A, B and F). Similar to ko, the kd lines also showed increased panicle number, reduced panicle filling rate, reduced seed number, and seed weight (Supplementary Fig. S4G-K). As expected, the kd phenotypes were less severe when compared to ko lines. Together, these results suggest a development-associated role of rice H2A.X beyond DNA damage repair.

### H2A.X mutants display altered expression of development and stress-related genes

In order to understand the molecular basis for the altered phenotypes in H2A.X mis-expression lines, we performed RNA sequencing (RNA-seq) of 14-day old seedlings and obtained a mapping efficiency of 95% to the rice genome (IRGSP 1.0) (Supplementary Table S1). We obtained 367 upregulated and 130 downregulated genes in ko as well as 612 upregulated and 353 downregulated genes in OE lines with a stringent log_2_ 1.5-fold cutoff (Fig. 3A, Supplementary Fig. S5A, Supplementary Data S1 and S2). In agreement with root growth and chlorophyll related phenotypes observed in the transgenic lines, several key genes in these pathways were altered in ko plants (Fig. 3A). GO analysis of differentially expressed genes (DEGs) in ko indicated enrichment of photosynthetic and pigment metabolism related genes (Fig. 3B). On the other hand, GO analysis of OE upregulated genes indicated enrichments in phenylpropanoid pathway genes, while downregulated genes in OE were associated with photosynthesis and related processes (Supplementary Fig. S5B, C). The upregulated genes common in ko and kd showed enrichment for important plant metabolic processes and stress response (Supplementary Fig. S5D and S6A, Supplementary Data S3). These results confirm that H2A.X is crucial for regulating genes involved in seedling development, especially root growth and metabolism. A principal component analysis (PCA) with ko DEGs indicated that the set of genes altered in h2a.x ko are very different from any other stress including DNA damage stress (Fig. 3C) (74). In agreement with the root-associated phenotypes, expression of genes involved in root development were altered in H2A.X mis-expression lines. The genes positively regulating root growth were highly expressed in ko, while those involved in negative regulation showed high expression in OE lines (Fig. 3D, Supplementary Fig. S5F and Supplementary Data S4). Increased chlorophyll content in ko plants was associated with increased expression of photosynthesis associated genes (Fig. 3E and Supplementary Fig. S5F). Similarly, expression of genes associated with pigmentation were highly expressed in OE plants as expected (Fig. 3F and Supplementary Fig. S5F).

**Fig. 3:**
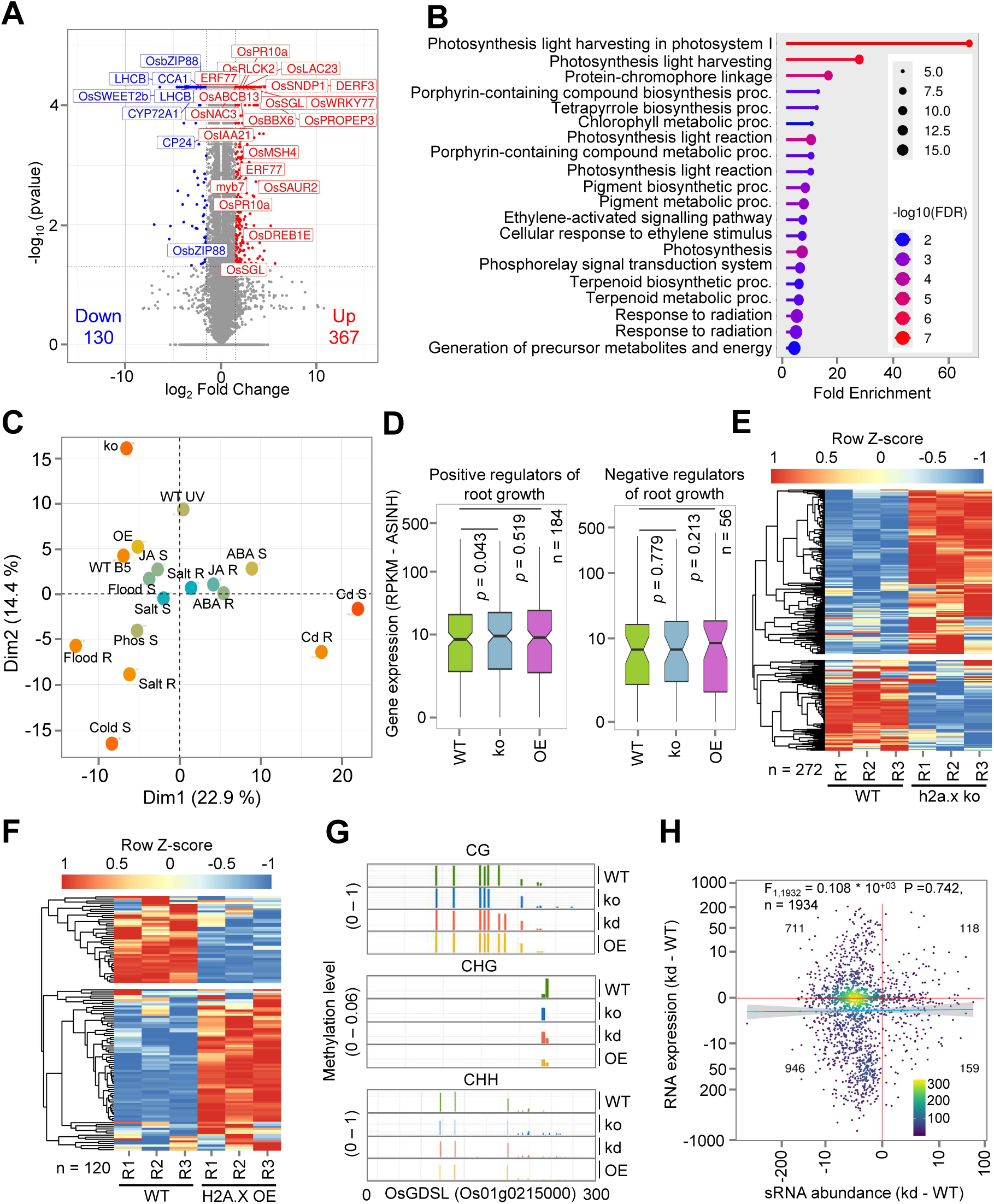
H2A.X mutants display altered expression of key genes. **A,** Volcano plot representing all DEGs in h2a.x ko. **B,** GO enrichment categories of ko DEGs. FDR-False discovery rate. **C,** PCA analysis of ko DEGs with different stress datasets. (R – root, S – shoot, ABA – abscisic acid, JA – jasmonic acid, Cd – cadmium, B5 - bleomycin) **D,** Boxplot showing expression of root growth regulators. **E,** Heatmap showing expression of photosynthesis associated genes. **F,** Heatmap showing expression of pigmentation associated genes. Row Z score is used to plot the heatmaps in **E** and **F**. **G,** Diagram showing DNA methylation in different contexts in the promoter of a mis-expressed gene *OsGDSL*. **H,** Correlation plot showing expression of sRNAs lost in h2a.x kd and mRNA expression of adjacent genes (RPKM) in h2a.x kd. Two-sided Wilcoxon test (*p* < 0.01 was considered significant) was used for **D**.

Previous studies (68, 75) indicated that h2a.x ko in *Arabidopsis* and kd in human cell lines led to DNA hypomethylation. H2A.X was also found to be essential for regulating DNA methylation specifically in endosperm tissues (75). In order to understand whether the gene expression changes observed in the rice h2a.x ko mutants were due to altered DNA methylation, we performed targeted gene-specific bisulfite PCRs in the promoters of several mis-expressed genes including *SMALL AUXIN-UP RNA 8* (*OsSAUR8*) and *GDSL ESTERASE/LIPASE PROTEIN 4* (*OsGDSL/OsGELP4*) (Fig. 3G and Supplementary Fig. S6C-G). There was clearly an absence of correlation with promoter DNA methylation levels and gene expression in transgenic lines (Fig. 3G and Supplementary Fig. S6C-G). Since promoter small (s)RNA levels and sRNA-directed DNA methylation (RdDM) are involved in regulating DNA methylation, we performed genome-wide sRNA sequencing in 14-day old seedlings derived from wild type, h2a.x kd and OE lines. This analysis clearly indicated that levels of sRNAs and DNA methylation did not correlate with gene expression changes in H2A.X mis-expression lines (Fig. 3H, Supplementary Fig. S5G and 6B and Supplementary Data S5). These results conclusively indicate that sRNA-mediated functions did not directly contribute to gene expression changes and associated plant phenotypes among H2A.X mis-expression lines, albeit in normal unstressed conditions.

### H2A.X suppresses deposition of active H3K4me3 marks over thousands of gene promoters

Since histone variants alter deposition and maintenance of histone marks over the genomic regions that they decorate, we performed chromatin immunoprecipitation followed by sequencing (ChIP-seq) for active and inactive histone marks (H3K4me3 and H3K9me2) in 14-day old seedlings derived from WT and h2a.x ko lines. As observed previously in rice (76), H3K4me3 mark was predominantly associated with promoters (Supplementary Fig. S7B). There were around 3700 additional peaks of H3K4me3, approximately 15% increase over 24109 normal peaks, in h2a.x ko plants based on a stringent 2-fold enrichment cut-off (Supplementary Fig. S7C and Supplementary Data S6). Apart from the increase in number of peaks there was an overall increase in H3K4me3 enrichment in h2a.x ko over the H3K4me3 marked regions shared with WT (Fig. 4A).

**Fig. 4:**
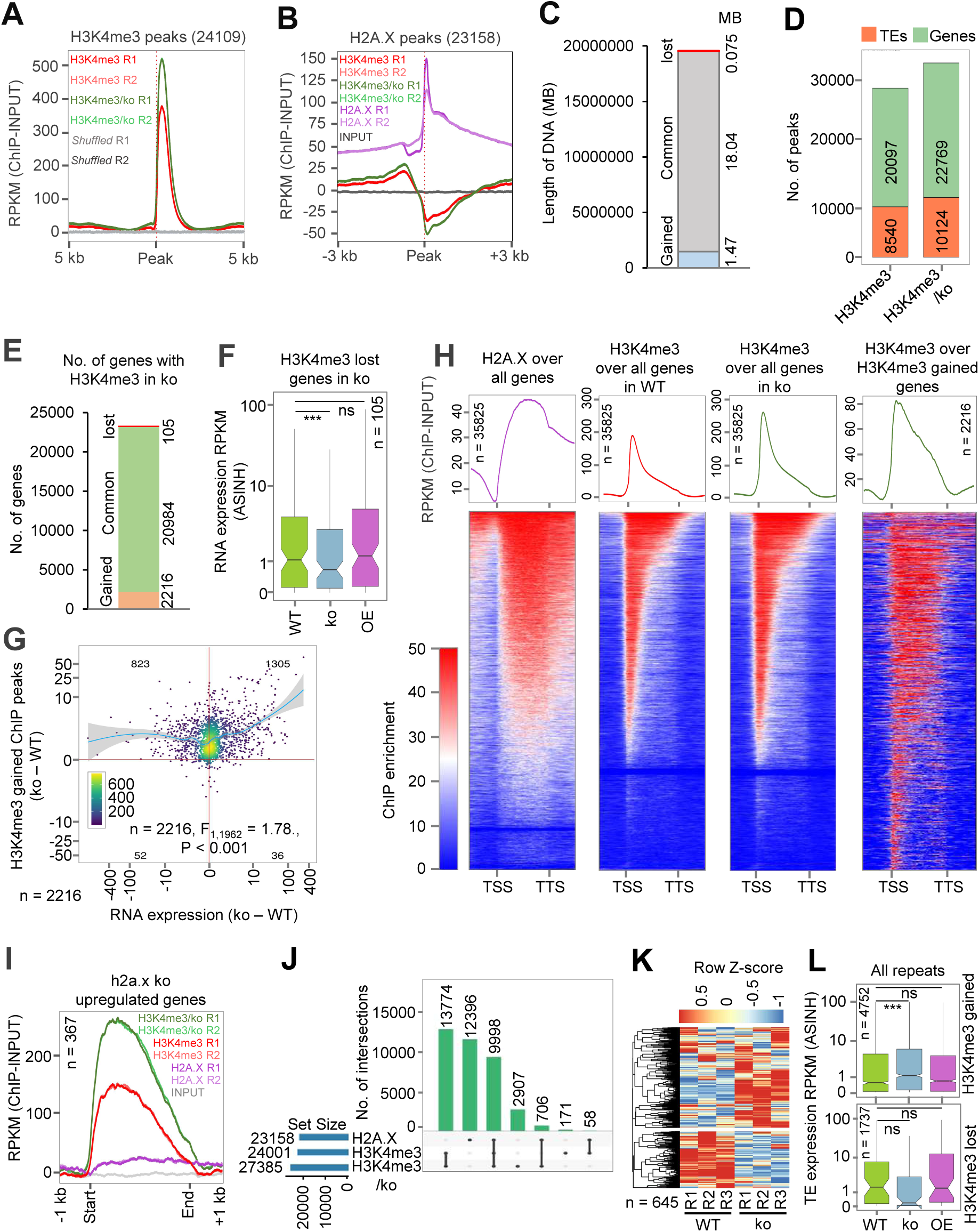
H2A.X suppresses deposition of active H3K4me3 marks over genes. **A,** Metagene plot showing enrichment of H3K4me3 in WT and h2a.x ko. **B,** Metagene plot showing enrichment of H2A.X binding and H3K4me3 mark over H2A.X peaks in WT and h2a.x ko. **C,** Barplot displaying length of DNA occupied by H3K4me3 in WT and h2a.x ko. **D,** Barplot showing number of H3K4me3 peaks over genic and TE features. **E,** Barplot showing number of gained and lost genes associated with H3K4me3 peaks in ko. **F,** Boxplot displaying expression of genes upon loss of H3K4me3 mark in ko. **G,** Correlation plot of ChIP signal and RNA expression from H3K4me3 gained genes. **H,** Heatmap showing the enrichment of H2A.X and H3K4me3 over all genes and H3K4me3 gained genes in ko. **I,** Metagene plot showing enrichment of H3K4me3 and H2A.X over upregulated genes in ko. **J,** Upset plot depicting number of peaks common between H2A.X (2 kb extended) and H3K4me3 marks. **K,** Heatmap displaying the expression of H2A.X bound genes that gained H3K4me3 in ko. **L,** Boxplot displaying expression of TEs that gained and lost H3K4me3 peaks in ko. For boxplots, two-sided Wilcoxon test (*p* < 0.01 was considered significant) in **F** and **L**.

In order to explore if the additional as well as increased peaks for H3K4me3 correlated with the H2A.X binding regions, we performed an H2A.X ChIP-seq experiment in WT plants using native antibody raised against rice H2A.X. There were around 23158 H2A.X peaks in seedling tissues with a stringent cut-off of 2-fold enrichment (Supplementary Data S7). These regions largely overlapped with promoters (60.9%) and distal intergenic regions (28.54%) (Supplementary Table S4). The overall distribution of H2A.X in rice matched the genome wide distribution in *Arabidopsis* except an unusually high binding to 3’-UTR in rice (3.48% compared to 0.28% in *Arabidopsis*) (7) (Supplementary Table S4). The peak width of H2A.X was broader with a median peak width of around 900 bp, while that of H3K4me3 was 704 bp in WT and 780 in h2a.x ko (Supplementary Fig. S7A). Surprisingly, the peak boundaries of H2A.X peaks exhibited enrichment of H3K4me3 marks and this mark showed even more enrichment in h2a.x ko (Fig. 4B). The increased H3K4me3 marks spread over to a total of 1.4 MB genomic region when compared to wild type levels, while its loss was only over 0.075 MB genomic regions in h2a.x ko (Fig. 4C). There was an increase of H3K4me3 marks over several TEs (1363), while the increase was substantial over 2216 protein-coding genes (Fig. 4D, E and Supplementary Fig. S7D, E).

The distribution of H2A.X peaks in genes was mostly in the gene body while exhibiting a sharp reduction in the TSS sites. On the other hand, H3K4me3 marks were highly enriched in the TSS with a sharp reduction in gene bodies (Fig. 4H). This opposite pattern of enrichment was even more specific in H2A.X bound genes. Most of the H3K4me3 was more strictly in the TSS and 3’UTR in H2A.X bound genes when compared to all genes where there is a more dispersed distribution of H3K4me3 over the TSS and gene body. Surprisingly in ko, there was an overall increase in H3K4me3 levels in TSS spreading to the gene bodies (Supplementary Fig. S7G). Strikingly, the genes that gained H3K4me3 in h2a.x ko had a distribution of H3K4me3 over their gene bodies without any strict bias towards the TSS regions (Fig. 4H). These results suggest that H2A.X restricts H3K4me3 over both TSS and gene bodies. As expected of increased active H3K4me3 marks, the genes that gained this mark displayed increased expression in h2a.x ko and were specifically enriched for development-associated biological processes (Fig. 4G and Supplementary Fig. S7F and S7H). Similarly, 115 genes that lost H3K4me3 showed a clear reduction in expression in h2a.x ko (Fig. 4E, F). These results indicate that H2A.X acts as a suppressor of H3K4me3 marks over protein coding genes.

In order to study if the loss of H2A.X and H3K4me3 neo-deposition led to gene expression changes, we checked H3K4me3 enrichment over DEGs obtained from h2a.x ko plants and found increased H3K4me3 in the upregulated genes (Fig. 4I and Supplementary Fig. S7I). On the other hand, the downregulated genes in h2a.x ko did not show any substantial increase of H3K4me3 (Supplementary Fig. S7J). Since H2A.X displayed co-operative binding near to H3K4me3 peaks, we overlapped peaks of H2A.X and H3K4me3 marks and found noticeable overlap of these features across 10000 sites when the H2A.X peaks overlapping 2kb regions were considered (Fig. 4J). From this analysis we obtained 645 genes which exhibited gain of H3K4me3 and significant H2A.X binding. As expected, these genes showed pronounced high expression in h2a.x ko plants (Fig. 4K). Among the genes which showed increased levels of H3K4me3 marks several genes were associated with development including *CYCLASE-LIKE 4* (*OsCYL4*) and *PHOSPHATE-INDUCED PROTEIN 1* (*OsPhi-1*) related gene, *ROOT GROWTH ASSOCIATED 1 CYS-PEROXIREDOXIN B* (*Os1-CysPrxB*) and *ZINC FINGER PROTEIN 350* (*OsZFP350*) and photosynthesis related *HYDROXYPYRUVATE REDUCTASE 1* (*OsHPR1*) gene (Supplementary Fig. S8A and S8D). Apart from this increased expression of specific genes, there were also stretches of genic regions that gained H3K4me3 and most of the genes in these regions displayed increased expression in ko (Supplementary Fig. S8B). All these results conclusively indicate that upregulated genes in h2a.x ko were due to increased H3K4me3 levels, and H2A.X suppresses the deposition of H3K4me3 mark over development-associated genes.

Although a genome-wide increase in H3K4me3 across many TEs and repeat regions was not observed (Fig. 4D), there were about 1363 TEs that gained H3K4me3 marks in ko (Supplementary Fig. S7E). This suggests that similar to genic regions, there was increased levels of H3K4me3 marks in the TEs also (Supplementary Fig. S9A). This increase in active H3K4me3 levels was sufficient to activate RNA expression of the TEs, predominantly from LTR elements and all other repeats (Fig. 4L and Supplementary Fig. S9B and S9D). This reinforces the observation in genic regions that H2A.X is a genome-wide suppressor of active marks. Transcription of TEs and expression from genomic regions did not lead to their RNA silencing, the hallmark of which is the production of 24 nt sRNAs in h2a.x kd or OE lines (Supplementary Fig. S9C). The fate and activities of these TE-encoded RNAs in h2a.x ko lines is an important avenue for future research.

The increase in active mark H3K4me3 might have resulted from increased expression of writers of this mark or due to the reduced expression of its erasers. In order to understand if that is the case, we checked the expression of these categories of histone modifiers in a candidate-gene specific manner. Among the writers which includes several histone methyltransferases under Trithorax family, there was a negligible change in the expression in h2a.x ko, while erasers of H3K4me3 including several Jumonji and *LSD1-LIKE ZINC FINGER PROTEIN* (*Lsd1*) showed marked reduction in the expression (Supplementary Fig. S8C). It is possible that the specific deposition pattern of H2A.X and H3K4me3 mark over genes and the reduced expression of several histone demethylases could have led to increased H3K4me3 and altered gene expression in h2a.x ko plants.

### Loss of H2A.X led to spreading of H3K9me2 across both genic and TE regions

In order to understand if H2A.X can influence other H2A variants and associated histone marks, we performed a ChIP-seq using H3K9me2 mark-specific antibody in 14-day old seedlings of WT and h2a.x ko plants. Using a stringent cut-off, we identified around 8585 peaks in WT conditions overlapping as expected with TEs and repeats (Supplementary Fig. S10A and Supplementary Data S6). The peak width of H3K9me2 peaks were around 350 bp. In h2a.x ko, the number of H3K9me2 peaks increased to 11593 indicating about a staggering 25% increase (Supplementary Fig. S10A-D). When we considered the length of DNA bound by the H3K9me2 mark there was negligible decrease in ko (0.65 MB region, 1775 peaks), whereas gained genomic region was 57% more than WT (4783 peaks, 1.9 MB) (Fig. 5A). Most of the H3K9me2 gained regions were enriched with TEs, while very little increase was observed over genes (Fig. 5B and Supplementary Fig. S10E, F). Among the 4752 TEs that overlapped with increased H3K9me2 marks in h2a.x ko, there were categories from both type I and type II TE members, such as Gypsy, Copia and En-spm (Supplementary Fig. S10G). Surprisingly these TEs showed negligible expression in WT and the change in the expression of these TEs or sRNAs from these TEs was not observed in h2a.x ko, indicating that the increased H3K9me2 did not alter either TE expression or sRNA production from these sites (Supplementary Fig. S10H-K). These results indicate that H2A.X somehow suppresses the distribution of H3K9me2 marks over TEs but is not involved in regulating the TE expression or in sRNA production. It is likely that these TEs were part of the constitutive chromatin that did not require RdDM or other regulation to keep this region suppressed.

**Fig. 5:**
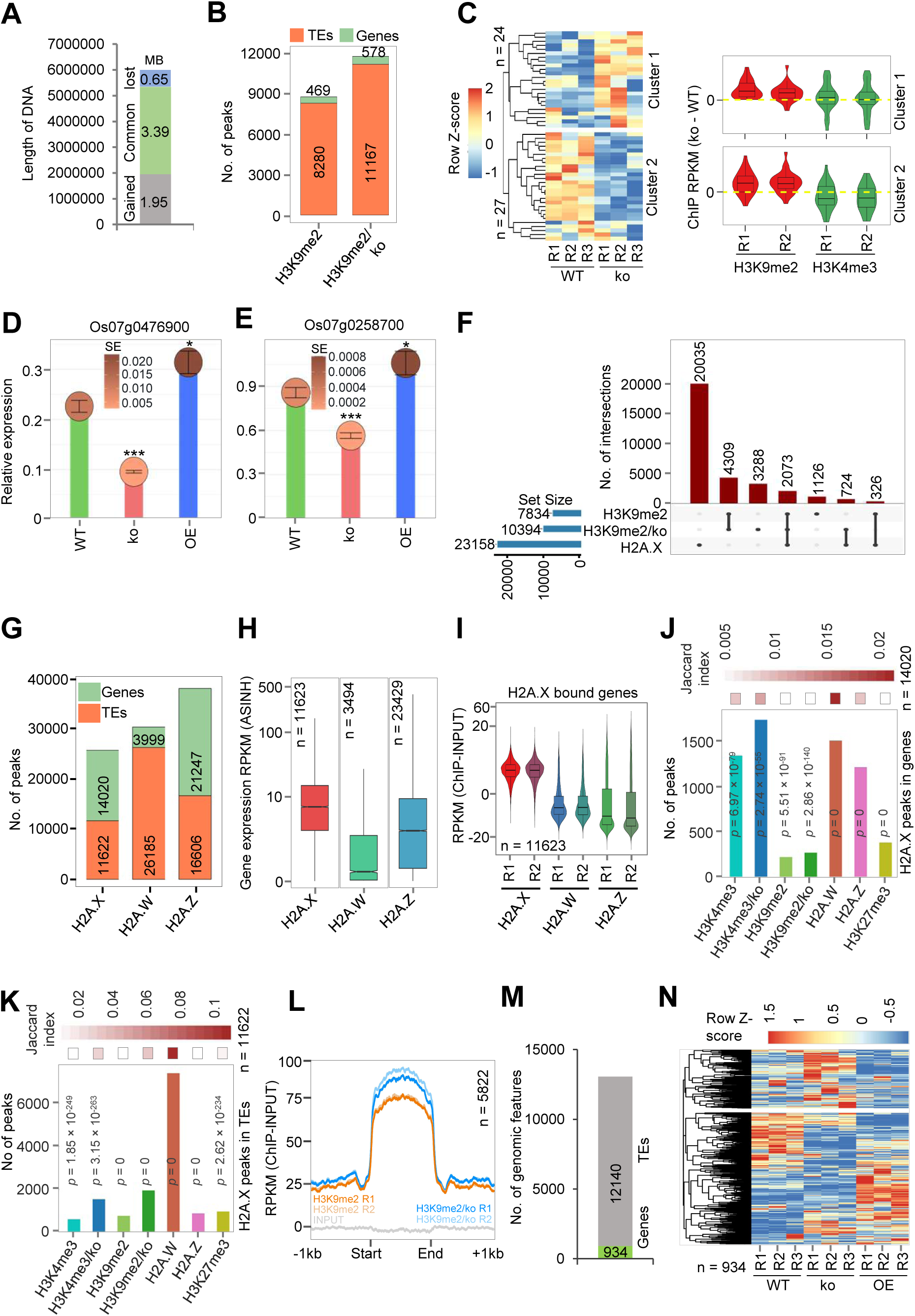
Loss of H2A.X led to spreading of H3K9me2. **A,** Barplot displaying length of DNA occupied by H3K9me2 in WT and ko. **B,** Barplot showing number of H3K4me3 peaks over genic and TE features across lines. **C,** Heatmap showing expression of H3K9me2 gained genes and boxplot showing the differential enrichment of H3K9me2 and H3K4me3 marks over these genes in ko. qRT-PCR showing the expression of representative H3K9me2 gained genes, Os07g0476900 (*OsCDSP32*) **D,** and Os07g0258700 (*C2-BTB-type E3 ubiquitin ligase 1*) **E,.** *OsGAPDH* served as internal control and two-tailed Student’s *t*-test was used for statistical comparison. Data represents means ± SE, n = 3. qRT-PCR experiments are performed twice with consistent results. **F,** Upset plot depicting number of peaks common between H2A.X (2 kb extended) and H3K9me2 marks in WT and ko. **G,** Barplot showing number of H2A.X, H2A.W and H2A.Z peaks over genic and TE features. **H,** Boxplot showing expression status of H2A variant bound genes. **I,** ChIP enrichment of H2A variants over H2A.X bound genes. **J,** Barplot showing number of overlaps of H2A.X peaks intersected with genes (14020) with peaks of histone variants and histone marks. **K,** Bar plot showing number of overlaps of H2A.X peaks intersected to TEs (11622) with peaks of histone variants and histone marks. **L,** Metagene plot showing enrichment of H3K9me2 mark over shared peaks of H2A.X and H2A.W in WT and ko. **M,** Boxplot showing number of genes and TEs in shared peaks of H2A.X and H2A.W. **N,** Heatmap showing expression of genes in shared regions in ko. Significance of overlap was tested using hyper-geometric test and *p*-values are mentioned. Jaccard index (shown as heatmap) represents the strength of overlap.

Interestingly, H3K9me2 marks also spread to genic regions encompassing approximately 200 genes when genes and their 1 kb promoter was considered. Among the 51 selected genes having expression in the tissues checked and having H3K9me2 peaks over gene body or 1 kb promoter regions, 27 genes showed reduced expression in h2a.x ko plants as expected (Fig. 5C, Supplementary Fig. S11A). The other 24 genes that had increased peaks of H3K9me2, also had increased levels of active H3K4me3 marks in the regulatory regions, indicating that repressive mark was not sufficient to compete with the active marks (Supplementary Fig. S11B). Among the 27 genes that showed reduced expression in ko, there were development-associated genes including Os07g0476900 (*CDSP32*, a plant thioredoxin involved in regulating photosynthesis) and an E3 ubiquitin ligase gene, Os07g0258700 (*OsC2-BTB1*). We performed qPCR to further confirm the repressive role of H3K9me2 in these genes and found that the genes showed reduced expression in ko while they were slightly upregulated in H2A.X OE (Fig. 5D, E). These results further confirm the regulatory role of H2A.X in a set of protein coding genes by altering repressive H3K9me2 mark. Apart from this, there were several genes that lost H3K9me2 in the ko and they showed increased expression (Supplementary Fig. S11C). Together, these results indicate that altered histone marks, both active and repressive, contributed to the gene expression changes in h2a.x ko plants, indicating that H2A.X dictates these important changes on a genome-scale.

### H2A.X inhibits incorporation of histone variant H2A.W to restrict H3K9me2 spreading

We observed overlapping distribution of H3K9me2 with H2A.X binding sites (Supplementary Fig. S10L), among at least 2073 sites when 2 kb extended H2A.X peaks were considered. However, there were also 724 unique regions with newer peaks of H3K9me2 in h2a.x ko (Fig. 5F). We hypothesized that this increased H3K9me2 spreading might be due to increased activity of writers of this mark or altered incorporation of other H2A variants. It is well known that H2A variants, especially H2A.W has the potential to influence H3K9me2 distribution (45). In order to further study if the gain of H3K9me2 is associated with H2A.W we performed ChIP-seq for histone variant H2A.W using the custom generated H2A.W antibody that did not cross-react with other variants (Supplementary Fig. S1C). We also analyzed the publicly available H2A.Z ChIP-seq datasets from rice seedlings (Supplementary Fig. S12A). We obtained around 37853 peaks of H2A.Z and most of them were associated with gene promoters (Supplementary Data S7). On the other hand, H2A.X peaks were mostly associated with promoters and distal intergenic regions (Supplementary Fig. S12A). We also obtained around 30197 peaks of H2A.W in WT and as shown in *Arabidopsis,* most of these peaks were associated with distal intergenic regions (61.77%). In addition to high levels of H2A.W over heterochromatin regions, H2A.W peaks were also enriched over promoters in rice (35.48%). Similar to the enrichment of these marks over specific chromatin regions, they displayed unique enrichment with either TEs or genes (Fig. 5G). Around 14020 peaks of H2A.X overlapped with 11623 genes, while 11622 peaks overlapped with 15013 TEs. H2A.Z peaks were mostly seen associated with genes, with around 21247 peaks overlapping with 23429 genes, while 16606 peaks overlapping with 26140 TEs. As expected, most of the H2A.W peaks, i.e., around 26185 peaks, overlapped with 47883 TEs, while, fewer peaks of around 3999 overlapped with 3494 genes (Fig. 5G, Supplementary Fig. S13D and Supplementary Data S8).

Unlike *Arabidopsis* where H2A.W preferentially binds to centromeric and peri-centromeric regions having TEs, rice H2A.W binding was spread-out, without a strict preference for peri-centromeric regions (Supplementary Fig. S12B and S13A, B). This might be due to the difference in the nature of heterochromatin distribution between these two model systems. While in *Arabidopsis*, H3K9me2 and H2A.W were strictly restricted to heterochromatic regions, such a distribution is not seen in rice, where heterochromatic marks are spread out to regions including genic regions in most of the chromosomes (Supplementary Fig. S12B) (77–79). In agreement with this, heterochromatin specific H2A.W was enriched in binding to repeats and in some gene-rich regions including promoters (Supplementary Fig. S12A and Supplementary Fig. S13A, B).

In order to understand the regulation of gene expression by these variants we checked the expression levels of genes bound by these variants and observed specific expression patterns for a given H2A variant. H2A.X bound genes were well-expressed without enrichment of either H2A.W or H2A.Z (Fig. 5H, I) and this pattern matched previous observations (80). On the other hand, H2A.W bound genes were low-expressed, while H2A.Z bound genes exhibited varied gene expression mostly in the range between the levels observed in H2A.X and H2A.W bound genes (Fig. 5H). In order to explore if the histone variants have shared genes where they bind to influence histone marks, the enrichment of H2A variants and histone marks with respect to H2A.X binding was analyzed. The peaks of H2A.X overlapping with genes and TEs were overlapped with the peaks of histone variants and histone marks. Both H2A.W and H2A.Z peaks were seen overlapping with H2A.X peaks across genes while H2A.X peaks overlapping with TEs also overlapped with H2A.W. Surprisingly, levels of both H3K4me3 and H3K9me2 marks over TEs were increased in h2a.x ko (Fig. 5J, K).

Further, we found H2A.X and H2A.W bound peaks overlapped, in around 5822 regions, while H2A.W overlapped with H2A.Z in 2223 genomic regions (Supplementary Fig. S13E). The common regions bound by H2A.X and H2A.W had more enrichment of H2A.W than H2A.X (Supplementary Fig. S13F). Since H3K9me2 marks that generally overlap with H2A.W were enriched in h2a.x ko we analyzed the levels of H3K9me2 in the 5822 common regions preferred by both these variants, and found that this mark enriched much more in h2a.x ko than wild type, indicating that H2A.X suppresses H3K9me2 mark (Fig. 5L). These 5822 genomic regions contained largely TEs (12140 TEs) and 934 genes. The increase in the level of H3K9me2 was more over TEs but not in genes (Supplementary Fig. S13G, H). Even though the gain of H3K9me2 was not seen over these genes, surprisingly, most of these genes exhibited reduced expression in h2a.x ko compared to WT or OE (Fig. 5M). In addition to this several genes which had H2A.X binding exhibited increased expression in ko (Supplementary Fig. S13I).

Since, H2A.X peaks overlapped mostly with H2A.W, and these variants in *Arabidopsis* appear to form homotypic nucleosomes (7), we explored if H2A.X and H2A.W can co-occupy in a single nucleosome in rice using micrococcal nuclease treatment followed by IP (MNase IP). In this assay, we could clearly see that both the variants form homotypic nucleosomes *in vivo* (Fig. 6A), indicating that these variants while competing for specific nucleosomes, cannot occupy one nucleosome together. We performed ChIP seq for H2A.W in h2a.x ko to check if there is any competition between these variants for specific nucleosomes. In fact, there was enriched levels of H3K9me2 over H2A.W peaks obtained in ko (Fig. 6B). Similar to the gain of H3K9me2 over TEs in h2a.x ko, there was increased levels of H2A.W over TEs in h2a.x ko plants, while the overall global distribution of H2A.W was not altered in ko (Fig. 6D and Supplementary Fig. S14A and S14C). The peak width of H2A.W was also similar between WT and ko (Supplementary Fig. S14B). Surprisingly, H2A.X showed reduced enrichment in TSS that is otherwise enriched with active H3K4me3 mark, whereas H2A.W was enriched specifically over TEs in proximal regions (Fig. 6C). We also observed a global redistribution of H2A.W in h2a.x ko where it distributed to the neighbouring non-coding regions sometimes upto a few kb long (Supplementary Fig. S14D). Such a redistribution was confirmed using immunostaining of nuclei in WT and h2a.x ko (Fig. 6E). Rice H2A.W was distributed to the periphery of the nuclei in WT unlike *Arabidopsis* where it forms specific puncta (45). The presence of TEs and heterochromatic regions spread out in rice might have resulted in this pattern of H2A.W in rice. Interestingly, H2A.W showed a more uniform distribution in ko and more signal towards the centre of the nuclei where H2A.X enrichment was observed in WT, indicating that H2A.W can replace this variant (Fig. 6E and Supplementary Fig. S14E, F).

**Fig. 6:**
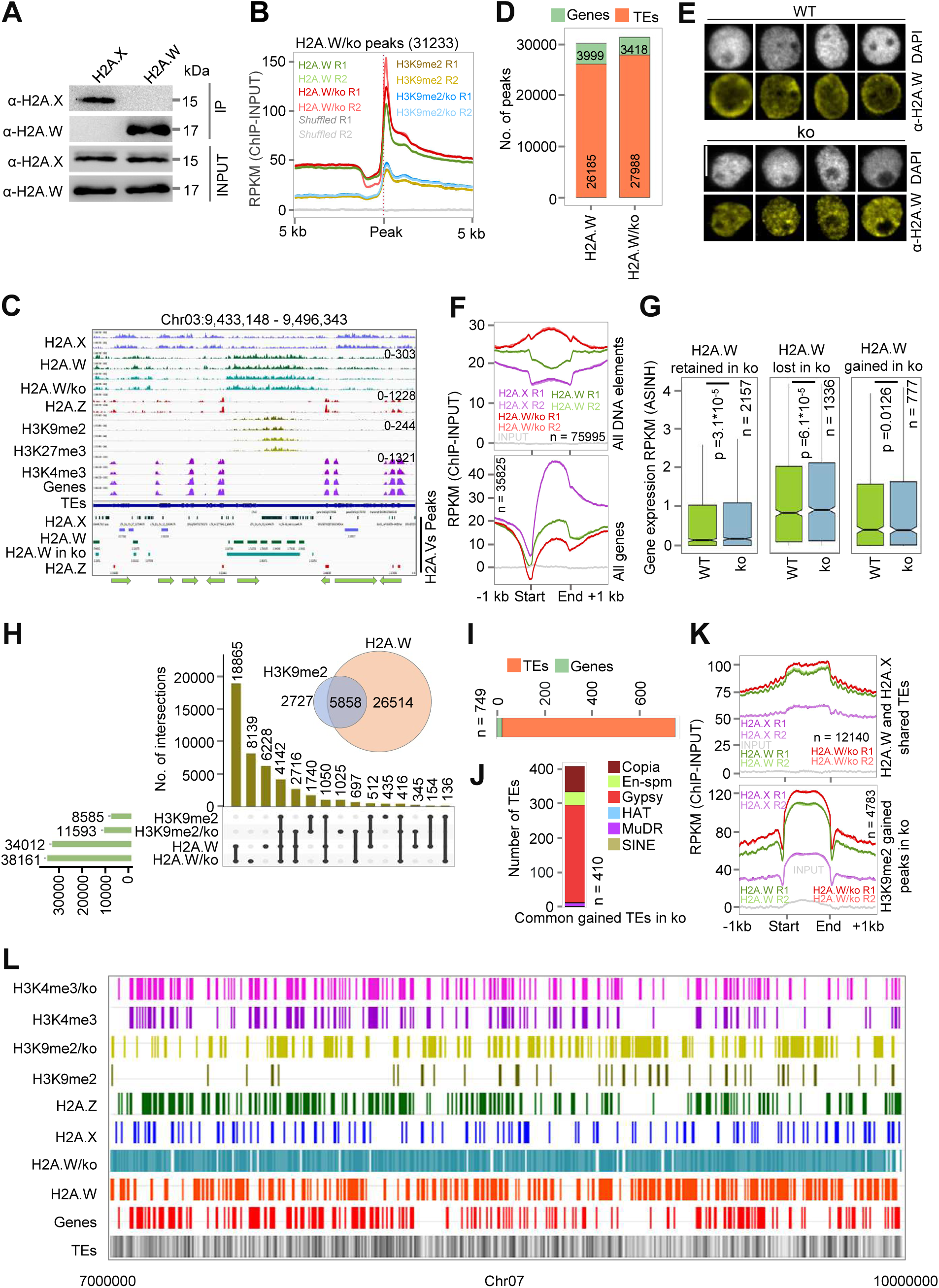
H2A.X inhibits incorporation of histone variant H2A.W to restrict H3K9me2. **A,** MNase IP showing exclusive binding of H2A.X and H2A.W in nucleosomes. **B,** Metagene plot showing enrichment of H2A.W and H3K9me2 in WT and ko over total H2A.W peaks in ko. **C,** Genome wide IGV screenshots showing distribution of H2A variants and histone marks in rice. **D,** Barplot showing number of H2A.W peaks over genes and TEs in WT and ko. **E,** IFL images of nuclei from WT and ko plants stained using α-H2A.W. Scale - 5 μm. **F,** Metagene plot showing enrichment of H2A.X and H2A.W over all TE elements and genes in ko. **G,** Boxplots showing expression of genes in H2A.W altered regions in ko. **H,** Upset plot showing enrichment of H2A.W and H3K9me2 in WT and ko **I,** Barplot showing number of TEs and genes bound by regions that gained both H3K9me2 and H2A.W in ko. **J,** Enrichment of important TE features bound by the regions which gained both H3K9me2 and H2A.W in ko. **K,** Metagene plot showing enrichment of H2A.X and H2A.W over H3K9me2 gained and H2A.X and H2A.W shared TEs in ko. **L,** Chromosome-wide map of specific region showing the distribution of H2A variants and histone marks in WT and ko. Significance calculated by Two-sided Wilcoxon test (*p* < 0.01 was considered significant) in **G**.

### H2A.W and H3K9me2 co-occupies in rice, while increased enrichment of both was seen over TEs in h2a.x ko

In order to understand the genome wide re-distribution of H2A.W in ko, we intersected the peaks overlapping with genes and TEs in WT and ko. The enrichment of H2A.W over genes was reduced in ko while it increased over TEs (Fig. 6D, F and Supplementary Fig. S14H). Such an increase of H2A.W did not distinguish types and families of TEs (Supplementary Fig. S14G, H). Increased occupancy of H2A.W over several TEs did not alter the expression levels of the TEs or sRNA from these regions in h2a.x ko, probably because of its low expression in WT conditions (Supplementary Fig. S14I, J). There was also a redistribution of H2A.W peaks over several genes in ko. There were around 1336 genes that lost H2A.W, while 777 genes gained H2A.W in h2a.x ko lines. Surprisingly, the genes that lost H2A.W in h2a.x ko showed increased expression, indicating the repressive role of H2A.W over these genes in WT conditions (Fig. 6G, Supplementary Fig. S15A-C). As expected, most of the H3K9me2 marks (5858) overlapped with H2A.W in WT condition while there were 697 regions which gained both H2A.W and H3K9me2 in h2a.x ko (Fig. 6H). These new H2A.W and H3K9me2 acquired peaks mostly overlapped with TEs, specifically of Gypsy family (Fig. 6I, J). Surprisingly, we could observe increased level of H2A.W in ko over H3K9me2 peaks obtained in h2a.x ko, suggesting there is a global level increase of H2A.W and H3K9me2 in h2a.x ko (Supplementary Fig. S15D). In addition to this, we observed higher levels of H2A.W over H3K9me2 gained regions (4783), as well as H2A.X and H2A.W bound common TEs (12140) (Fig. 6K, Supplementary Fig. S15E). The increase of H2A.W over H3K9me2 gained regions was also mostly in TEs, including all DNA elements and LTRs (Supplementary Fig. S15F). Among the 697 regions that overlapped with 21 genes where H2A.W and H3K9me2 gained in h2a.x ko, most of the genes were very less expressed even in WT conditions (Supplementary Fig. S15H). RT-qPCR analysis of two selected genes that gained H3K9me2 and H2A.W in promoter, Os07g0430501 (*Thionin-like peptide*), an antimicrobial peptide and Os04g0269600 (*OsDA02*), which is important for oxidative deamination in plants, showed clear downregulation of gene expression, indicating that H2A.X positively regulated their expression in WT condition by inhibiting both H2A.W and H3K9me2 repressive marks (Supplementary Fig. S15I). Several of the genes in this category did not show clear downregulation in h2a.x ko, probably because of their very low expression in WT seedlings, or involvement of other regulations (Supplementary Fig. S15G, H). Surprisingly, several genes which gained H3K9me2 mark in ko and exhibited reduced expression in ko had increased levels of H2A.W in the H3K9me2 gained regions, suggesting the collective gain of both these repressive marks in these genes in ko (Supplementary Fig. S16). These results indicate that, although H2A.X suppresses gene expression, its competition with H2A.W and suppression of H3K9me2 ensures expression of a few specific genes. Apart from the collective gain of H2A.W and H3K9me2 in specific gene promoters, there were large chromosomal regions which gained both these marks in ko (Fig. 6L).

### Development-associated regulatory role of H2A.X is largely independent of its role in DNA damage responses

Since H2A.X is a well-known player in DNA damage response across eukaryotes, we explored if development-associated genes under its control are also contributing to DNA damage response. Since DNA damage response across plants requires H2A.X phosphorylation at C-terminal serine position, we expected the OE lines to mimic DNA damage (51). We probed DNA damage responsive genes in rice and found that H2A.X is the major histone variant which is bound to these genes (Fig. 7A and Supplementary Data S9). We analyzed the expression levels of DNA damage responsive genes with H2A.X mis-expression lines in unstressed conditions, and found that several of these genes are upregulated in H2A.X OE, indicating that H2A.X is indeed a regulator of these genes under its overexpression conditions (Supplementary Fig. S17A). Most importantly, the levels of DNA damage responsive genes were not altered in h2a.x ko, indicating that the genes that are directly regulated by H2A.X for development are not among the DNA damage response genes. Further, we treated WT plants with DNA damaging agent bleomycin (B5) (5 mg/litre) which is a slow DNA damaging agent that induces DNA damage over a time period as well as with UV treatment (3KJ) which is a quick DNA damaging agent (Supplementary Fig. S17B). Surprisingly the OE plants showed similar expression pattern to WT plants upon treatment with bleomycin but not with UV treatment (Supplementary Fig. S17C, D). In addition to this, the expression of DEGs in H2A.X OE plants was similar between H2A.X OE and WT plants treated with bleomycin (Fig. 7B). Further we selected the bleomycin responsive DNA damage responsive genes which showed higher expression under bleomycin treatment in WT and found that similar to all DNA damage responsive genes, bleomycin responsive genes are also bound predominantly by H2A.X (Supplementary Fig. S17E-G and Supplementary Data S9). These bleomycin responsive genes consist of several well-known DNA damage responsive genes which comes under both nucleotide excision repair (NER) and DNA double strand break repair (DSB) pathway and contains well-studied genes like OsMORC and OsRAD family proteins (Supplementary Fig. S17H, I). The expression of these bleomycin responsive genes was upregulated in H2A.X OE without any DNA damage and H2A.X OE plants mimicked bleomycin treatment, suggesting the enhanced stress response activated in OE plants without any DNA damage (Fig. 7C, D, Supplementary Fig. S18A and Supplementary Data S10). In addition to this H2A.X OE plants which exhibited DNA damage sensitive phenotypes in normal condition were resistant to external DNA damage given by bleomycin treatment while the h2a.x ko plants were very sensitive to DNA damage (Supplementary Fig. S18B-D). As shown in *Arabidopsis* previously the root growth of h2a.x ko plants were much more reduced under DNA damage condition, while OE plants were largely unaffected mimicking enhanced DNA damage response (Supplementary Fig. S18B-G).

**Fig. 7:**
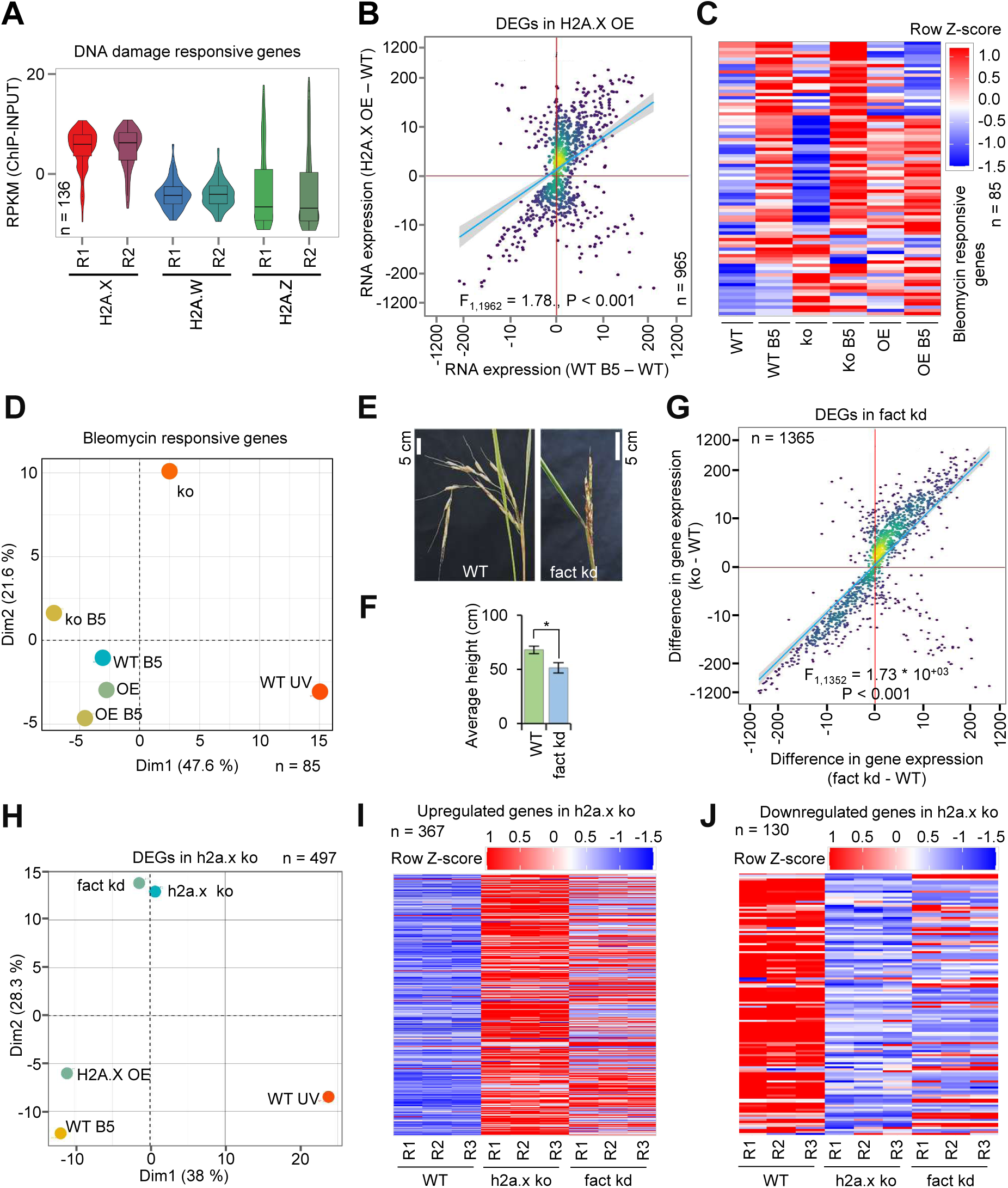
DNA damage associated role of H2A.X is independent of its role in development. **A,** Boxplot showing enrichment of H2A variants over DNA damage response genes. **B,** Density scatter plot displaying comparison of H2A.X OE DEGs and their expression upon bleomycin treatment in WT plants (WT B5 (lines treated with 5 mg/litre bleomycin)). **C** Heatmap showing expression of bleomycin responsive DNA damage repair genes in H2A.X misexpression lines. **D,** PCA of bleomycin responsive genes in H2A.X mis-expression lines. **E,** Panicle phenotypes of WT-PB-1 and fact kd. **F,** Boxplot showing average height of fact kd plants. **G,** Density scatter plot displaying similarity of fact kd DEGs and their expression in ko. **H,** PCA analysis of different stress datasets with ko DEGs. Heatmap showing expression of genes upregulated in ko **I,** and downregulated in ko **J,** in fact kd lines. For **B** and **G**, point density is color coded. Blue line represents the linear regression fit line, and grey shade represents the 95% confidence interval. Difference in gene expression is plotted and x- and y-axis are scaled to inverse sine hyperbolic function. Fischer’s test was used for statistical testing of the dependence of two perturbations. Two tailed student’s *t*-test was used for statistical comparison in **F**.

### Phenotypic and molecular analysis of loss of function mutants of histone chaperone of H2A.X correlated with the gene regulatory role of H2A.X

In order to confirm the regulatory role of H2A.X in gene expression, we generated double kd lines of Suppressor of Ty (SPT) 16, the partner of STRUCTURE SPECIFIC RECOGNITION PROTEIN 1 (SSRP1) in forming the FACT complex (Supplementary Fig. S19A). Probably since the protein complex is critical for several biological processes, and knocking out these partners lead to lethality as shown in different systems (57, 59, 81), we failed to generate viable ko lines. The transgenic lines with abundant levels of amiR expression and reduced target mRNA levels were viable and were selected for further analysis. The transgenic lines had reduced expression of both the isoforms of SPT16 (Supplementary Fig. S19B, C). These amiR plants showed strong developmental and reproductive defects including reduced height and abnormal panicles (Fig. 7E, F). Although h2a.x ko plants showed similar phenotypes, it is important to note that only a part of phenotypes observed in fact kd lines might be due to H2A.X. In order to understand whether the gene expression changes observed in h2a.x ko plants are also controlled by FACT complex, we performed RNA seq of 14-day old fact kd lines. As expected, the expression of both the isoforms of SPT16 were reduced in fact kd lines (Supplementary Fig. S19D). We obtained around upregulated 807 genes and downregulated 558 genes in fact kd lines and these genes had similar enrichment of GO category as observed among h2a.x ko DEGs (Fig. 3B, Supplementary Fig. S19E, F and Supplementary Data S11). Surprisingly, the DEGs in fact kd lines were similar in expression pattern in h2a.x ko plants, suggesting that the RNA levels in fact kd lines are also reflected in h2a.x ko lines (Fig. 7G and Supplementary Fig. S19G). Most of the upregulated and downregulated genes in h2a.x ko and fact kd lines were showing similar expression pattern in fact kd lines and h2a.x ko (Fig. 7H-J and Supplementary Fig. S19G, H). Similarly, several developmental, reproductive and stress associated genes were mis-expressed similarly in h2a.x ko and fact kd lines (82–87). In these genes, most of the common upregulated genes had increased levels of H3K4me3 while common downregulated genes did not show enrichment for H3K9me2 (Supplementary Fig. S20A, B). These results indicate that the developmental role of H2A.X is partly attributed to the abnormal phenotypes in the mutants of its chaperone complex.

## Discussion

Histone variants are conserved across different organisms and mostly provide specific functions important for development and to mediate stress responses, sometimes in a tissue-specific manner (6, 7, 17, 23–26, 88). Functions of H2A variants are well explored in plants, including those that are unique to them, partly because they exhibit clear amino acid variations in C-terminal tail*s* (80, 89). Variations in the C-terminal tails such as the presence of multiple H2A.W isoforms with duplicated SPKK motif are observed in rice, but functional studies on these aspects are yet to be carried out (29, 45). Most of the H2A variants exhibit tissue specific expression in rice with H2A.X having high expression in seedling and reproductive tissues. It is not surprising that rice h2a.x ko plants displayed defects in both seedling and reproductive development. The increased root growth phenotype observed in rice h2a.x ko and kd was also observed in *Arabidopsis* mutant under ABA treatment (69). Several genes including *OsHAC1*, *Os1-CysPrxB*, *OsRCc3, EXO70FX15* and *OsZFP350*, involved in positively regulating the root growth were upregulated in h2a.x ko (90–93). The chlorophyll content in both h2a.x ko and kd lines were high compared to WT, and the photosynthesis associated genes that exhibited higher enrichment of H2A.X including OsHPR1 and CDSP32 were upregulated in h2a.x ko (94, 95). These results document previously unappreciated role of H2A.X in mediating specific development-associated processes.

The distribution of H2A.X in gene bodies per se, along with TE elements suggests its potential in regulating these features. The general trend of H2A.X enrichment over gene body and its absence over TSS is an opposite pattern to that of the H3K4me3 enrichment over TSS and its reduced levels on the gene bodies. The enrichment of H2A.X over gene body might be specifically ensuring that active H3K4me3 marks are enriched over TSS but not in the gene bodies. Such a role for H2A.X can be proposed based on the observations that the levels of H3K4me3 over TSS was much more in H2A.X bound genes when compared to all other genes. Also, H2A.X bound genes displayed higher expression compared to genes bound by other H2A variants. More importantly, the genes that gained H3K4me3 in h2a.x ko had increased enrichment of H3K4me3 over gene bodies but not in TSS, further suggesting the regulatory role of H2A.X in influencing H3K4me3 marks. Molecular basis for such a repressive function of H2A.X is unclear and these might be either due to increased occupancy of other histone variants or canonical H2A. The repression of H3K4me3 by H2A.X was seen even over TEs with a functional consequence, since most of the TEs that gained H3K4me3 in ko plants exhibited increased expression (Fig. 4l). Recent studies indicate that H3K4me3 mark can induce DNA de-methylation by attracting DNA de-methylases in plants and it will be interesting to check whether such a mechanism is seen in H2A.X associated H3K4me3 gained genes and TEs (96). Signatures of suppression of gene expression activity of H2A.X differs from other H2A variants such as H2A.Z, since H2A.Z bound genes enrich with repressive H3K27me3 marks and the inducibility of H2A.Z occupied genes under stress is very different from highly expressed active genes having both H3K4me3 and H2A.X (35, 38, 97–99). The Polycomb group proteins (PcG) consist of polycomb repressive complex 1 and 2 (PRC 1 and 2) regulates H2A and H2A.Z ubiquitination to induce H3K27me3 marks at several development-associated genes (43, 97, 100). The C-terminal lysine residue required for this function is conserved in H2A.X (101), hence, there is a possibility that PRC-mediated control intervenes H2A.X functions but this aspect is not explored.

The ALFIN-LIKE (AL) proteins contain a C-terminal PHD finger that binds to H3K4me3 modification and facilitates H2AUb in canonical H2A and H2A.Z variant throughout the genome, especially over genes (102). AL proteins interact with the SWR1 remodelling complex which is a chaperone of H2A.Z (103). For AL proteins to act on the genome, H3K4me3 mark is essential and therefore it will be interesting to see if the incorporation of either H2A or H2A.Z in place of H2A.X in ko lead to increased levels of H3K4me3 through the H2AUb. Such a regulation in plants is highly unlikely at least with H2A.Z, since this variant inhibits active H3K4me3 from gene promoters to promote repressive H3K27me3 marks (38).

ChIP-seq of FACT complex in *Arabidopsis*, a known chaperone of H2A.X (54, 55), revealed its binding to the euchromatin regions containing active genes, whereas it was less enriched over heterochromatic regions (58). The distribution of FACT complex with actively expressed genes is very similar to the enrichment of H2A.X over active genes which is observed in both rice and *Arabidopsis*. Similar to altered gene expression that we observed here, there are several studies in *Arabidopsis* that mention the gene regulatory function of FACT complex (59, 104). Similarity between the transcriptome of fact kd and h2a.x ko plants suggests that at least a part of gene expression change in fact kd is associated with the defects in H2A.X incorporation. The involvement of FACT complex in regulating anthocyanin accumulation under stress conditions correlates with the increased levels of anthocyanin observed in H2A.X OE plants (105). The shared developmental and reproductive defects observed in h2a.x ko and fact kd along with shared transcriptomic changes are interesting correlations, however, it is also important to note that FACT complex has a bigger role in regulating several key processes including transcription and replication and hence, secondary effects cannot be ruled out (106). FACT complex is crucial for suppressing the transcription from intergenic TSS by regulating H3K4me1 and H3K4me3 marks in *Arabidopsis* (107). The suppressive role of H2A.X in H3K4me3 deposition is similar to the suppressive role of FACT complex in preventing deposition of H3K4me3 over intergenic TSS. It will be interesting to see what proportion of the genome-wide H3K4me3 levels are dependents on H2A.X and FACT complex. It is striking to note that the gene regulatory function of H2A.X under DNA damage state is de-coupled from its role in development. The influence of H2A.X in regulating global DNA damage repair might be linked to its well-known PTM, a phosphorylation at the conserved SQEF motif at the C-terminal tail. Interestingly, several genes that were upregulated under bleomycin induced DNA damage were already upregulated in H2A.X OE. Moreover, the H2A.X OE plants showed severe stressed phenotypes, some of which can be correlated with DNA damage response, even without any DNA damage. The H2A.X OE plants might have accumulated high levels of gamma H2A.X protein mimicking DNA damage. Surprisingly, these OE plants were resistant to DNA damage, even the well-known DNA damage-related phenotypes including reduced root growth, was not observed in these plants. It is known across several model systems that phosphorylated H2A.X helps in upregulating gene expression either by binding to the nicked DNA to increase the transcription from these sites, or by partnering with different proteins to bind to the promoter of specific genes (67–69). It will be interesting to check if that kind of gene expression trigger based on H2A.X phosphorylation is seen in H2A.X OE plants to specifically regulate the DNA damage responsive genes.

The nature of specific H2A variants binding to specific chromatin types and regions might not be similar across flowering plants. It has been well established that occupancy of heterochromatic regions in the nuclei among diverse plants showed unique properties (17, 108). Any variation in the distribution of chromatin territories has the potential to alter functions of histone variants that have specific regulatory regions to decorate. The distribution of TEs is strictly towards the heterochromatic regions including centromeric repeats and telomere containing chromatin arms in *Arabidopsis* (109). However, in plants with larger genomes including rice, the TEs are distributed across the genome interspaced with genic regions (8, 9). Enrichment of rice H2A.W in the TEs proximal to protein-coding genes was clearly observed, as was the H2A.X enrichment in the gene bodies adjacent to them. This pattern of distribution might have led to the enrichment of H2A.W in ko plants contributing to the altered expression of genes coming under the control of H2A.X and its competitor H2A.W. This diversity of functions of variants between two species is extremely interesting and a valuable starting point to explore its evolutionary significance. The genes bound by H2A.W were very diverse and low expressed when compared to genes bound by other H2A variants. The increased enrichment of H2A.W over TE regions in ko plants suggests that H2A.X also has a regulatory role in H2A.W enrichment of non-coding regions. Unlike in *Arabidopsis,* among monocots including rice, the nuclear puncta representing the centromere is not formed when H3K9me2 mark is assayed (108). Similarly, H3K9me2 marks in rice and wheat show distribution in the periphery of the nuclei without forming the puncta. Similar distribution of H2A.W in rice and strong overlap of H3K9me2 and H2A.W peaks suggests that these two features are important in regulating heterochromatin in rice. It is also known that H2A.W restricts H1 to provide access for DNA methyltransferases to the centromeric repeats (110). Even though the DNA methylation in the specific targeted regions was not altered in h2a.x ko in rice, it will be interesting to check if global DNA methylation pattern gets altered. Curiously, increased expression of specific TEs in ko plants, resulting from increased H3K4me3 levels, did not induce RNA-directed DNA methylation. It is possible that other regulatory functions initiated by sRNAs might be active in ko lines that we did not explore. Most of the regions with H2A.X and H2A.W peaks were on TEs, often creating mosaics in the rice genome. These TEs exhibited increased levels of both H2A.W and H3K9me2 levels in ko, further suggesting the competition of H2A.X and H2A.W over TE regions. Surprisingly, the increased enrichment of H2A.W was enough to enrich H3K9me2 in these regions and also globally increased levels of H2A.W was observed in the peaks of H3K9me2 in ko plants. The evolutionary significance of specific distribution of H2A.W and H3K9me2 and reasons why specific genes and TEs are enriched in such regions will be worth exploring. It will be also interesting to explore how the chromatin remodeller DDM1, which is the chaperone of H2A.W, functions to maintain H2A.W to regulate heterochromatin processes in rice and how H2A.X influences its functions.

## Methods

### Plasmid construction and cloning

To generate CRISPR based double knockout plants of H2A.X, gRNAs were cloned in to pYLCRISPR/Cas9Pubi-H-based construct (111) using BsaI sites. The gRNA1 (5’ cguccaggacgaaggguucu 3’) to target OsHTA704 was cloned under OsU6a promoter whereas the gRNA2 (5’ tcccgagguucagccacagc 3’) to target HTA711 was cloned under OsU3 promoter. To generate overexpression (OE) plants, rice *H2A.Xa* (OsHTA704) CDS was amplified (the primers are listed in Supplementary Table S3) from PB-1 seedling RNA with Superscript III reverse transcriptase (Invitrogen). The amplified CDS was inserted with a N-terminal FLAG tag in to a derivative of pCAMBIA1300 plasmid under maize Ubiquitin promoter. For double artificial microRNA mediated knockdown (amiR) lines of OsH2A.X and OsSPT16, OsamiR528 coding region was swapped using specific amiR sequence in pNW55 vector with the help of WMD3 tool. For generating h2a.x kd lines, amiR1 (3’ aacgacagagccagcagguuu 5’) was cloned to maize Ubiquitin promoter while amiR2 (3’ agcgacaccgccagcagguuu 5’) was cloned under 35S promoter. For fact kd plant generation amiR1 (3’ ugcacguauuaagacuagauu 5’) was cloned to 35s promoter while amiR2 (3’ agcagaguucuagaagcaauu 5’) was cloned under maize Ubiquitin promoter. All these binary vectors were mobilised to *Agrobacterium tumefaciens* strain LBA4404 with additional virulence plasmid pSB1.

### Plant transformation

Pusa basmati-1, indica rice variety (*O. sativa indica*) was used for the study. The plants were maintained in the greenhouse in 22 °C. Rice transformation for generating transgenic rice plants were performed by following already standardised protocol (112–114). PB-1 rice seeds were dehusked and sterilized using 70% ethanol, 4% Bleach and 0.1% mercuric chloride. Seeds were placed in callus induction media (CIM) for 21-days in dark to induce callus. These calli were used for *Agrobacterium* mediated plant transformation and further selected using hygromycin. The regenerated plants were transformed to ½ MS media for rooting and further changed to soil.

### RT-qPCR

RNA was isolated from different tissues using Trizol reagent and qPCR analysis was performed using cDNA made from the total RNA using Thermo RevertAid RT kit following manufacturer’s protocol. SYBR green master mix (Solis Biodyne- 5x HOT Firepol Evagreen qPCR master mix) was used for qPCR. *OsActin* (LOC_Os03g50885) or *OsGAPDH* (LOC_Os04g40950) was used as internal control. All the primers (Table 1) and gene ID of the genes (Table S2) are mentioned.

### Phylogeny

The phylogenetic tree was made using MEGA11 tool using maximum likelihood method with bootstrap values (1000 replicates) (115).

### Southern hybridisation

Southern hybridisation to confirm the presence of T-DNA was performed as described previously (116, 117). The hygromycin probe was amplified and labelled using Rediprime random primer labelling kit (Cytiva).

### sRNA northern hybridisation

Northern blot to detect the sRNA was performed as mentioned previously (118, 119). For blotting around 8 µg of total RNA was loaded on denaturing gel and electrophoresed. These sRNAs were blotted to a Hybond N+ membrane (Cytiva) and crosslinked using UV light. The membrane was hybridized with specific radiolabelled DNA (T4 PNK, M0201, New England Biolabs (NEB) probes which was amplified using PCR. Overnight hybridisation of the membrane and the probe was performed in ULTRAhyb buffer at 35 °C. The membrane was exposed to a phosphor screen and then scanned using Typhoon scanner (Cytiva).

### Chlorophyll and anthocyanin estimation

Chlorophyll estimation was done using 200mg tissue of 14-day old seedling leaves using previously mentioned protocol (120). Anthocyanin estimation was performed from 200mg stem tissue of 14-day old rice seedlings as described previously (121, 122).

### Immunofluorescence and microscopy

Immunofluorescence and microscopy was performed as described previously (45, 123). 14-day-old seedlings were crosslinked using 1% PFA was used to isolate nuclei and these nuclei were fixed to a positively charged slide. anti-H2A.X (1:100) and anti-H2A.W (1:100) antibodies was used as the primary antibodies and these were detected using fluorescently tagged secondary antibodies (Invitrogen, goat-raised anti-mouse 488 and anti-rabbit 555). The nuclei were also counterstained using DAPI to detect the DNA and the slides were imaged using confocal microscope (Olympus FV3000).

### Antibody generation

Polyclonal antibodies are made for OsH2A.X and OsH2A.W with specific peptides. Peptide C-GGGKGDIGSASQEF-GGKDKADIGSASQEF was used for OsH2A.X (both the isoforms) while C-KEGKTPKSPKKATTKSPKKA was used to make OsH2A.W (OsH2A4) antibody. Both the antibodies are raised in rabbit and antigen-affinity purified by LifeTein LLC.

### Transcriptome analysis

Transcriptomic analysis was performed as mentioned previously (124). 14-day old whole seedlings were used for RNA sequencing. Total RNA was extracted using TRIzol followed by poly(A) enrichment before sequencing. The libraries were sequenced using Novaseq6000, 2×100 bp format was used for all other sequencing. For all the samples more than 30 million reads were obtained. The adapter content from the obtained reads were removed using Trimmomatic (125) and aligned to the IRGSP-1 genome using HISAT2 (126). Cufflinks were used to analyse the difference in gene expression and transcript abundance. Log2 (fold change) cut off greater than 1.5-fold with P-value cutoff lesser than 0.05 compared to WT sample was selected as DEGs (Supplementary Data S1, S2, S3 and S11). Specific plots were made using R and gene ontology analysis was performed using ShinyGO v0.81 platform (127). The IRGSP gene IDs were used and GO categories were found using general biological processes with FDR cut-off of *p*-value:0.05. For enrichment studies BEDtools multicov was used and the values were normalised to RPKM (128). Multistress dataset analysis was performed using TENOR datasets (https://tenor.dna.affrc.go.jp/) as described previously (17, 74). Photosynthesis, pigmentation, and root growth genes are selected from AgriGO https://systemsbiology.cau.edu.cn/agriGOv2/ and Funrice https://venyao.xyz/funRiceGenes/ (129). For DNA damage response genes ensemble plants GO category genes under DNA damage response was used.

### Targeted bisulphite sequencing

The targeted bisulphite sequencing was performed using protocol mentioned previously (130). For targeted bisulphite sequencing around 200-400ng of DNA was isolated from 14-day old seeding and treated using EZ DNA Methylation-Gold Kit (Zymo Research). PCR for specific promoter regions was performed using this converted DNA as template using JumpStart™ Taq DNA Polymerase (Sigma). The obtained PCR products from transgenic lines were sequenced using paired end mode (100 bp) in Hiseq2500 platform. The obtained reads were trimmed using cutadapt and aligned to target regions generated using Bismark aligner tool (131–133). Obtained results were analyzed using methylation package under ViewBS and plots are modified using R (133).

### Bleomycin treatment and phenotyping

For bleomycin treatment WT and transgenic seeds were sterilized and germinated in ½ strength MS (Murashige and Skoog) medium with 0.3% Phytagel (Sigma Aldrich). 3 days after germination plants were changed to ½ MS media supplemented with 5mg/litre bleomycin sulphate (European Pharmacopoeia (EP)). These plants were grown for another 14 days and used for phenotyping and RNA sequencing.

### Western blotting and MNase IP

For western blotting analysis total nuclear protein was isolated from 500mg of seedling tissue otherwise mentioned specifically, using protocol described previously (29). These histones were electrophoresed in 14% SDS-PAGE and transferred to Protran supported nitrocellular membrane (134). These membranes were hybridized with specific antibodies anti-H2A.X (custom generated, 1:1, 000), anti-H2A.W (custom generated, 1:1, 000), anti-H3K9me2, anti-H3K4me3 and anti-H3 (Merck 07-10254, 1:20, 000) antibodies in a 5% milk in 1x TBST buffer. Bacterial protein expression, bacterial induced cells were grown upto OD 2.0 and lysed by heating with 2x Laemmli buffer. MNase IP was performed from seedling tissues by following previously mentioned protocol (29).

### ChIP sequencing and analysis

For all ChIP sequencing experiments 14-day old seedlings were used. Around 1.5 g of 1% formaldehyde crosslinked tissue was used for nuclei isolation and chromatin shearing. Sheared chromatin was incubated with 5 µg of anti-H2A.X, anti-H2A.W, anti-H3K9me2 and anti-H3K4me3 antibodies overnight at 4 °C. These chromatin fractions were incubated with protein-G Dynabeads (Thermo Fisher Scientific) for four hours. Purified DNA from the IP was used prepare ChIP seq library using NEBNext Ultra II DNA library prep kit (NEB E7103) using the manufacturer’s protocol. The sequencing of ChIP seq libraries were performed in NovaSeq 6000 platform (Supplementary Table S1). Adapter content from the sequencing reads were removed using Cutadapt and mapped the reads to IRGSP-1 genome using Bowtie (131, 135) using parameters *v*, 1; *k*, 1; y; a; best; strata. PCR duplicates from the aligned reads were removed and then converted the reads to coverage tracks by subtracting the reads to the sheared input DNA using deepTools (136). Enrichment was calculated using ComputeMatrix under deepTools and plotprofile tool was used to make plots. These plots were modified using custom R script using ggplot2 package (137). ChIP peaks were obtained from each experiment using MACS2, broad peaks were selected for histone variants (H2A.X and H2A.W) while narrow peaks were called for histone marks (H3K9me2 and H3K4me3). All these peaks had a significant enrichment of 2-fold compared to input sheared DNA. These peaks were merged and ChIPseekeR was used to find the chromosomal regions of enrichment. Publicly available datasets were also analyzed the same way (Supplementary Table S2). shinyCircos-V2.0 (https://venyao.xyz/shinycircos/) and ShinyChromsome (https://venyao.xyz/shinyChromosome/) tools were used for generated circos and chromosome wide plots respectively.

## Data access

All raw and processed sequencing data generated in this study have been submitted to the NCBI Gene Expression Omnibus (GEO; https://www.ncbi.nlm.nih.gov/geo/) and these are accessible *via* accession numbers GSE308770, GSE308771, GSE308772.

## Acknowledgements

We thank Prof. K. Veluthambi for *Agrobacterium* strains, PB-1 seeds, and binary plasmids. We thank central imaging facility (CIFF), genomics, electron microscopy, IT, radiation, greenhouse, and laboratory-kitchen facilities at the NCBS. We thank all the laboratory members for discussions and comments. This work was supported by NCBS-TIFR core funding, Department of Atomic Energy, Government of India, project identification no. RTI 4006 (1303/3/2019/R&D-II/DAE/4749 dated 16.7.2020) and grant number BT/PR53711/BSA/33/312/2024 from Department of Biotechnology, and grant number CRG/2023/003849 dated 23.03.2024 from SERB, Government of India. AM and SRS thanks Council of Scientific and Industrial Research (CSIR) for the funding. These funding agencies did not participate in the designing of experiments, analysis, or interpretation of data or in writing the manuscript.

## Author contributions

P.V.S. designed all experiments, discussed results, and wrote the manuscript with A.M. A.M generated the h2a.x ko and fact kd transgenic lines, performed most of the experiments and bioinformatic analysis. V.H.S. generated most of the transgenic lines, and discussed results. S.R. and R.D. performed microscopy. All authors have read and approved the manuscript.

## This study contains the following Supplementary materials

### Supplementary Figures S1 - S20

**Supplementary Fig. S1.**
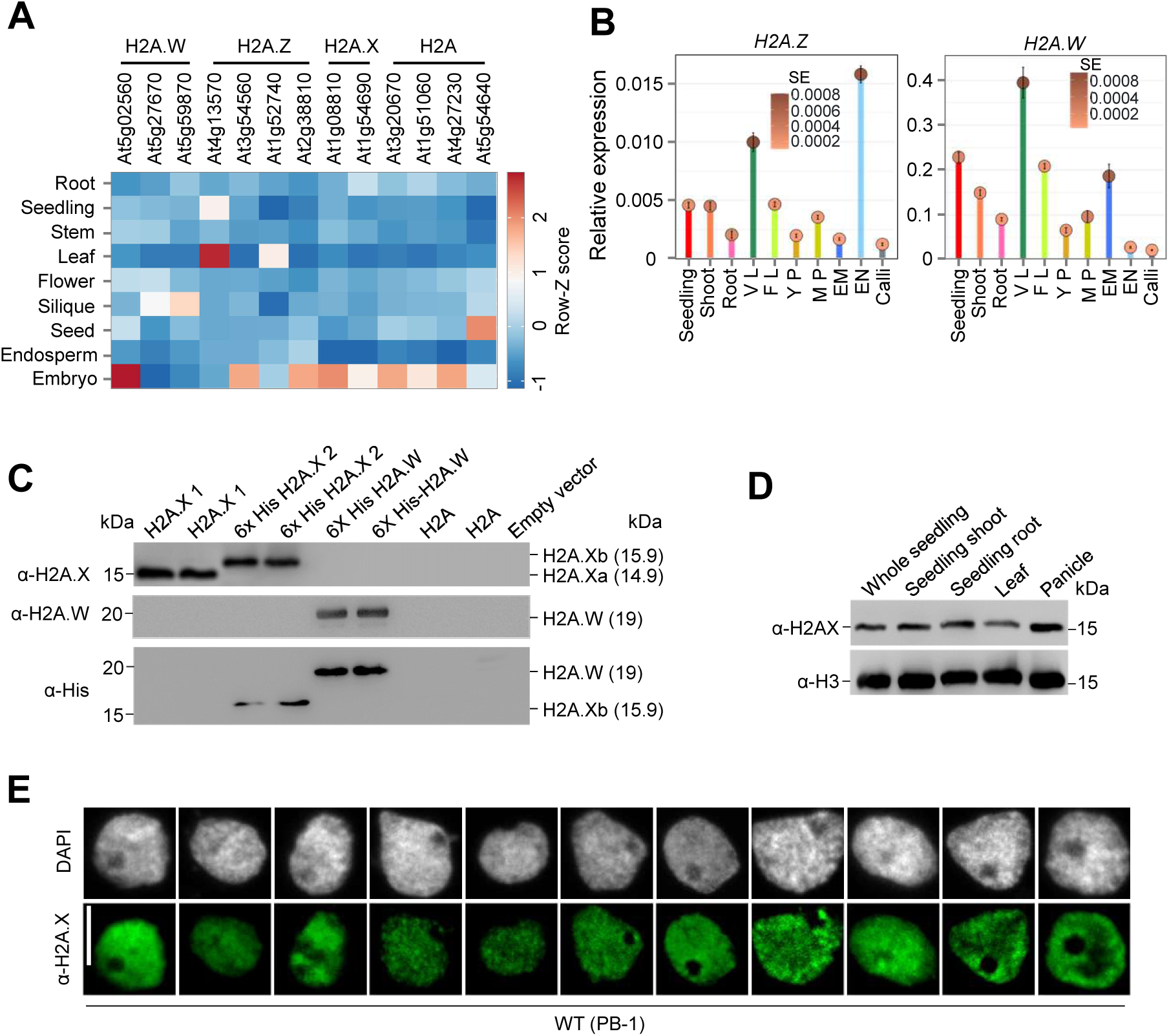
Validation of H2A.X and H2A.W specific antibodies. **A,** Heatmap depicting tissue-specific expression of H2A variants in *Arabidopsis*. **B,** Relative expression of *OsH2A.Z* and *OsH2A.W* across different tissues. *OsGAPDH* served as an internal control. Data represents means ± SE, n = 3. qRT-PCR experiments are performed twice with consistent results. **C,** Western blot showing validation of α-H2A.X and α-H2A.W specific antibodies against different H2A variants expressed in *E.coli* codon plus cells. **D,** Western blot showing tissue specific enrichment of H2A.X. **E,** IFL images of PB-1 nuclei probed with α-H2A.X antibody. Scale bars, 5 µm. H3 is used as the loading control in **C** and **D**.

**Supplementary Fig. S2.**
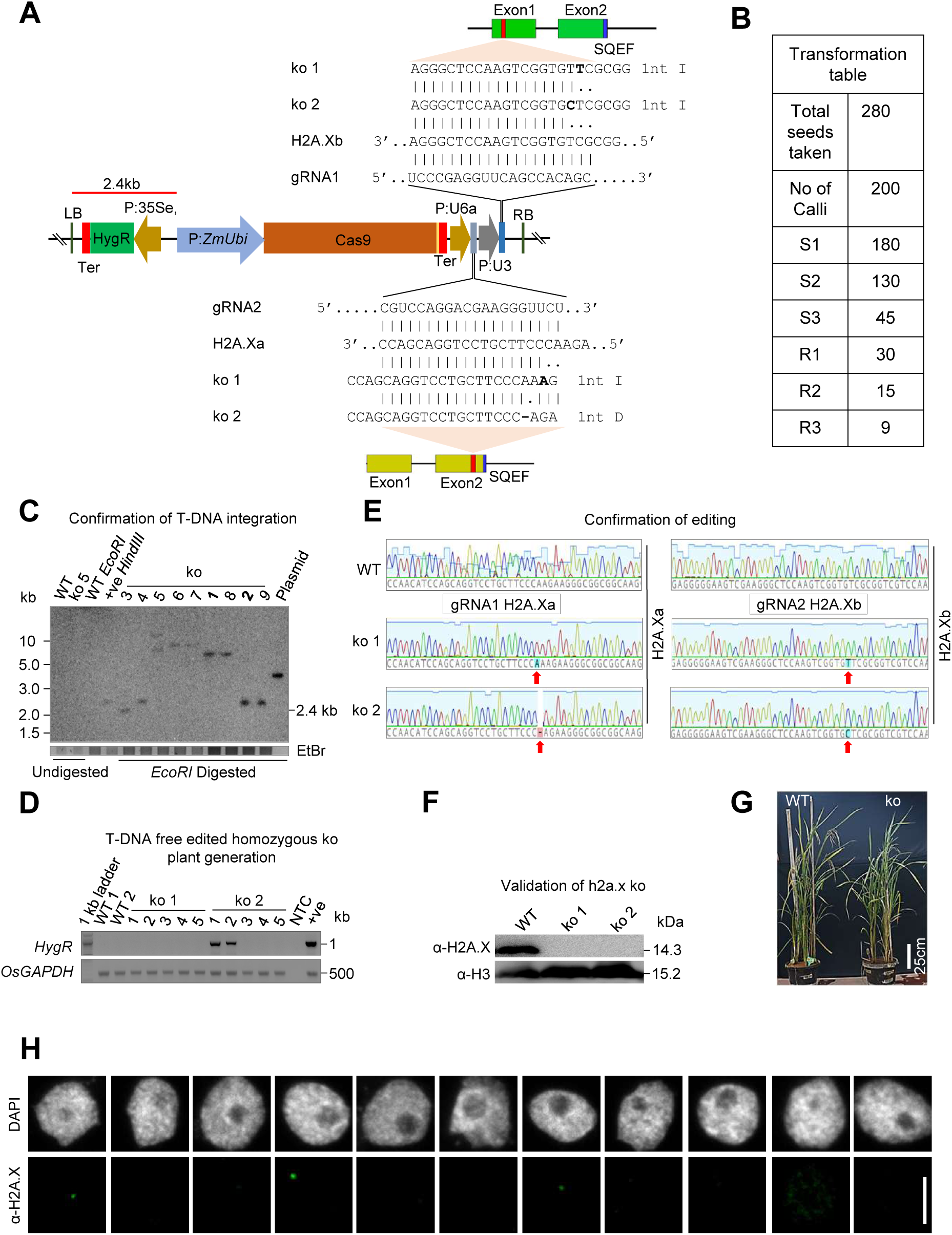
Generation and validation of H2A.X double knockout (ko) lines. **A,** T-DNA map showing gRNA targeting both the isoforms of H2A.X. The target sites and the mutations in ko 1 and ko 2 are displayed. **B,** Summary of *Agrobacterium*-mediated transformation to generate ko plant (S-selection, R-Regeneration). **C,** Junction fragment Southern blot showing copy number of T-DNA in ko lines. DNA fragment corresponding to the digestion of gDNA is shown in red coloured line (2.4kb). Minimum fragment length of the DNA fragment is mentioned in the blot. Single copy insertion lines from ko 1 and ko 2 are selected for further studies. EtBr-stained gel served as loading control. **D,** RT-PCR (semi-quantitative) analysis showing generation of T-DNA free lines from ko 1 and ko 2 plants. NTC-no template control. *OsGAPDH* was used as the loading control. +ve – positive control plant with *HygR.* **E,** Chromatogram showing the genome edited region in PB-1 and ko for both the genes. The gRNA region is highlighted. **F,** Western blot showing the absence of H2A.X protein in both the ko lines. H3 is used as the loading control. **G,** Phenotypes of H2A.X mis-expression lines (90 days old). **H,** IFL images showing absence of H2A.X protein in the nuclei derived from ko plants. Scale bar-5 µm.

**Supplementary Fig. S3.**
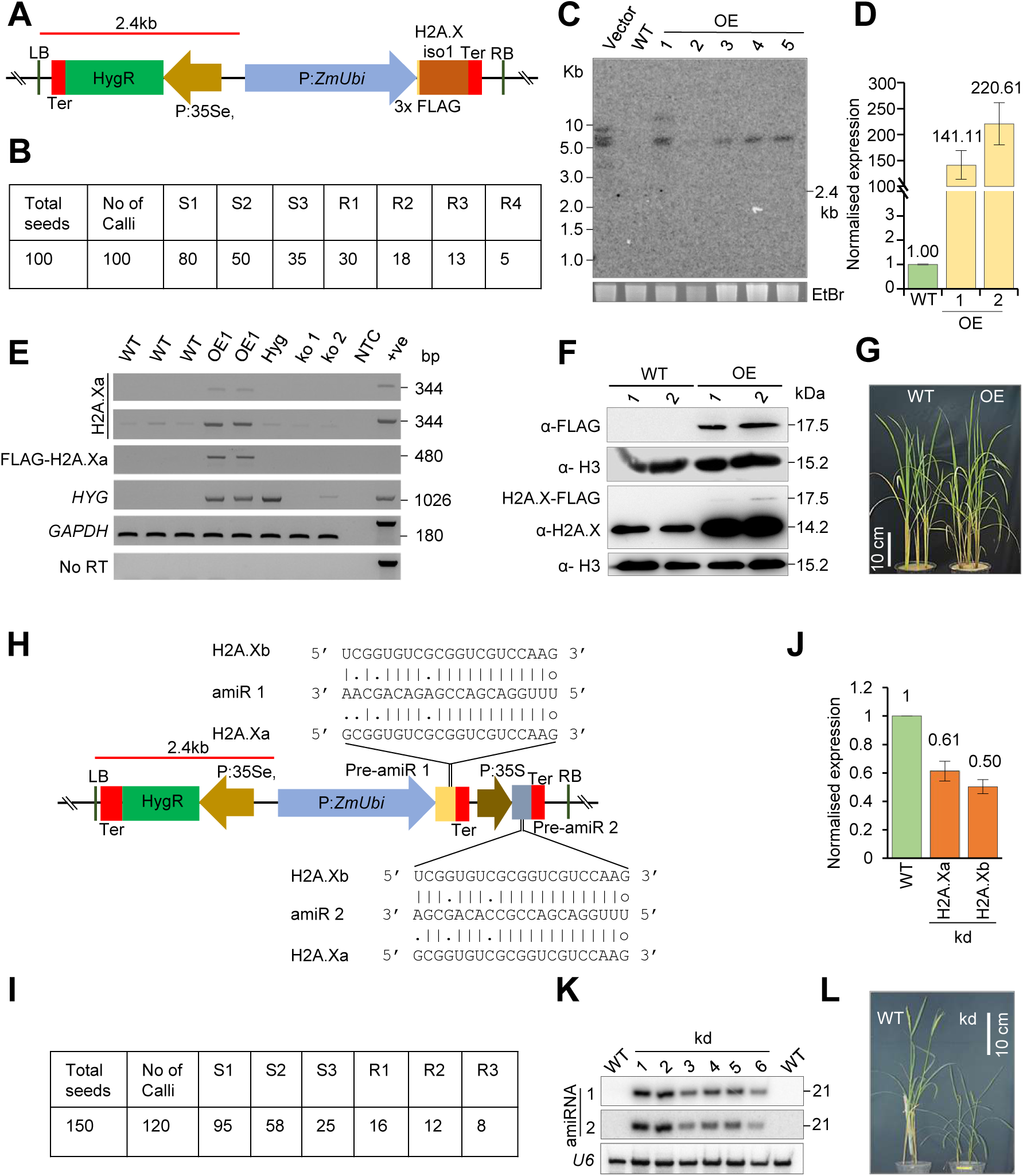
Generation and validation of H2A.X overexpression (OE) and knockdown (kd) lines. **A,** T-DNA map showing construct used to generate H2A.X OE with 3X-FLAG tag. **B,** Table showing summary of transformation to generate H2A.X OE plants (S-selection, R-Regeneration). **C,** Junction fragment Southern blot displaying integration pattern of T-DNA in OE plants. Minimum length of the DNA junction fragment (2.4 kb) is mentioned. EtBr-stained gel served as loading control. **D,** Bar plot showing expression of H2A.X in H2A.X OE lines. *OsGAPDH* served as internal control. qRT-PCR experiments are performed twice with consistent results. **E,** qRT-PCR (semi-quantitative) gel showing H2A.X OE along with controls. NTC-no template control. *OsGAPDH* was used as loading control. **F,** Western blot showing accumulation of FLAG tagged H2A.X. H3 served as the internal control. **G,** Image showing phenotypes of 45 d-old H2A.X OE lines. **H,** Vector map of amiR constructs used to target both the isoform of OsH2A.X. **I,** Table showing the summary of transformation to generate h2a.x knockdown (kd) lines. **J,** Bar plot showing expression of H2A.X in h2a.x kd lines. qRT-PCR experiments are performed twice with consistent results. **K,** Northern blots showing expression of amiR in h2a.x kd lines. **L,** Image showing phenotype of 60 d-old h2a.x kd lines.

**Supplementary Fig. S4.**
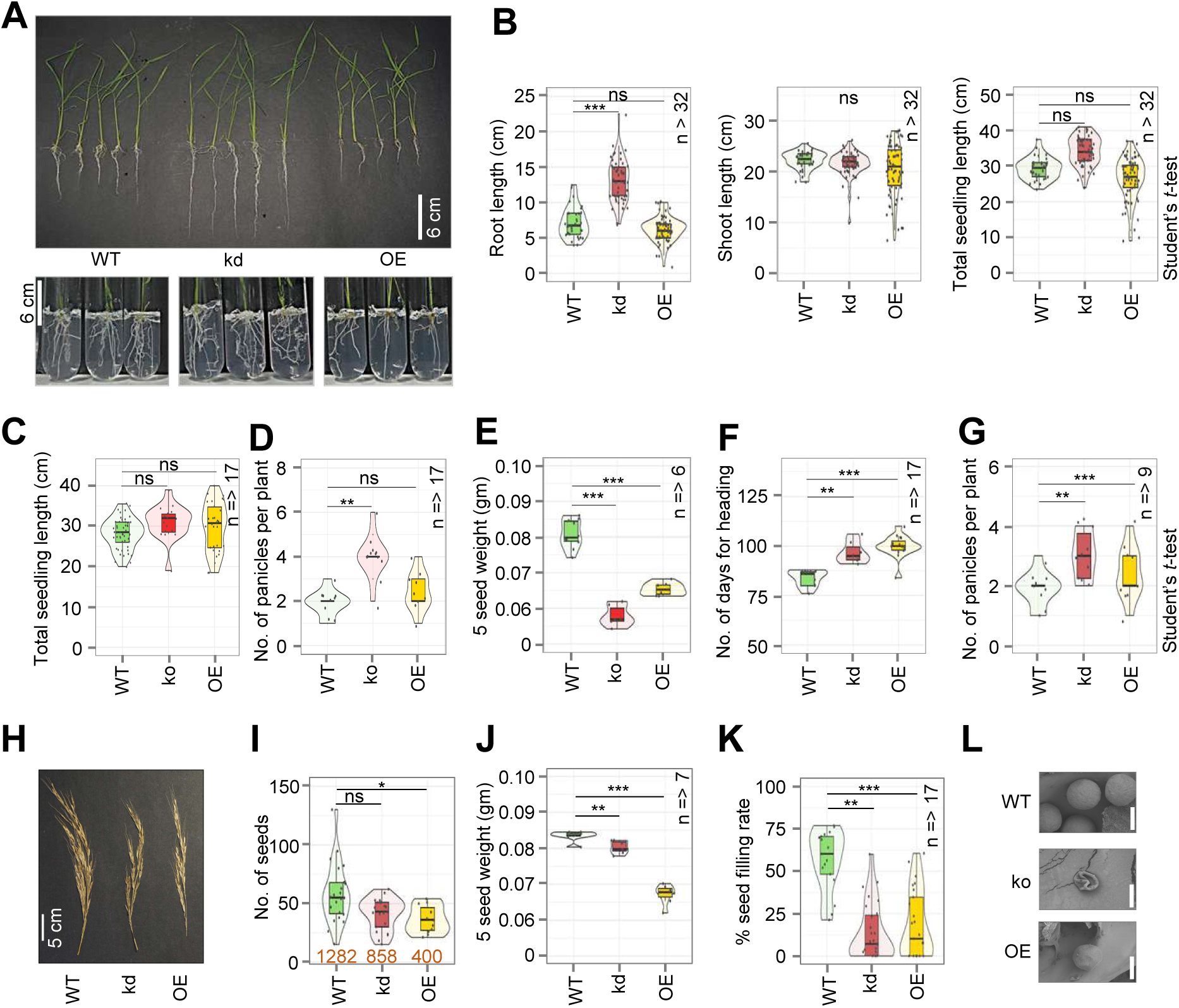
Phenotypes of H2A.X mis-expression lines. **A,** Morphology of 14-day-old seedling of H2A.X mis-expression lines. **B,** Box plots showing root length, shoot length and total seedling length of H2A.X mis-expression lines. **C,** Box plot showing height of H2A.X misexpression lines. **D,** Boxplot showing number of panicles in H2A.X mis-expression lines. Box plots showing 5 seed weight **E,** number of days for heading **F,** and number of panicles per plant **G,** in H2A.X misexpression lines. **H,** Image showing panicle phenotypes of H2A.X transgenic lines. **I,** Boxplot showing number of seeds obtained in H2A.X misexpression lines. **J,** Boxplots showing 5 seed weight, and **K,** percentage of filled seeds in H2A.X transgenics. **L,** SEM image showing morphology of pollens obtained from transgenic lines. Scale bar - 30 μm. Two-tailed Student’s *t*-test was used for statistical comparison in **B, C, D, E, F, G, I, J and K**. p-value (*) < 0.05, (**) < 0.01, (***) < 0.001, (ns) non-significant.

**Supplementary Fig. S5.**
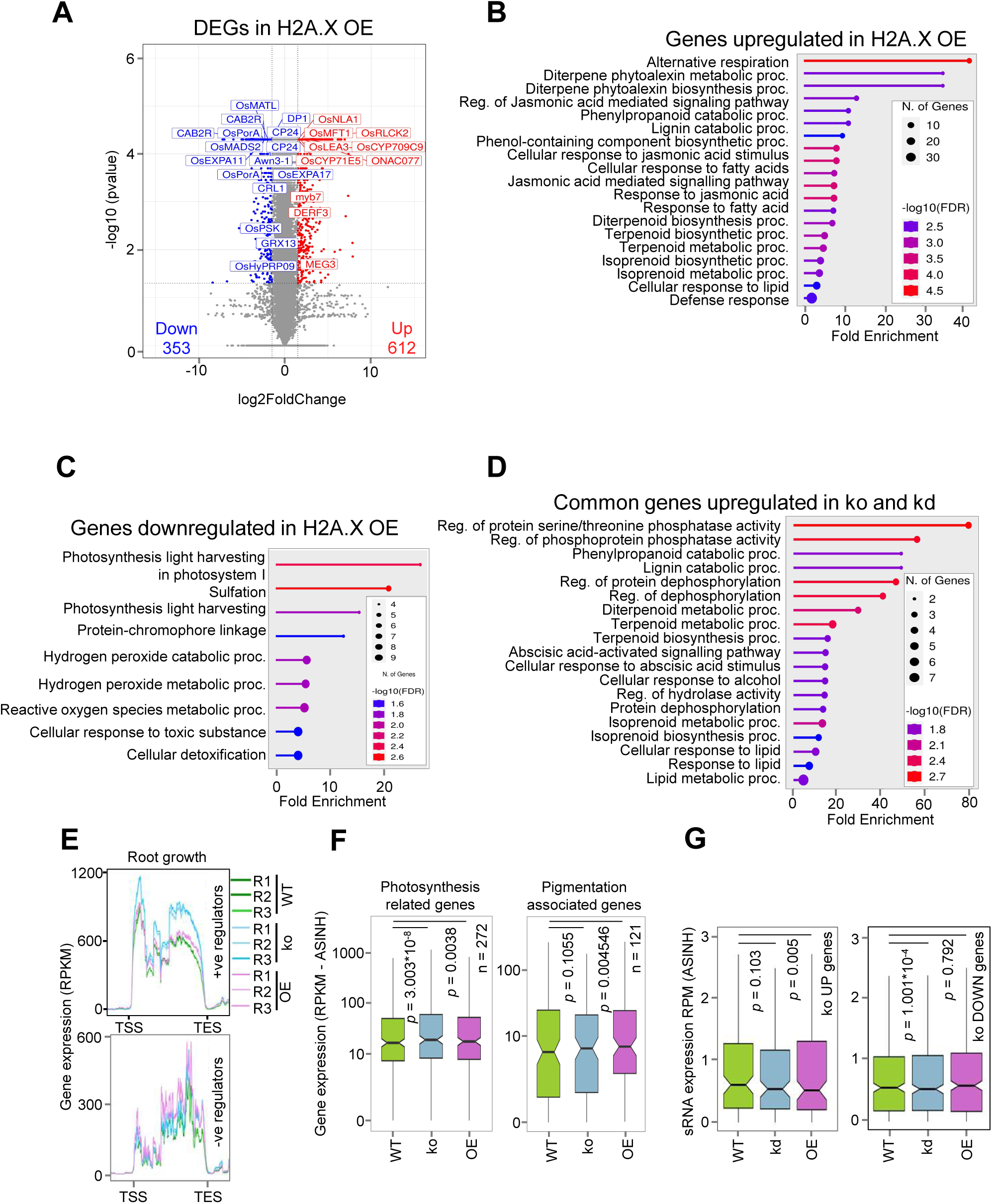
Transcriptome analysis of H2A.X mis-expression lines. **A,** Volcano plot representing all DEGs of H2A.X OE. **B,** GO enrichment categories of upregulated DEGs in OE. FDR-False discovery rate. **C,** GO enrichment categories of genes downregulated in H2A.X OE. **D,** GO enrichment categories of shared genes upregulated in ko and kd. **E,** Metaplots showing gene expression of positive (+ve) and negative (-ve) regulators of root growth in H2A.X transgenic lines. **F,** Boxplot showing expression of photosynthesis-associated and pigmentation-associated genes in transgenic lines. **G,** sRNA expression associated to DEGs in h2a.x ko plants. Two-sided Wilcoxon test (*P* < 0.01 was considered significant) in **F** and **G**.

**Supplementary Fig. S6.**
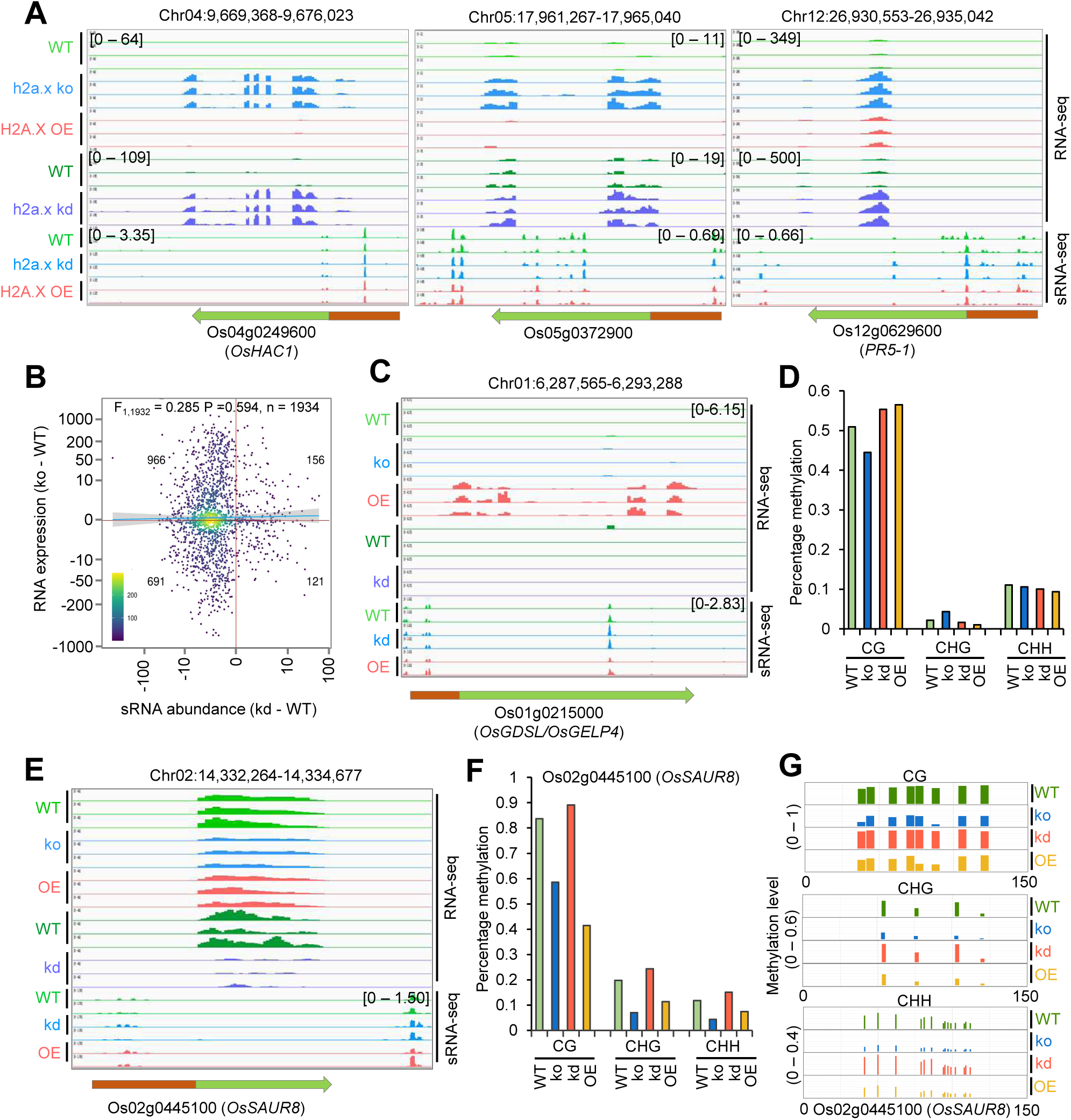
Targeted DNA methylation and sRNA analysis of genes mis-expressed in H2A.X transgenic lines. **A,** IGV screenshots showing common representative genes upregulated in h2a.x ko and kd lines. B, Correlation plot showing expression of sRNAs lost in h2a.x kd and mRNA expression (RPKM) from adjacent genes (RPKM) in h2a.x ko. C, IGV screenshot of representative gene *OsGDSL* that gained expression in H2A.X OE. D, Percentage methylation in different DNA methylation contexts in the promoter of *OsGDSL* in H2A.X mis-expression lines. E, IGV screenshot of OsSAUR8 with reduced gene expression in h2a.x ko and kd lines. F, Percentage methylation in different DNA methylation contexts in the promoter of *OsSAUR8* in H2A.X mis-expression lines. G, Diagram showing methylation coverage in different contexts in promoter of *OsSAUR8*.

**Supplementary Fig. S7.**
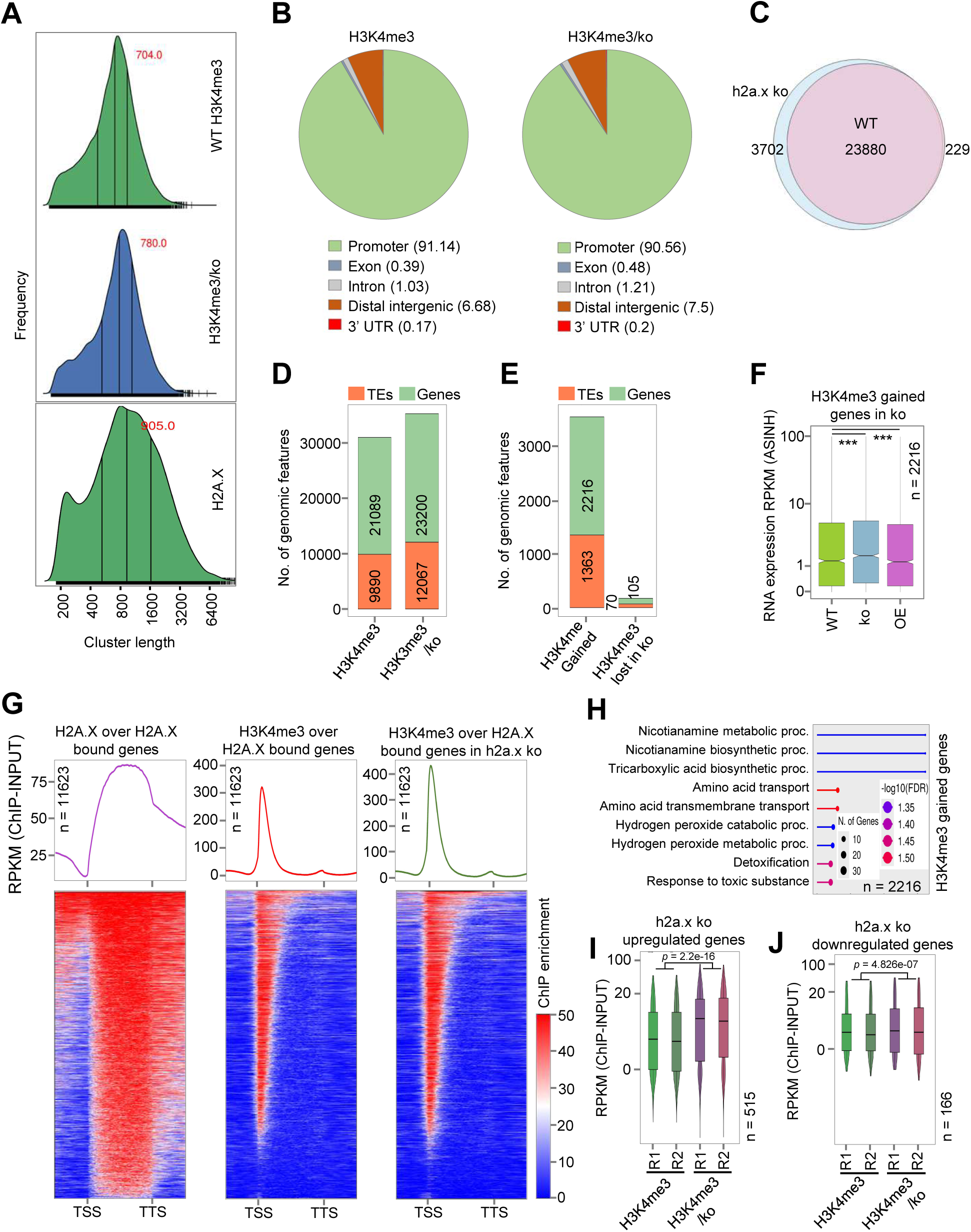
Gain of H3K4me3 was observed over genes in h2a.x ko lines. **A,** Density plot showing mean peak width of H3K4me3 in WT and h2a.x ko and H2A.X in WT. **B,** Vennpie chart showing the proportion of H3K4me3 bound genomic regions in WT and h2a.x ko plants. Percentage (%) of peaks occupied in specific genomic features is mentioned inside the bracket. **C,** Venn diagram showing overlap between the number of H3K4me3 peaks in WT and h2a.x ko. **D,** Barplot showing the number of TEs and genes bound by H3K4me3 in WT and h2a.x ko plants. **E,** Barplot showing the number of genomic features overlapping with H3K4me3 gained and lost regions. **F,** Boxplot showing expression of H3K4me3 gained genes in h2a.x ko. **G,** Heatmap showing enrichment of H2A.X and H3K4me3 over H2A.X bound genes in WT and h2a.x ko. **H,** GO enrichment categories of H3K4me3 gained genes in h2a.x ko. Boxplots showing H3K4me3 enrichment over upregulated **I,** and downregulated genes **J,** in h2a.x ko. Significance calculated by Two-sided Wilcoxon test (*p* < 0.01 was considered as significant).

**Supplementary Fig. S8.**
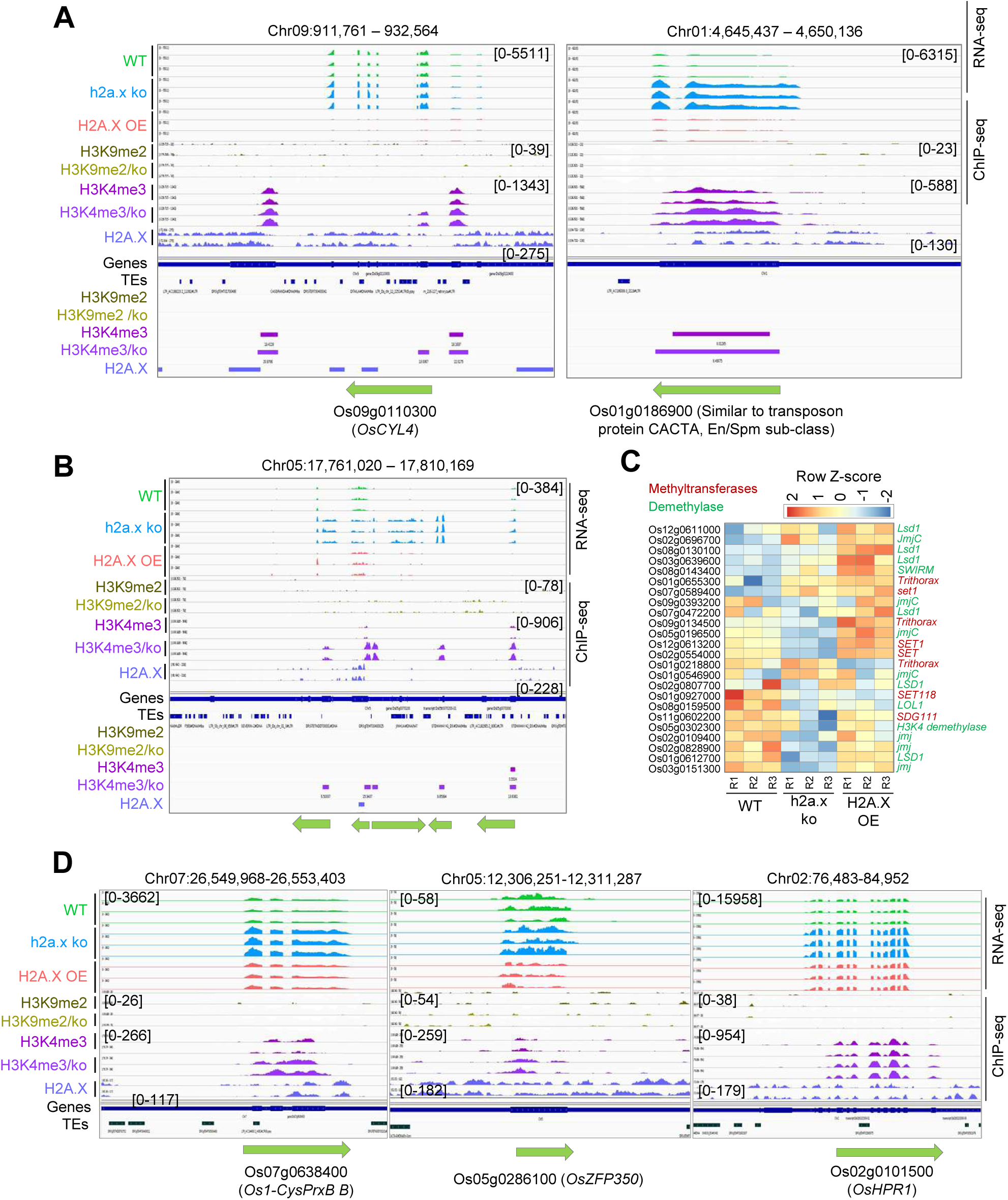
H3K4me3 gained genes exhibited increased expression in h2a.x ko lines. **A,** Representative IGV screenshots showing H3K4me3 gained genes which showed increased expression in h2a.x ko. **B,** IGV screenshot showing increased levels of H3K4me3 over genomic region and increased expression of the genes in that regions. **C,** Heatmap showing expression of histone methyltransferases and demethylases in H2A.X mis-expression lines. **D,** Representative IGV screenshots showing expression of development associated genes which gained H3K4me3 in H2A.X mis-expression lines.

**Supplementary Fig. S9.**
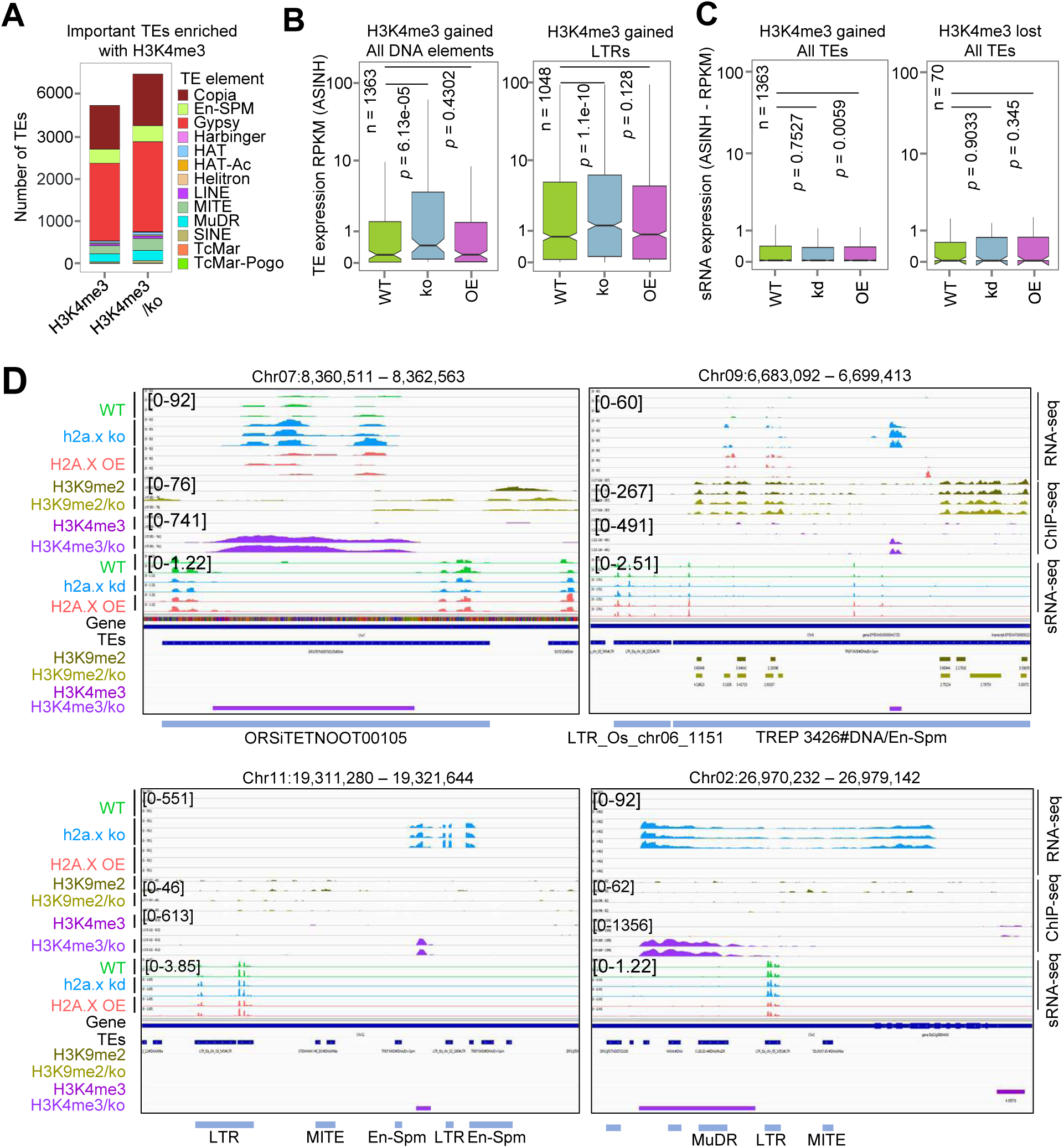
H3K4me3 gain in TEs led increased TE expression. **A,** Barplot showing number of TE features bound by H3K4me3 in WT and h2a.x ko. **B,** Boxplot showing the normalised RNA expression from H3K4me3 enriched all DNA elements and LTR elements. **C,** Boxplot showing expression of sRNA from H3K4me3 gained and lost TEs. **D,** IGV screenshots showing increased RNA expression from the H3K4me3 gained TEs and sRNAs derived from these regions in H2A.X mis-expression lines. Two-sided Wilcoxon test (*p* < 0.01 was considered as significant).

**Supplementary Fig. S10.**
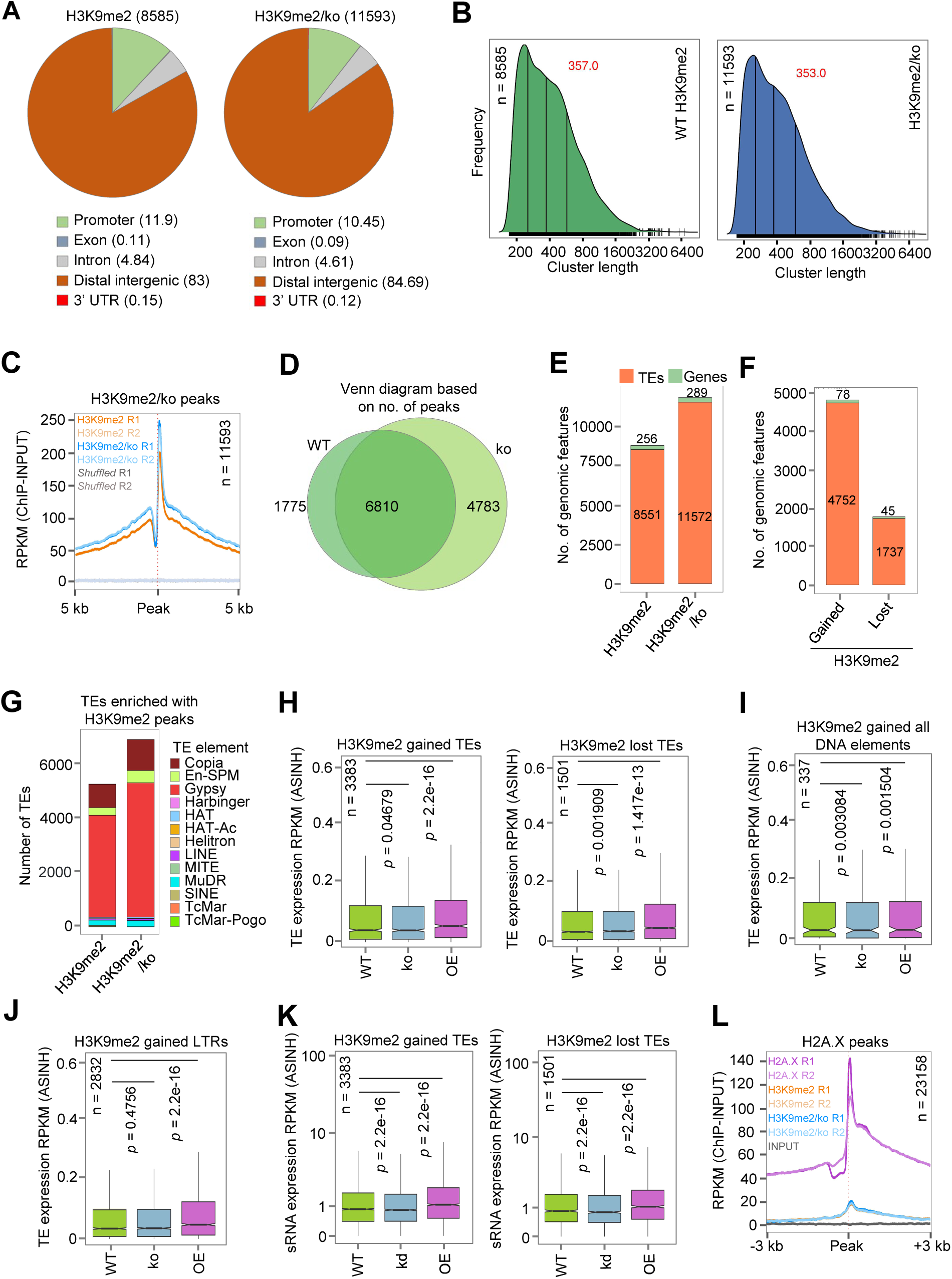
Gain of H3K9me2 was observed over TEs in h2a.x ko lines. **A,** Vennpie chart showing the proportion of genic and intergenic regions overlapping H3K9me2 marks in WT and h2a.x ko plants. Percentage (%) of peaks occupied in specific genomic features is mentioned inside the bracket. **B,** Density plot showing mean peak width of H3K9me2 in WT and h2a.x ko. **C,** Metagene plot showing enrichment of H3K9me2 in WT and h2a.x ko over H3K9me2 peaks in ko. **D,** Venn diagram showing overlap between peaks of H3K9me2 in WT and h2a.x ko. **E,** Barplot showing the number of TEs and genes bound by H3K9me2 in WT and h2a.x ko. **F,** Barplot showing the number of genes and TEs overlapping with H3K9me2 gained and lost regions. **G,** Barplot displaying number of TE features bound by H3K9me2 in WT and h2a.x ko. **H,** Boxplots showing TE expression from H3K9me2 gained and lost TEs. Boxplots showing RNA expression of all DNA elements **I,** and LTR elements **J,** which gained H3K9me2 in H2A.X mis-expression lines. **K,** Boxplot showing sRNA levels in H3K9me2 gained and lost TE elements in H2A.X mis-expression lines. **L,** Metagene plot showing enrichment of H2A.X and H3K9me2 over H2A.X ChIP peaks. Significance calculated by Two-sided Wilcoxon test (*p* < 0.01 was considered as significant).

**Supplementary Fig. S11.**
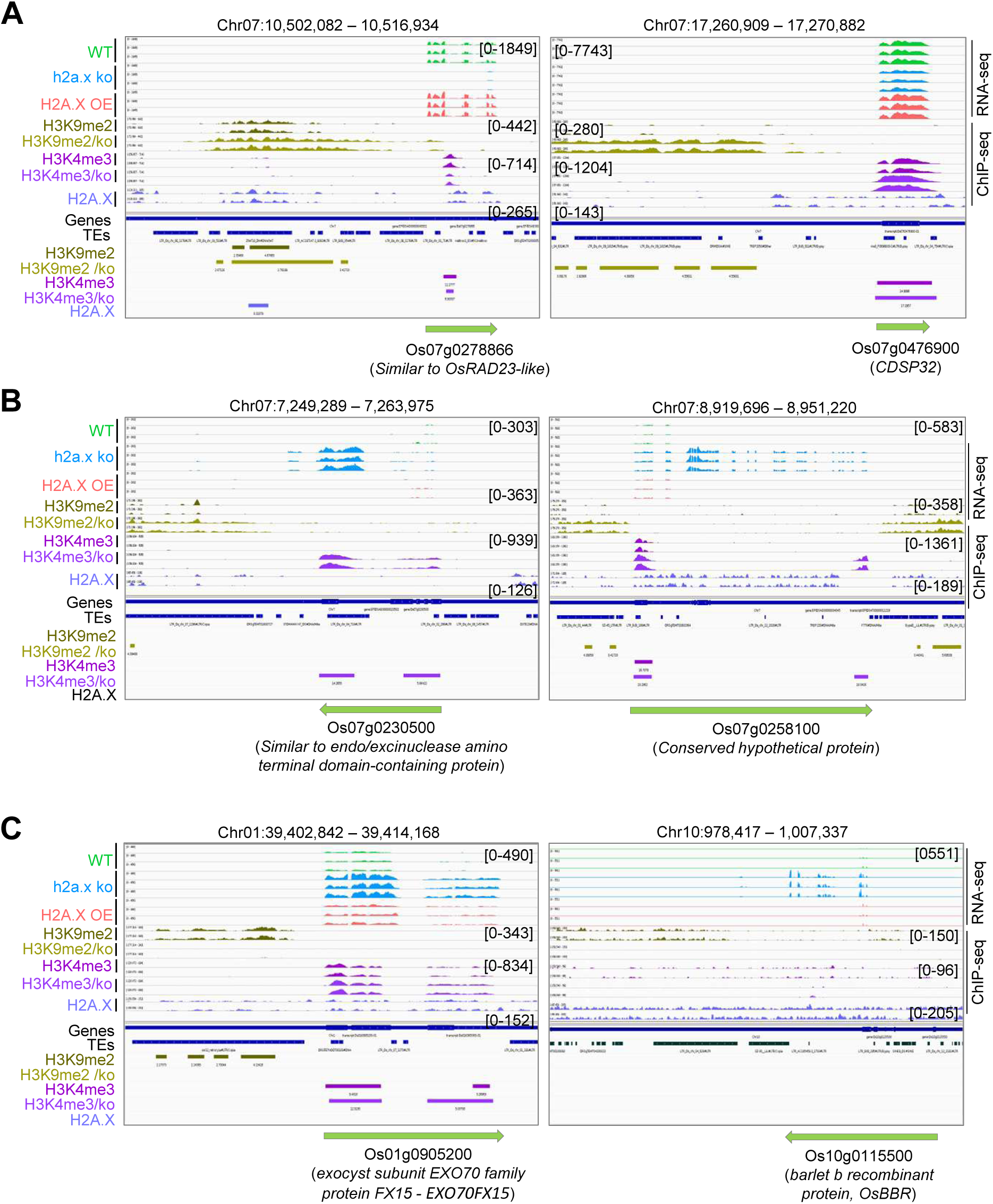
Altered H3K9me2 mark influenced gene expression in h2a.x ko. **A,** Representative IGV screenshots of genes showing increased H3K9me2 marks and reduced mRNA expression in h2a.x ko. **B,** Representative IGV screenshots of ko upregulated genes showing increased levels of both H3K9me2 and H3K4me3 marks. **C,** Representative IGV screenshots showing loss of H3K9me2 from candidate genes and their increased mRNA expression in h2a.x ko.

**Supplementary Fig. S12.**
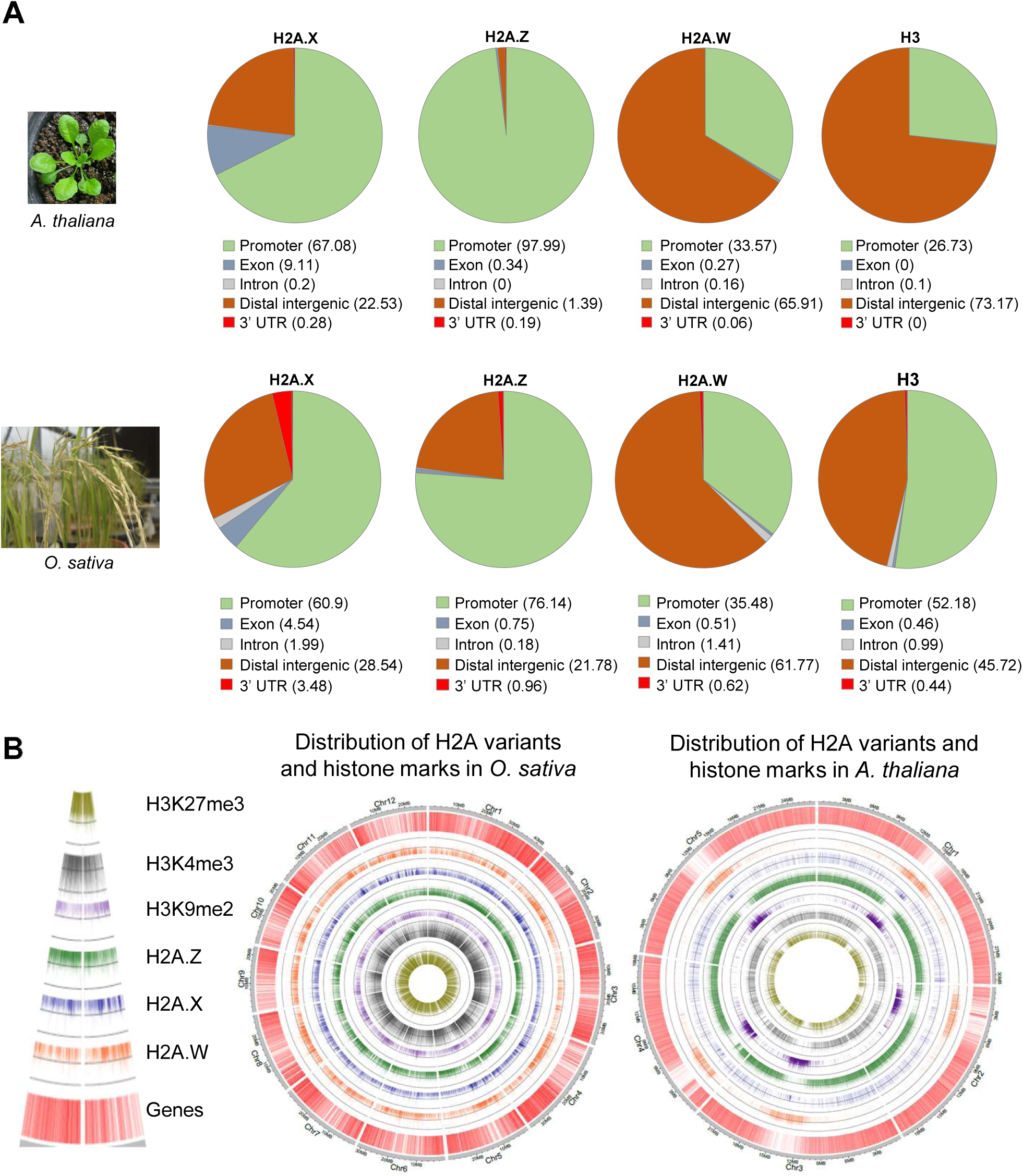
Genome-wide occupancy of histone variants and histone marks in *Arabidopsis* and *O. sativa*. **A,** Vennpie chart showing the genomic distribution of H2A variants in *A. thaliana* and *O. sativa.* Percentage (%) of peaks occupied by specific genomic features is mentioned inside the bracket. **B,** Circos plots showing the distribution of histone variants and histone marks in *A. thaliana* and *O. sativa*.

**Supplementary Fig. S13.**
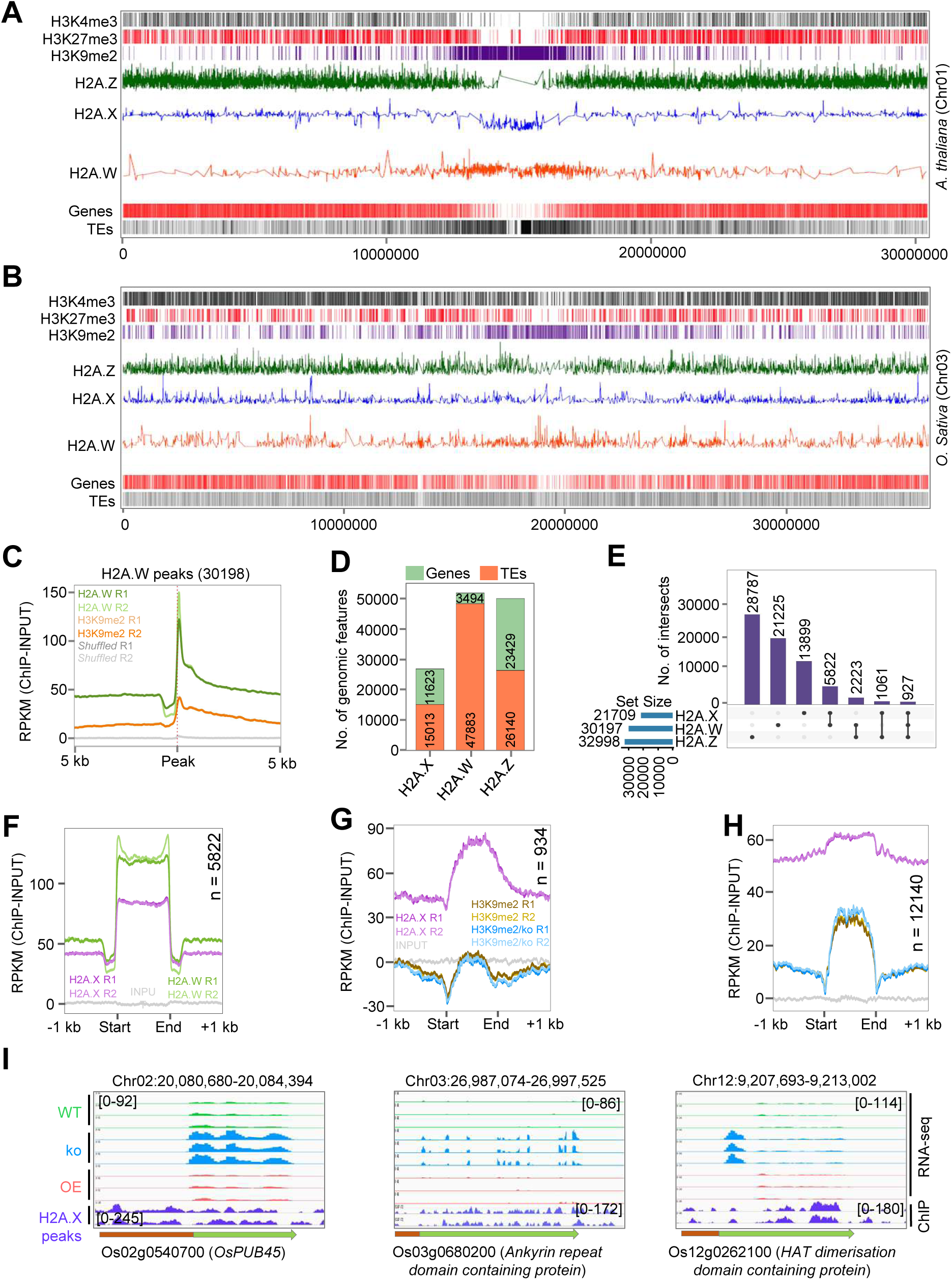
Histone variants occupy specific regions in the chromosome. Chromosome-wide map of histone marks and histone variant distribution in *A. thaliana* **A,** and *O. sativa* **B,. C,** Metagene plot showing enrichment of H2A.W ChIP signal over H2A.W peaks. **D,** Barplot showing the number of TEs and genes bound by H2A variants in WT plants. **E,** Upset plot showing overlap between H2A variant peaks in WT plants. **F,** Metagene plots showing enrichment of H2A.W and H2A.X ChIP signal over H2A.W and H2A.X common (5822) peaks. Enrichment of both H2A.X and H3K9me2 over common H2A.X and H2A.W gained genes **G**, and TEs **H,. I,** Representative IGV screenshot showing upregulation of genes bound by H2A.X in h2a.x ko plants.

**Supplementary Fig. S14.**
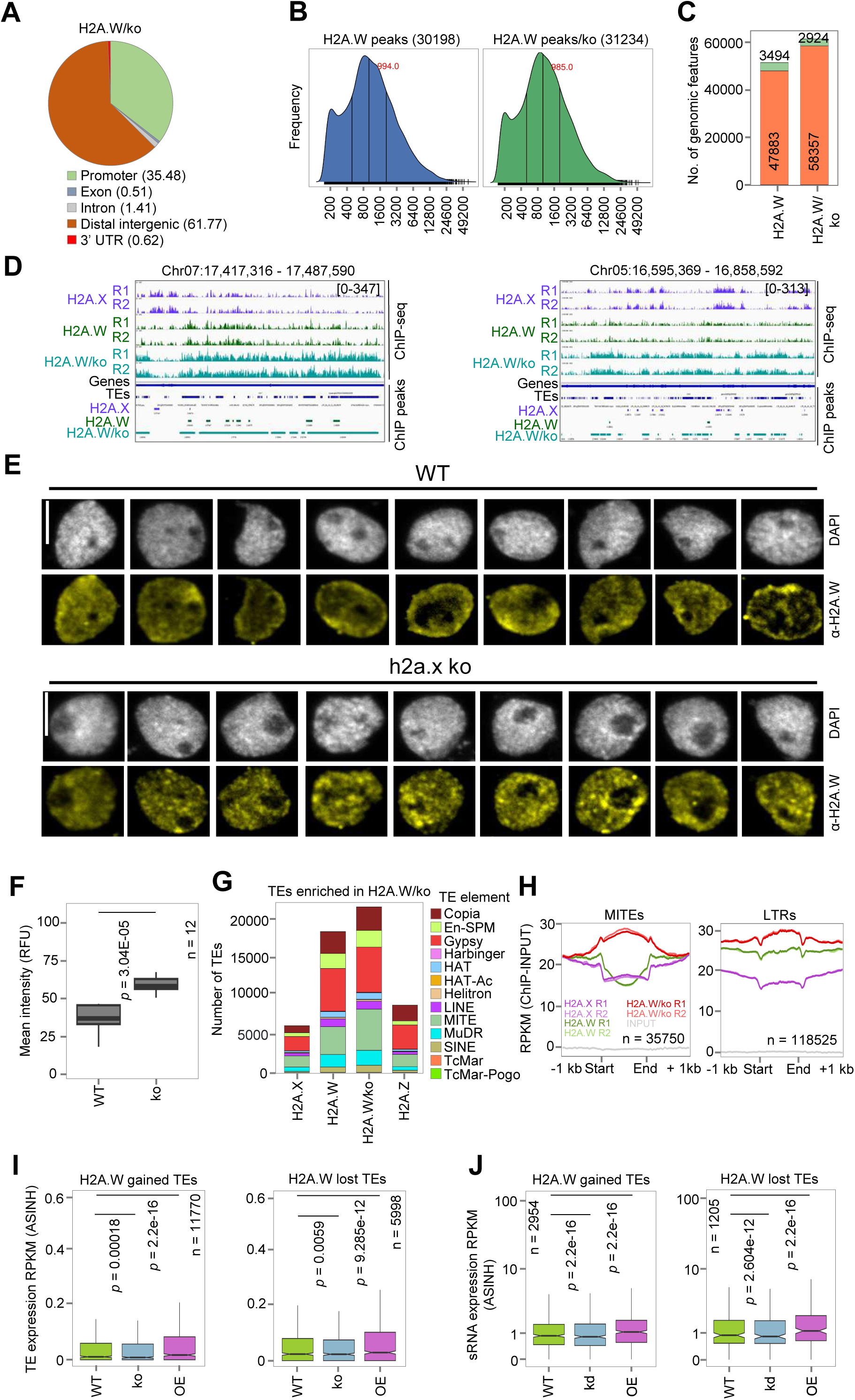
Distribution of H2A.W was altered in h2a.x ko. **A,** Vennpie chart showing the proportion of genic and intergenic regions overlapped by H2A.W in h2a.x ko plants. Percentage (%) of peaks occupied in specific genomic features is mentioned inside the bracket. **B,** Density plot showing mean peak width of H2A.W peaks in WT and h2a.x ko. **C,** Barplot showing number of TEs and genes bound by H2A.W in WT and h2a.x ko. **D,** IGV screenshot showing increased enrichment of H2A.W in h2a.x ko. **E,** IFL images of nuclei from WT and h2a.x ko plants stained using α-H2A.W. Scale: 5 μm. **F,** Boxplot showing mean intensity of relative fluorescence units (RFU) signal across nuclei in WT and h2a.x ko. **G,** Boxplot showing number of TEs bound by H2A variants. **H,** Metagene plot showing enrichment of H2A.W over MITEs and LTRs. **I,** Boxplot showing RNA expression from TEs in H2A.W gained and lost regions. **J,** Boxplot displaying sRNA levels in H2A.W gained and lost TE elements in H2A.X mis-expression lines. Significance calculated by Two-sided Wilcoxon test (*p* < 0.01 was considered significant).

**Supplementary Fig. S15.**
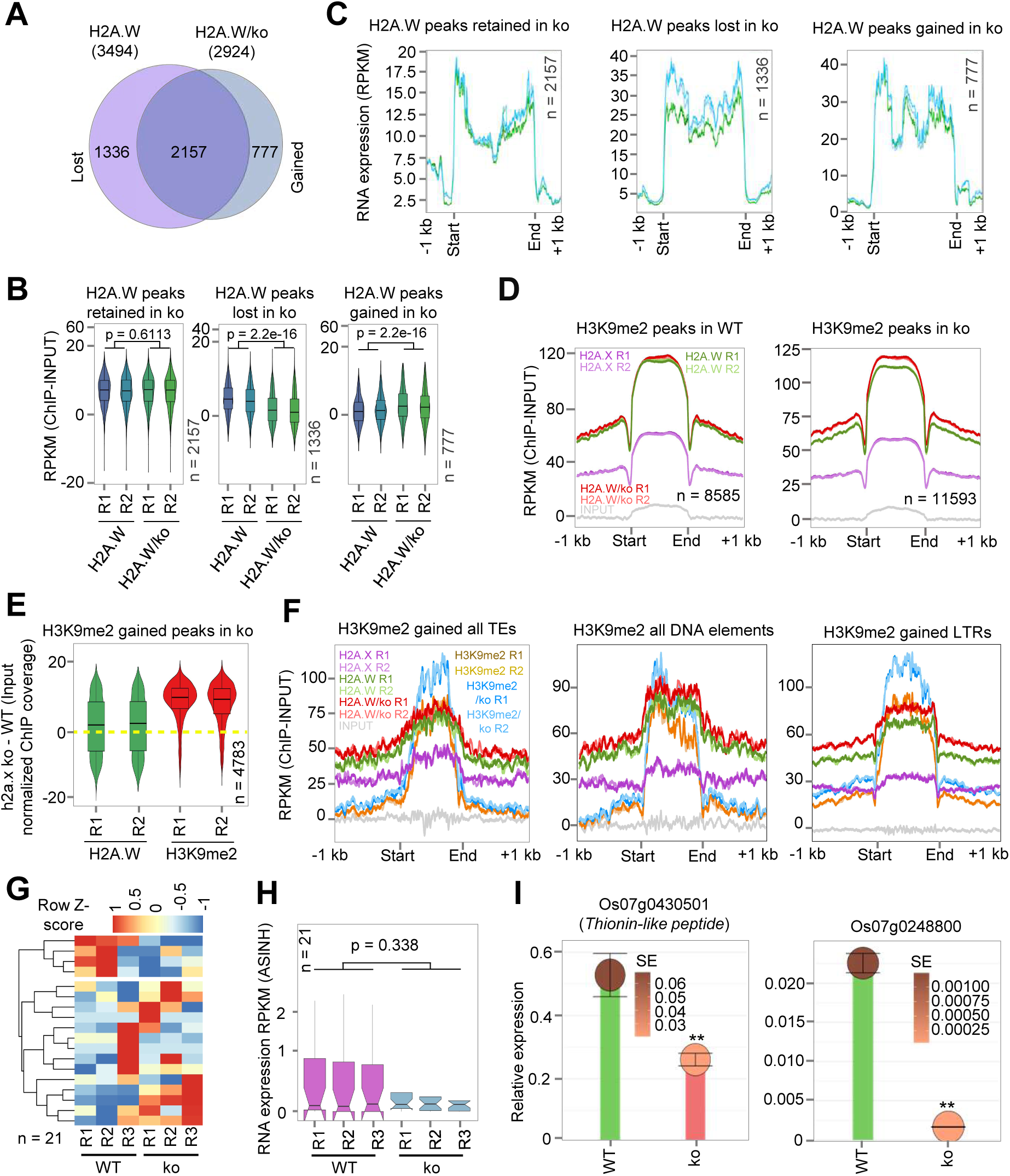
Altered H2A.W occupancy in h2a.x ko had minimal influence in gene expression. **A,** Venn diagram showing number of genes bound by H2A.W in h2a.x ko compared to WT. **B,** Boxplot showing enrichment of H2A.W over H2A.W altered genes in h2a.x ko. **C,** Metagene plot showing gene expression in H2A.W altered genes in h2a.x ko. **D,** Metagene plot showing enrichment of H2A.W and H2A.X over H3K9me2 peaks in WT and h2ax ko. **E,** Boxplot showing the enrichment of H2A.W and H3K9me2 in h2a.x ko over H3K9me2 gained regions. **F,** Metagene plot showing enrichment of H2A.X, H2A.W and H3K9me2 over all TE elements, all DNA elements and LTRs which gained H3K9me2. **G,** Heatmap showing expression of both H2A.W and H3K9me2 gained genes in h2a.x ko. **H,** Boxplot showing expression of genes gained both H2A.W and H3K9me2 in h2a.x ko. **I,** qRT-PCR showing the expression of representative genes Os07g0430501 and Os07g0248800 that gained both H2A.W and H3K9me2 in h2a.x ko. *OsGAPDH* served as internal control and two-tailed Student’s *t*-test was used for statistical comparison. Data represents means ± SE, n = 3. qRT-PCR experiments are performed twice with consistent results. Significance calculated by Two-sided Wilcoxon test (*p* < 0.01 was considered as significant) for **B, H and I**.

**Supplementary Fig. S16.**
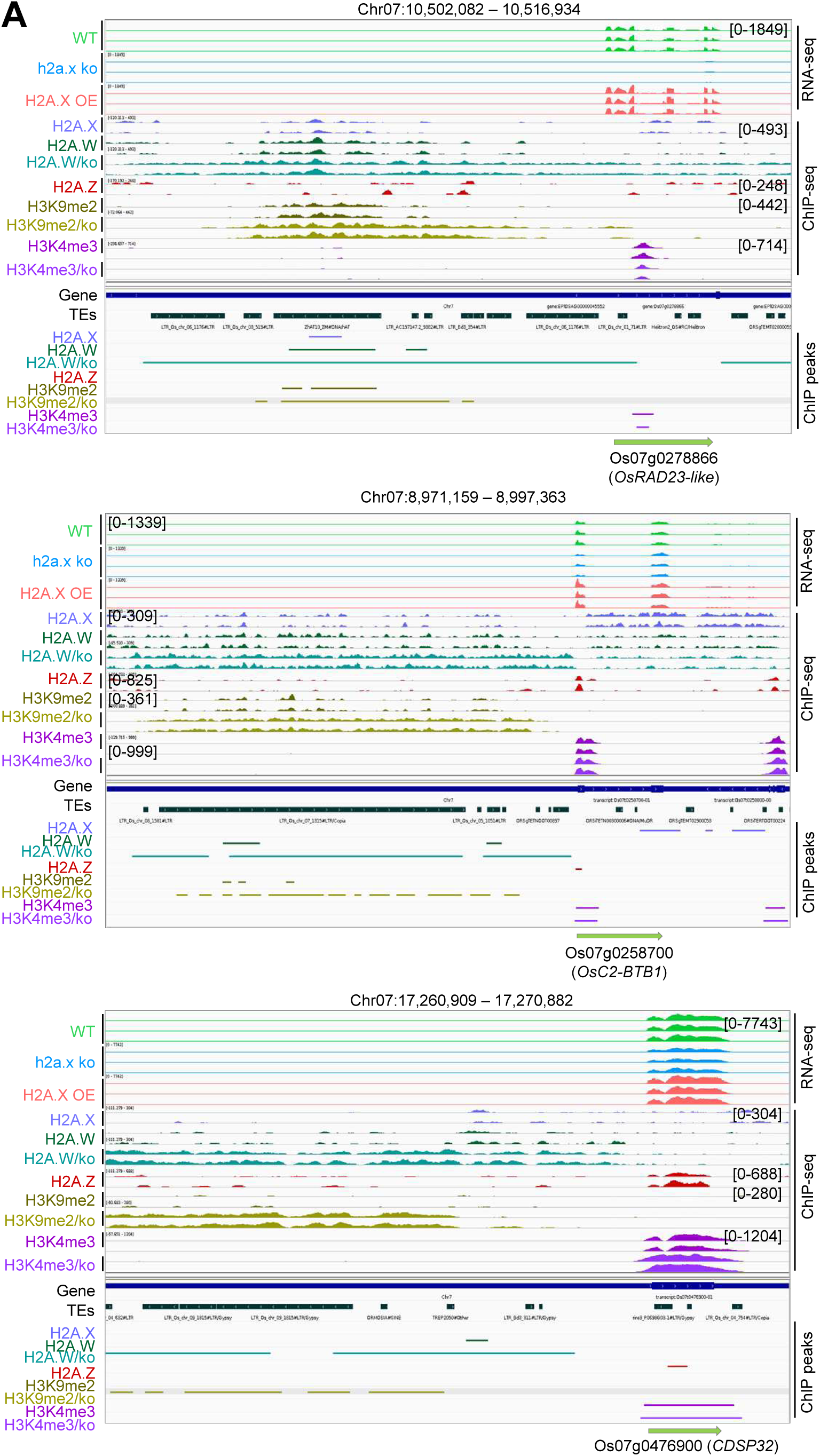
Genes that gained H3K9me2 had increased levels of H2A.W. **A,** Representative IGV screenshots showing genomic regions that gained H2A.W and H3K9me2 and show reduced expression of proximal genes.

**Supplementary Fig. S17.**
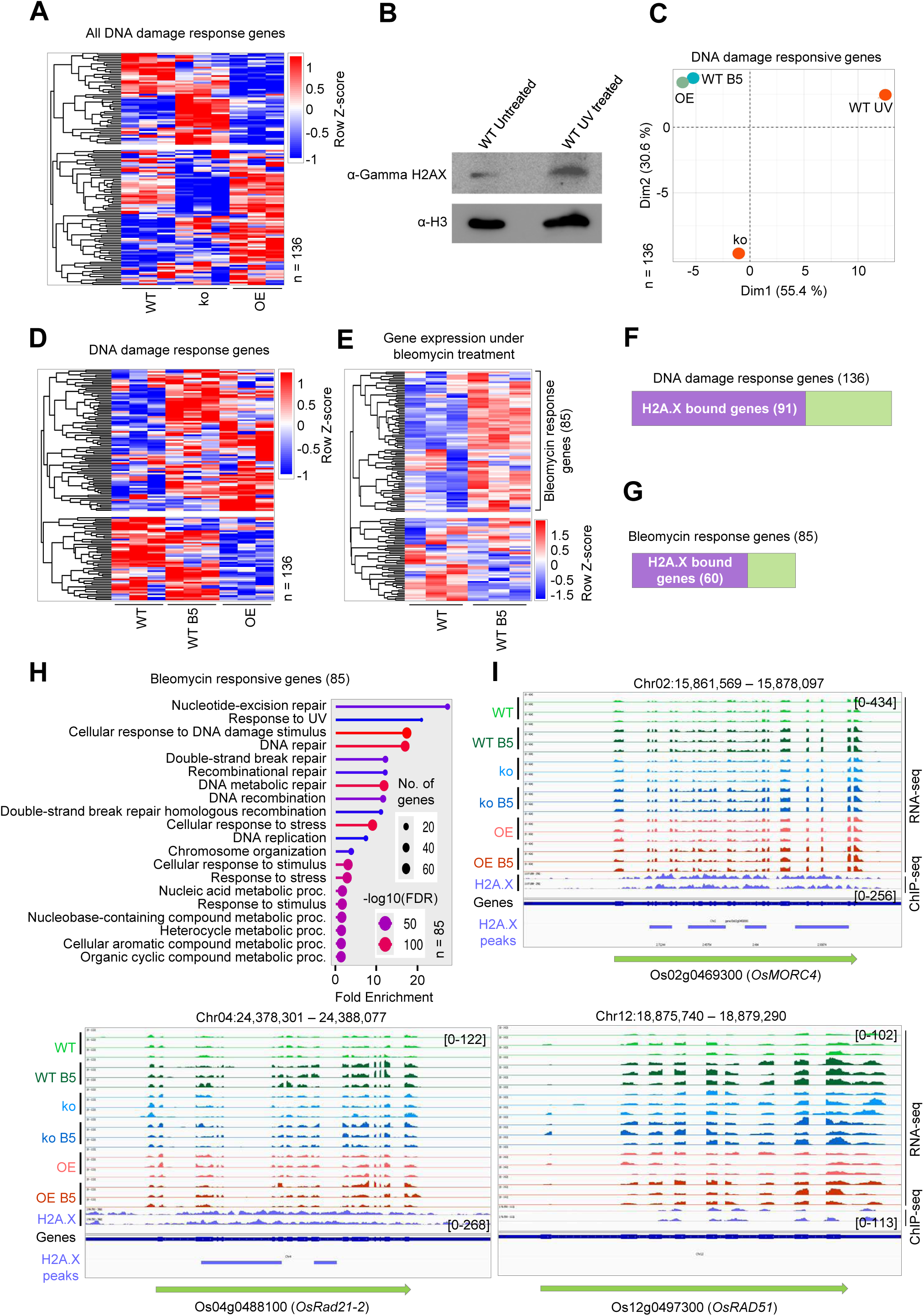
H2A.X regulates DNA damage response in rice. **A,** Heatmap showing expression of DNA damage responsive genes in H2A.X mis-expression lines. **B,** Western blot of phospho-H2A.X under UV treatment (3KJ). **C,** PCA analysis of DNA damage responsive genes in WT bleomycin treatment (WT B5), UV treatment and among H2A.X mis-expression lines. **D,** Heatmap showing expression of DNA damage genes in WT B5 and H2A.X OE plants. **E,** Heatmap showing expression of DNA damage responsive genes in WT B5. **F,** Plot showing number of DNA damage responsive genes bound by H2A.X. **G,** Plot showing number of bleomycin responsive genes bound by H2A.X. **H,** GO enrichment categories of bleomycin responsive genes. **I,** IGV screenshots showing expression of well-known DNA damage genes in H2A.X misexpression lines.

**Supplementary Fig. S18.**
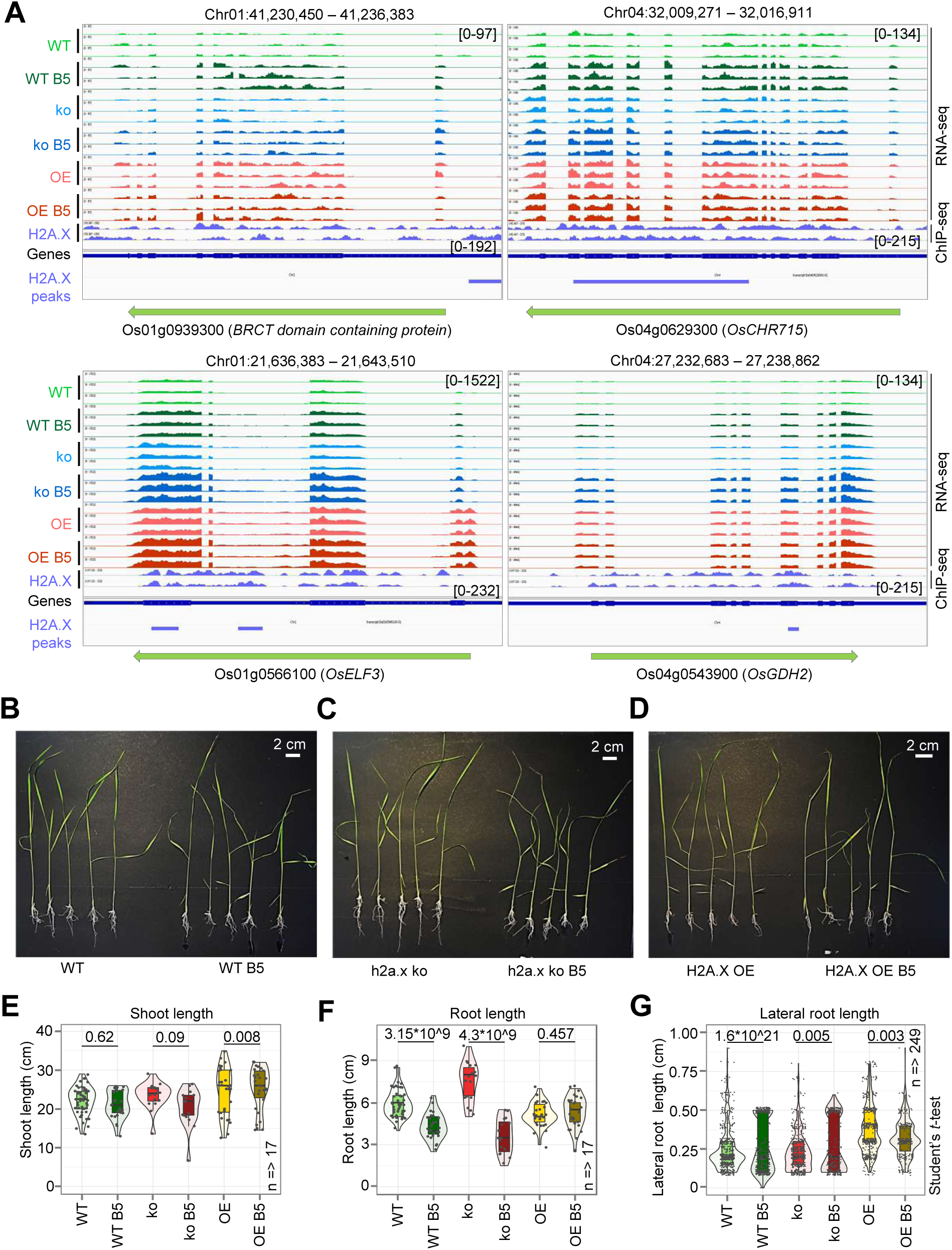
H2A.X OE mimicked DNA damage without any stress. **A,** Representative IGV screenshots showing expression of well-known stress responsive genes which are upregulated in H2A.X OE. Seedling phenotypes of WT **B,** h2a.x ko **C,** and H2A.X OE **D,** under bleomycin treatment. Boxplot showing shoot length **E,** root length **F,** and lateral root length **G,** among H2A.X misexpression lines after B5 treatment. Two-tailed Student’s *t*-test was used for statistical comparison.

**Supplementary Fig. S19.**
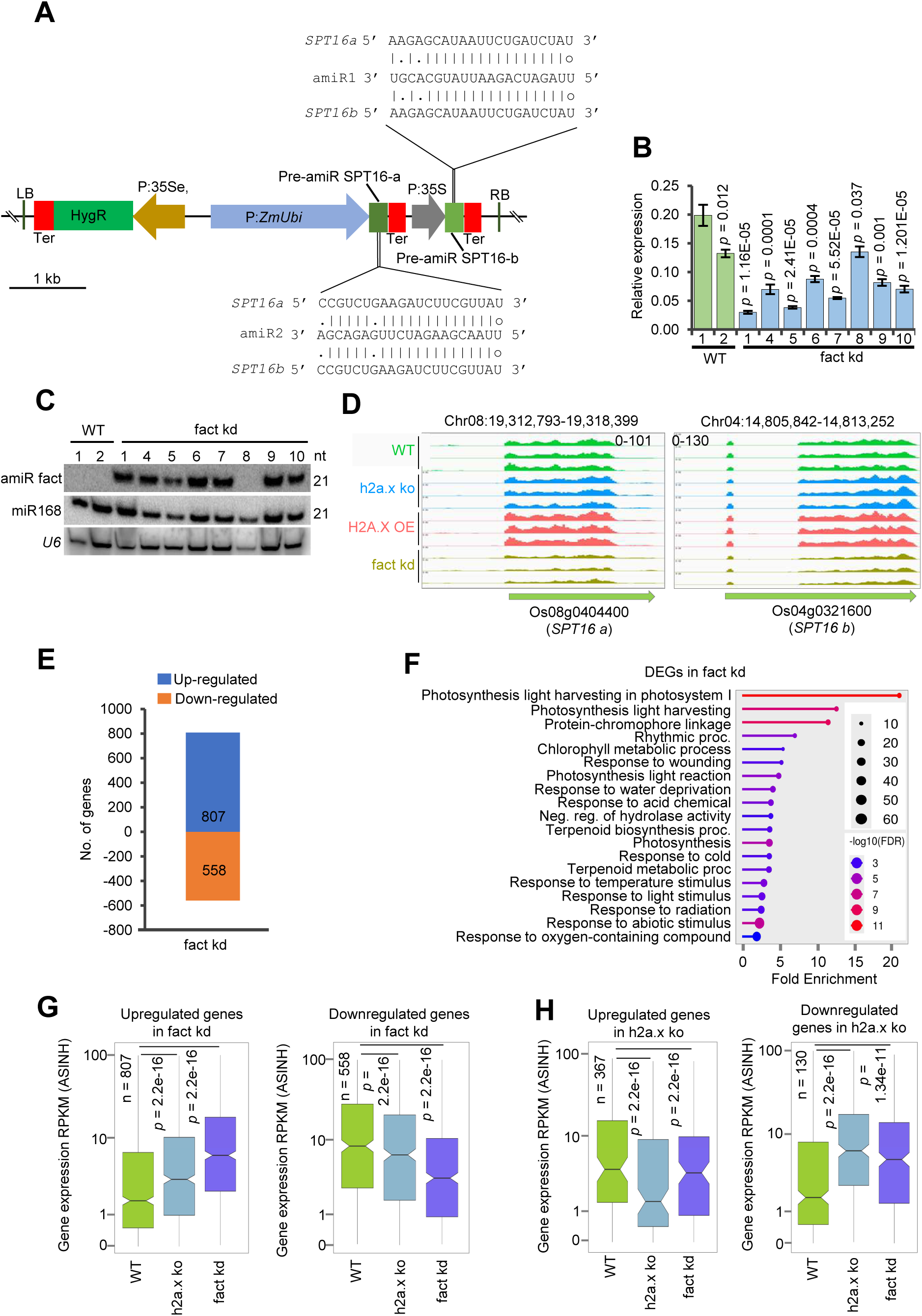
fact kd and h2a.x ko exhibited similar transcriptome. **A,** Vector map of amiR construct to target both the isoform of *OsSPT16*. **B,** Boxplot showing expression of *SPT16* isoforms in fact kd lines. Data represents means ± SE, n = 3. **C,** sRNA northern blot depicting the expression of amiR in fact kd lines **D,** IGV screenshot showing the reduced expression of *OsSPT16a* and *OsSPT16b* in fact kd lines. **E,** Barplot showing the summary of mis-expressed genes in fact kd. **F,** Gene ontology showing the category of genes altered in fact kd lines. **G,** Boxplots showing the expression of fact kd upregulated and downregulated genes in h2a.x ko and fact kd lines. **H,** Boxplots showing the expression of h2a.x ko upregulated and downregulated genes in h2a.x ko and fact kd lines. Significance calculated by Two-sided Wilcoxon test (*p* < 0.01 was considered significant) for **B, G** and **H**.

**Supplementary Fig. S20.**
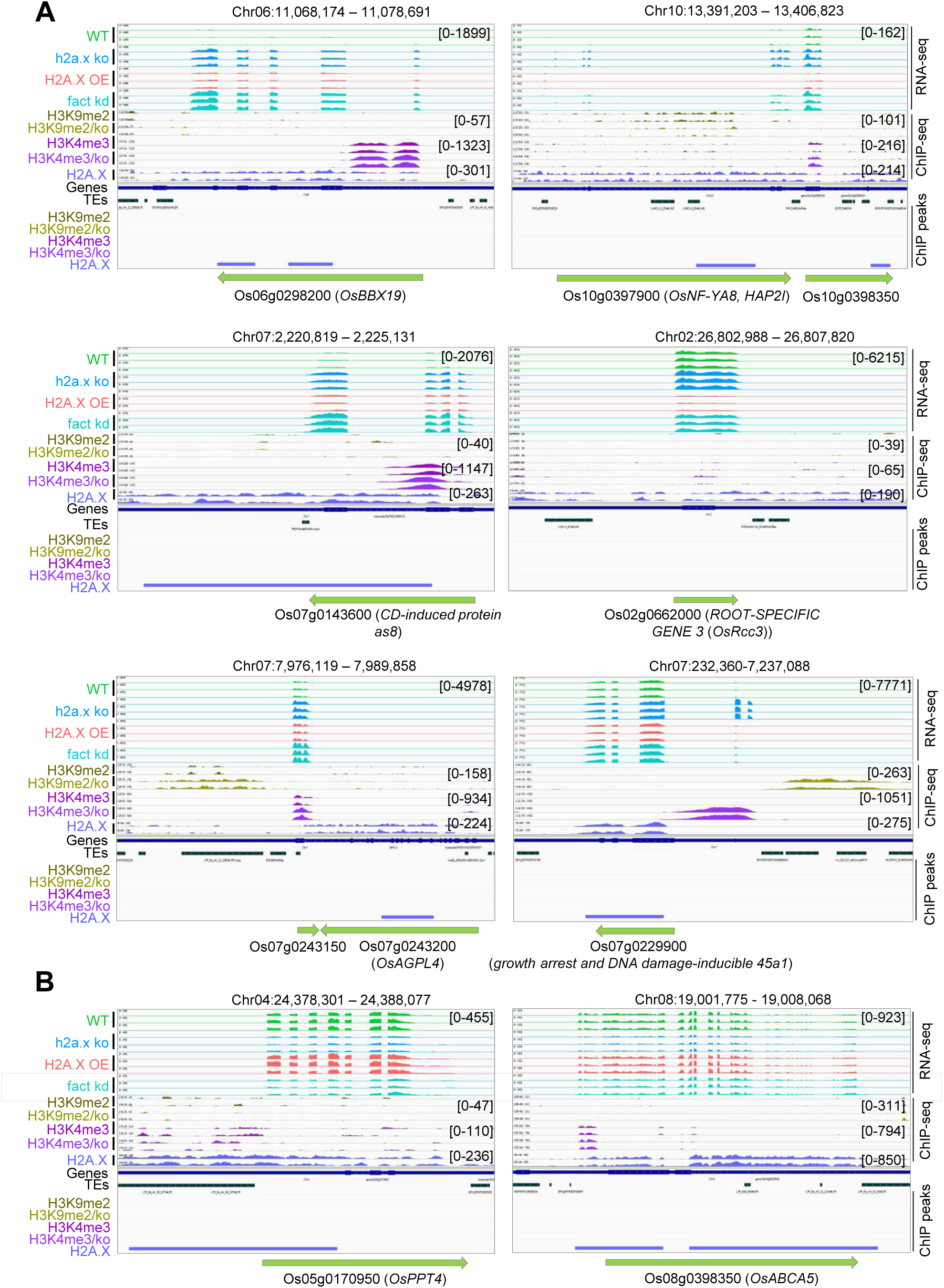
Development-associated genes exhibited similar pattern of expression in fact kd and h2a.x ko plants. **A,** Representative IGV screenshot showing upregulation of H2A.X bound genes in h2a.x ko and fact kd lines. **B,** Representative IGV screenshot showing downregulation of H2A.X bound genes in h2a.x ko and fact kd lines.

### Supplementary Tables S1 - S4

**Supplementary Table S1:**
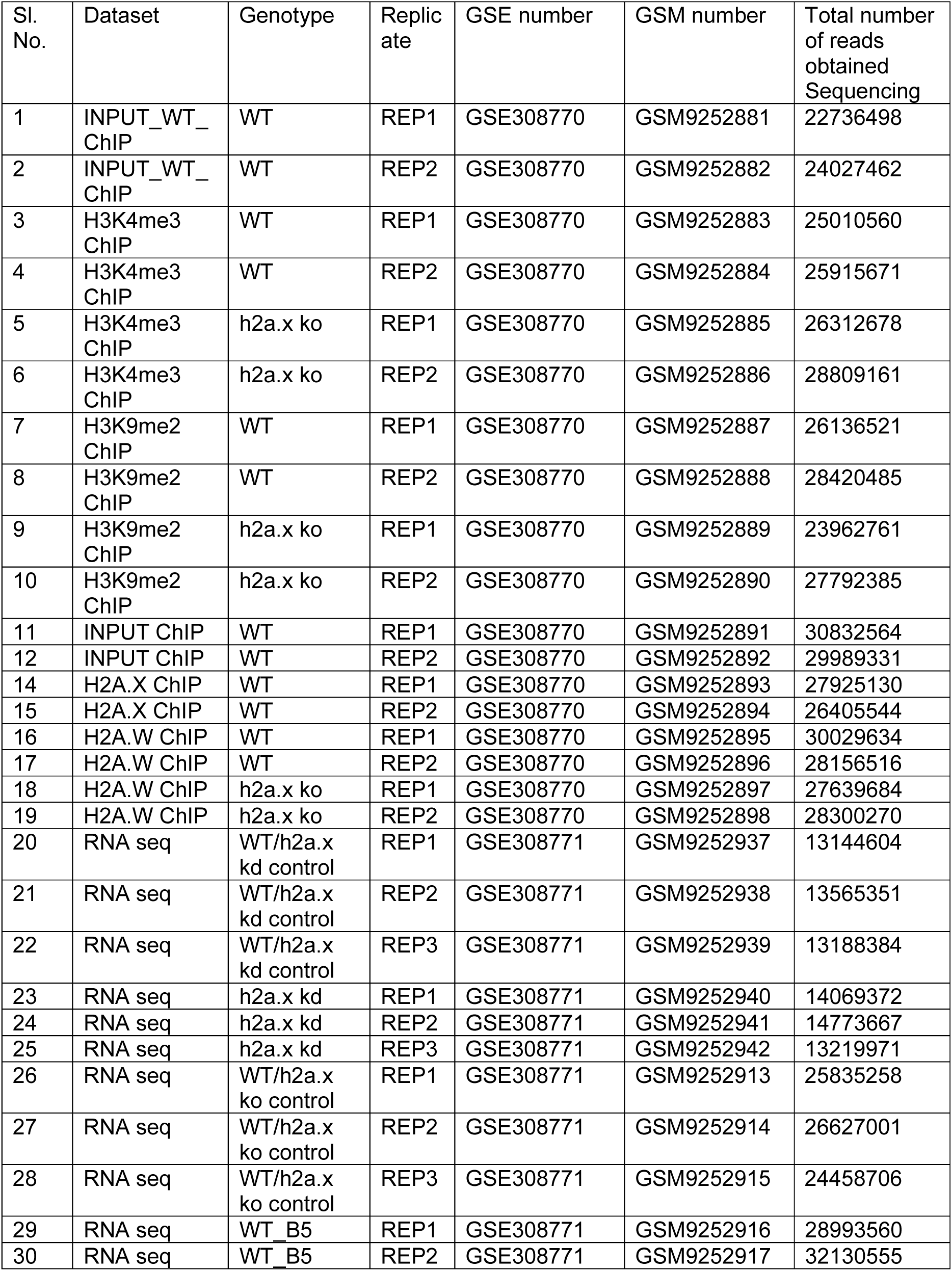
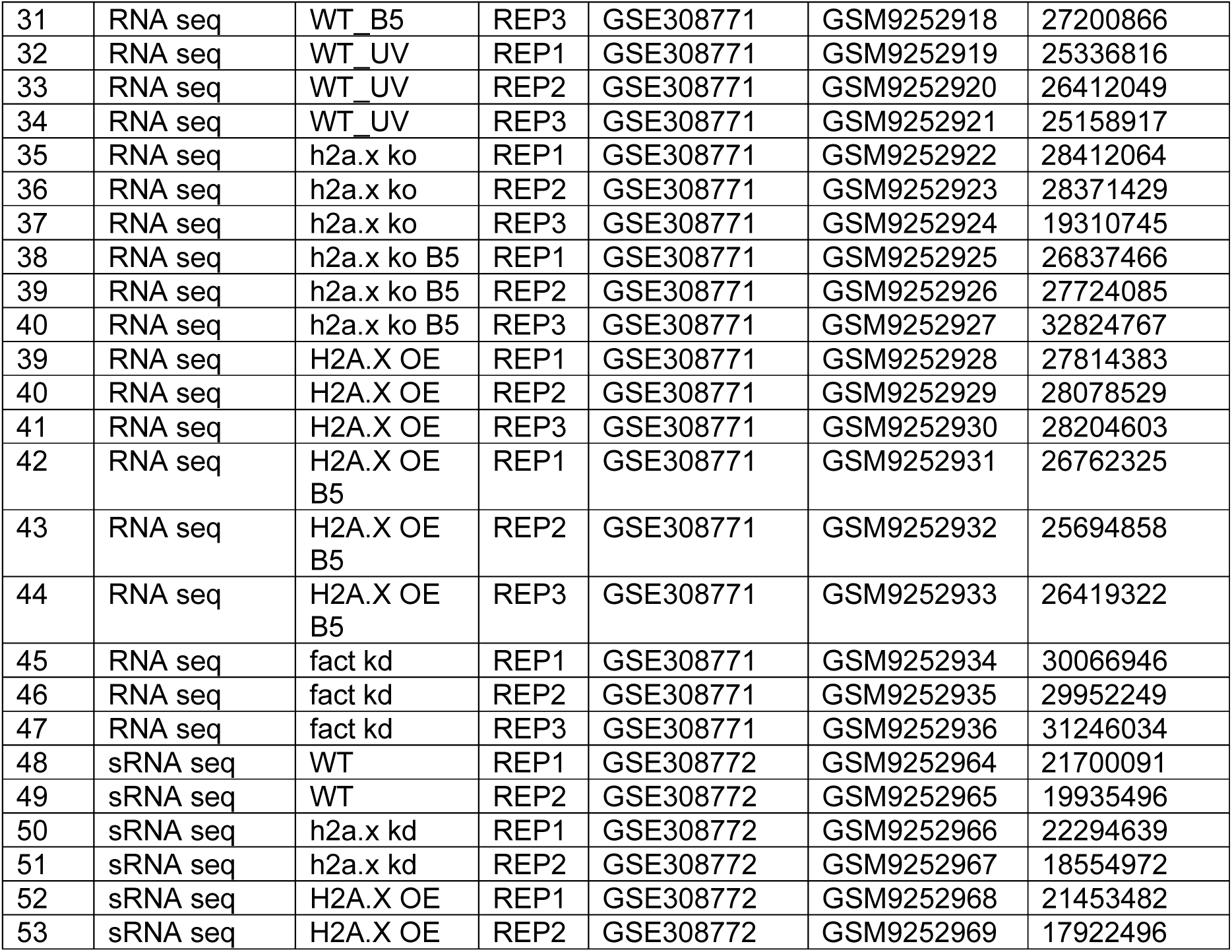
Details of high-throughput genomics data generated in this study.

**Supplementary Table S2:** Details of high-throughput genomics data obtained from publicly available datasets.

| Sl. No. | Species | Dataset type | Source tissue | SRA number | GSE number | Reference |
| --- | --- | --- | --- | --- | --- | --- |
| 1 | *Arabidopsis* | OsH2A rep1 ChIP seq | 10-day old seedling | SRR113048 32 | GSE1469 48 | (Bourguet et al., 2021) |
| 2 | *Arabidopsis* | OsH2A rep2 ChIP seq | 10-day old seedling | SRR113048 33 | GSE1469 48 | (Bourguet et al., 2021) |
| 3 | *Arabidopsis* | OsH2A.X rep1 ChIP seq | 10-day old seedling | SRR113048 34 | GSE1469 48 | (Bourguet et al., 2021) |
| 4 | *Arabidopsis* | OsH2A.X rep2 ChIP seq | 10-day old seedling | SRR113048 35 | GSE1469 48 | (Bourguet et al., 2021) |
| 5 | *Arabidopsis* | OsH2A.Z rep1 ChIP seq | 10-day old seedling | SRR113048 36 | GSE1469 48 | (Bourguet et al., 2021) |
| 6 | *Arabidopsis* | OsH2A.Z rep2 ChIP seq | 10-day old seedling | SRR113048 37 | GSE1469 48 | (Bourguet et al., 2021) |
| 7 | *Arabidopsis* | OsH3 rep1 ChIP seq | 10-day old seedling | SRR11304838 | GSE146948 | (Bourguet et al., 2021) |
| 8 | *Arabidopsis* | OsH3 rep2 ChIP seq | 10-day old seedling | SRR11304839 | GSE146948 | (Bourguet et al., 2021) |
| 9 | *Arabidopsis* | INPUT rep1 ChIP seq | 10-day old seedling | SRR11304846 | GSE146948 | (Bourguet et al., 2021) |
| 10 | *Arabidopsis* | INPUT rep2 ChIP seq | 10-day old seedling | SRR11304847 | GSE146948 | (Bourguet et al., 2021) |
| 11 | *Arabidopsis* | OsH2AW rep2 ChIP seq | 3-week-old *Arabidopsis* seedlings | SRR988544 | GSE50942 | (Yelagandula et al., 2014) |
| 12 | *Arabidopsis* | H3K4me3 ChIP seq | 10-day old seedling | SRR23686114 | GSE226469 | (Jamge et al., 2023a) |
| 13 | *Arabidopsis* | H3K27me3 ChIP seq | 10-day old seedling | SRR23686113 | GSE226469 | (Jamge et al., 2023a) |
| 14 | *Arabidopsis* | H3K9me2 ChIP seq | 10-day old seedling | SRR23686116 | GSE226469 | (Jamge et al., 2023a) |
| 15 | Rice | H2A.Z rep1 ChIP seq | 30-day old rice seedling | SRR12888663 | GSE155265 | (Du et al., 2020) |
| 16 | Rice | H2A.Z rep2 ChIP seq | 30-day old rice seedling | SRR12888664 | GSE155265 | (Du et al., 2020) |
| 17 | Rice | H3 rep1 ChIP seq | 30-day old rice seedling | SRR12888665 | GSE155265 | (Du et al., 2020) |
| 18 | Rice | H3 rep2 ChIP seq | 30-day old rice seedling | SRR12888666 | GSE155265 | (Du et al., 2020) |
| 19 | Rice | Input rep1 ChIP seq | 30-day old rice seedling | SRR12888667 | GSE155265 | (Du et al., 2020) |
| 20 | Rice | Input rep2 ChIP seq | 30-day old rice seedling | SRR12888668 | GSE155265 | (Du et al., 2020) |
| 21 | Rice | H3K27me3 rep1 ChIP seq | 12-day old rice seedling | SRR2142303 | GSE71640 | (Zhou et al., 2016) |
| 22 | Rice | H3K27me3 rep2 ChIP seq | 12-day old rice seedling | SRR4004040 | GSE71640 | (Zhou et al., 2016) |

**Supplementary Table S3:**
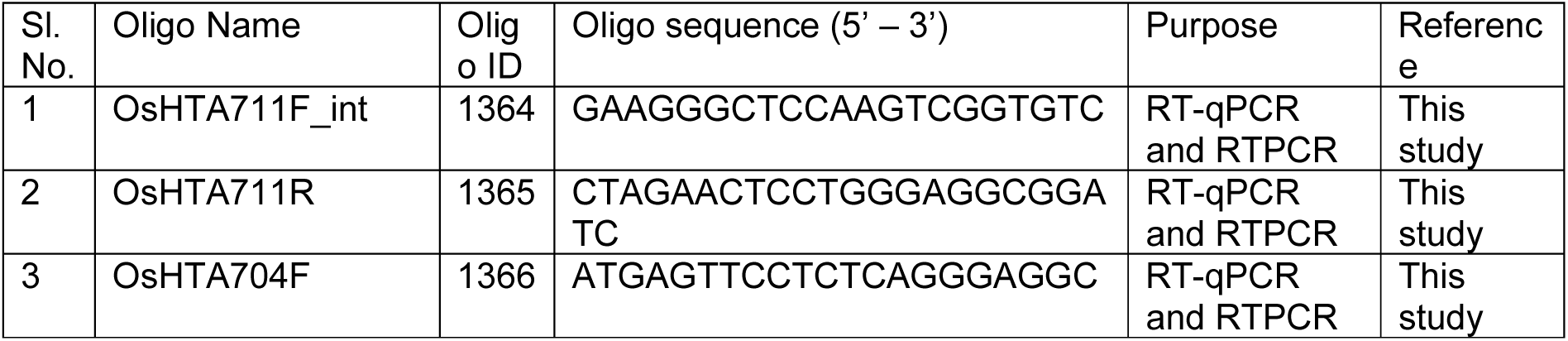
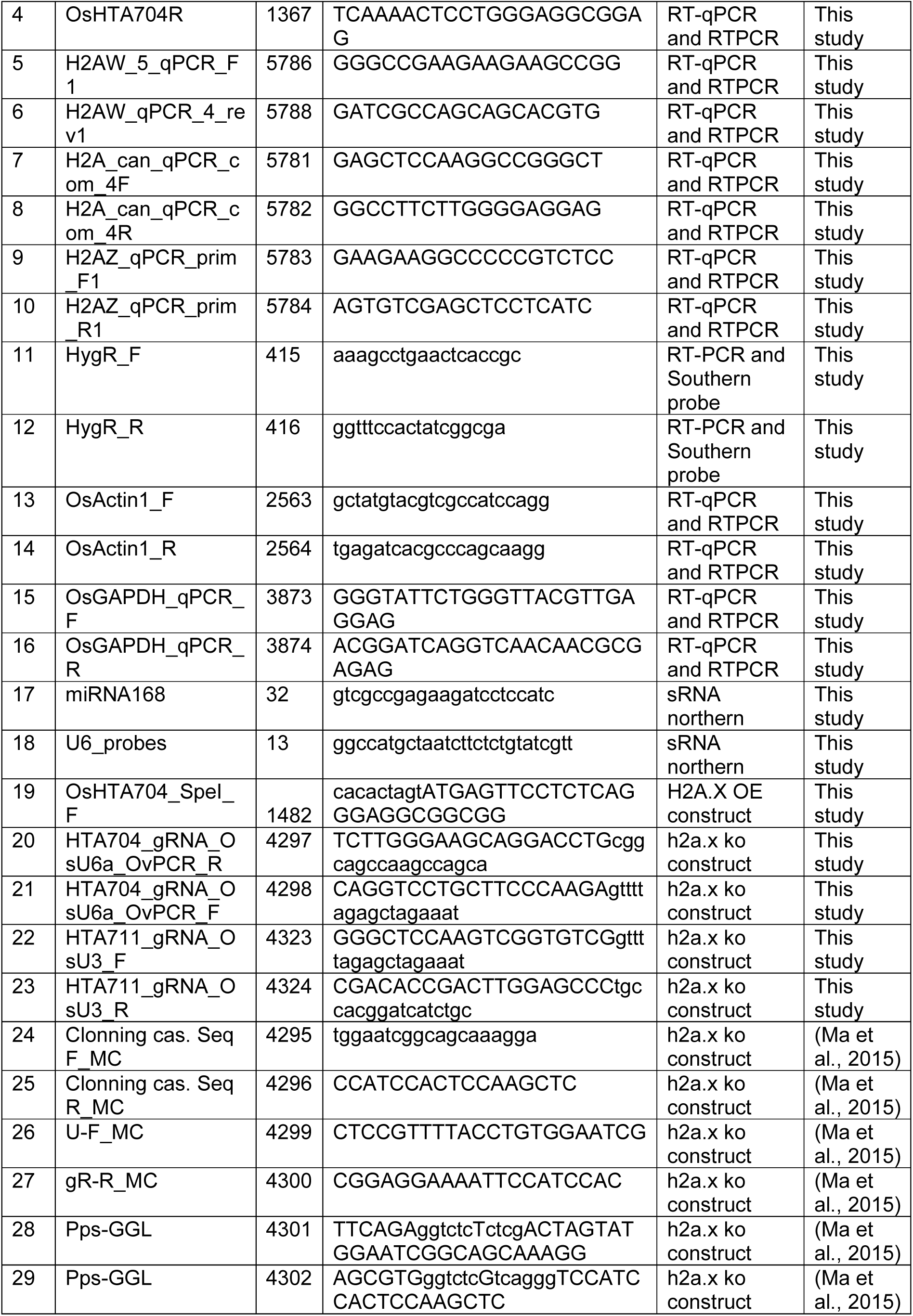
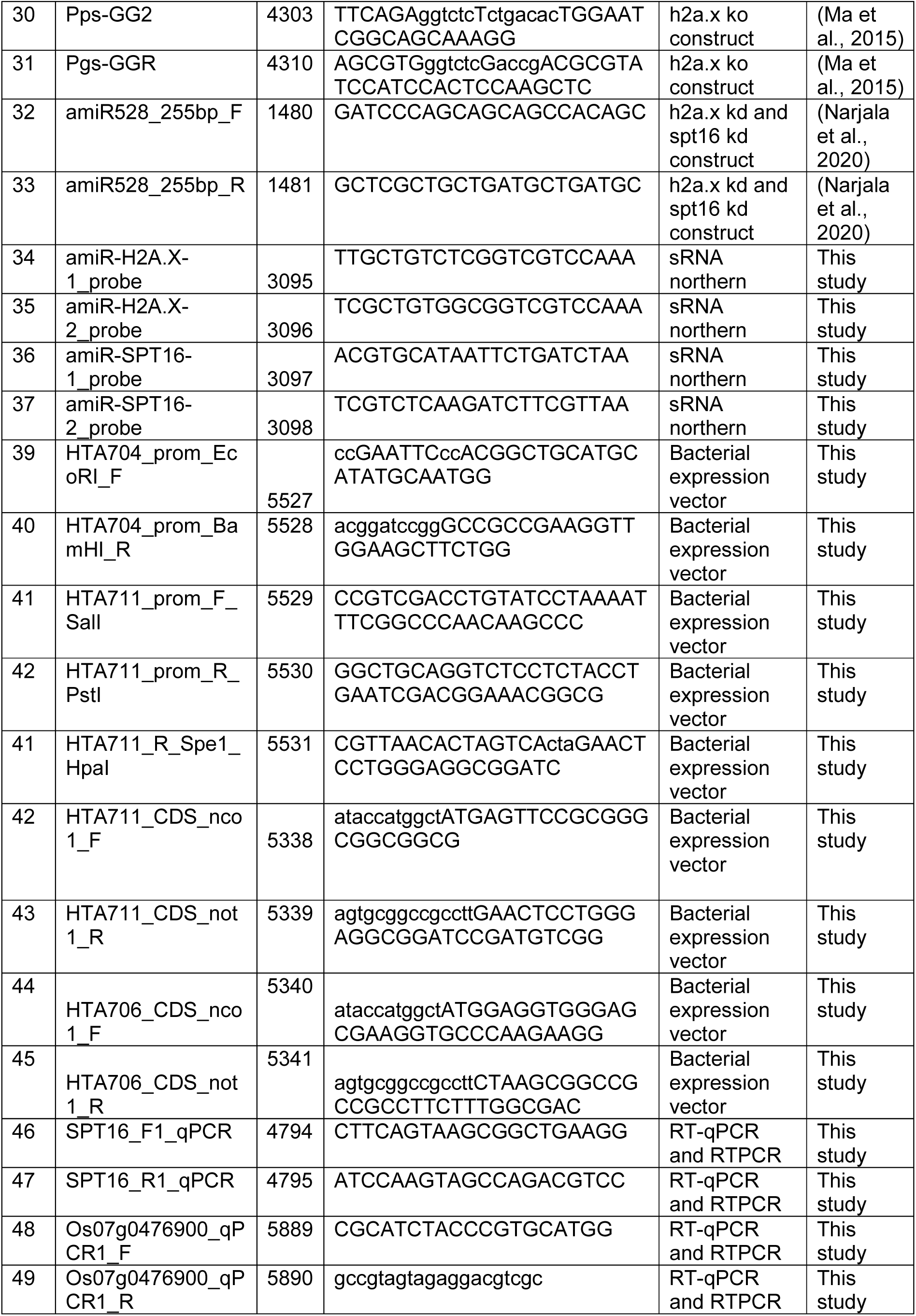
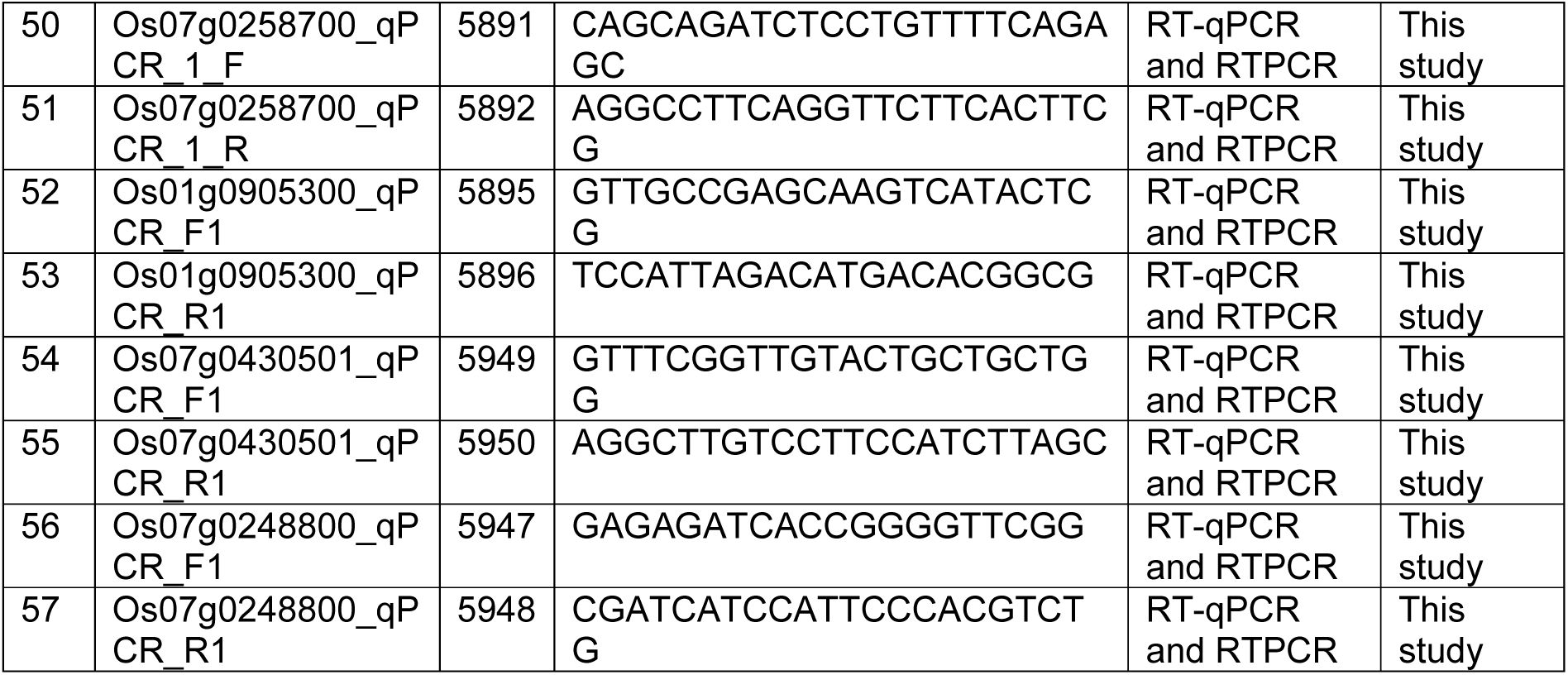
List of oligos and probes used in this study.

**Supplementary Table S4:** Details of genomic occupancy of histone variants and histone marks in rice and *Arabidopsis*.

| ChIP peaks | Promoter (%) | Exon (%) | Intron (%) | Distal intergenic region (%) | 3'UTR (%) |
| --- | --- | --- | --- | --- | --- |
| H3K4me3 | 91.14 | 0.39 | 1.03 | 6.68 | 0.17 |
| H3K4me3/ko | 90.56 | 0.48 | 1.21 | 7.5 | 0.2 |
| H3K9me2 | 11.9 | 0.11 | 4.84 | 83 | 0.15 |
| H3K9me2/ko | 10.45 | 0.09 | 4.61 | 84.69 | 0.12 |
| H2A.X | 60.9 | 4.54 | 1.99 | 28.54 | 3.48 |
| H2A.Z | 76.14 | 0.75 | 0.18 | 21.78 | 0.96 |
| H2A.W | 35.48 | 0.51 | 1.41 | 61.77 | 0.62 |
| H3 | 52.18 | 0.46 | 0.99 | 45.72 | 0.44 |
| H2A.W/ko | 35.48 | 0.51 | 1.41 | 61.77 | 0.62 |

### Supplementary Datasets S1 - S11

Supplementary Data S1. List of log2-1.5-fold change altered genes in h2a.x ko seedling

Supplementary Data S2. List of log2-1.5-fold change altered upregulated genes in H2A.X OE seedling

Supplementary Data S3. List of log2-1.5-fold change altered genes in h2a.x kd seedling

Supplementary Data S4. Gene expression of phenotype related genes used in this study

Supplementary Data S5. Expression of sRNA in h2a.x kd lost sRNA shortstack loci and associated gene expression in H2A.X mis-expression lines

Supplementary Data S6. List of log2-2-fold enriched ChIP peaks of histone marks in WT and h2a.x ko plants

Supplementary Data S7. List of log2-2-fold enriched ChIP peaks of histone variants in WT and h2a.x ko plants

Supplementary Data S8. Gene expression of histone variant bound genes in WT plants

Supplementary Data S9. List of DNA damage responsive and bleomycin responsive genes

Supplementary Data S10. List of log2-1.5-fold change altered genes in bleomycin and UV treated seedlings

Supplementary Data S11. List of log2-1.5-fold change altered genes in fact kd seedling

## References

1. Kornberg, R.D. (1974) Chromatin structure: a repeating unit of histones and DNA. Science, 184, 868–871.

2. Luger, K., Mäder, A.W., Richmond, R.K., Sargent, D.F. and Richmond, T.J. (1997) Crystal structure of the nucleosome core particle at 2.8 Å resolution. Nature, 389, 251–260.

3. Woo, Y.H. and Li, W.-H. (2012) Evolutionary conservation of histone modifications in mammals. Mol. Biol. Evol., 29, 1757–1767.

4. Kurumizaka, H., Kujirai, T. and Takizawa, Y. (2021) Contributions of histone variants in nucleosome structure and function. J. Mol. Biol., 433, 166678.

5. Vivek Hari Sundar, G., Madhu, A., Archana, A. and Shivaprasad, P.V. (2024) Plant histone variants at the nexus of chromatin readouts, stress and development. Biochim. Biophys. Acta Gen. Subj., 1868, 130539.

6. Foroozani, M., Holder, D.H. and Deal, R.B. (2022) Histone variants in the specialization of plant chromatin. Annu. Rev. Plant Biol., 73, 149–172.

7. Jamge, B., Lorković, Z.J., Axelsson, E., Osakabe, A., Shukla, V., Yelagandula, R., Akimcheva, S., Kuehn, A.L. and Berger, F. (2023) Histone variants shape chromatin states in Arabidopsis. Elife, 12.

8. Hirsch, C.D. and Springer, N.M. (2017) Transposable element influences on gene expression in plants. Biochim. Biophys. Acta Gene Regul. Mech., 1860, 157–165.

9. Pellicer, J., Hidalgo, O., Dodsworth, S. and Leitch, I.J. (2018) Genome size diversity and its impact on the evolution of land plants. Genes (Basel), 9.

10. Hammond, C.M., Strømme, C.B., Huang, H., Patel, D.J. and Groth, A. (2017) Histone chaperone networks shaping chromatin function. Nat. Rev. Mol. Cell Biol., 18, 141–158.

11. Xue, M., Ma, L., Li, X., Zhang, H., Zhao, F., Liu, Q. and Jiang, D. (2025) Single amino acid mutations in histone H3.3 illuminate the functional significance of H3K4 methylation in plants. Nat. Commun., 16, 4408.

12. Davarinejad, H., Huang, Y.-C., Mermaz, B., LeBlanc, C., Poulet, A., Thomson, G., Joly, V., Muñoz, M., Arvanitis-Vigneault, A., Valsakumar, D., et al. (2022) The histone H3.1 variant regulates TONSOKU-mediated DNA repair during replication. Science, 375, 1281–1286.

13. Deal, R.B. and Henikoff, S. (2011) Histone variants and modifications in plant gene regulation. Curr. Opin. Plant Biol., 14, 116–122.

14. Wu, X., Zhang, X., Huang, B., Han, J. and Fang, H. (2023) Advances in biological functions and mechanisms of histone variants in plants. Front. Genet., 14, 1229782.

15. Henikoff, S. and Smith, M.M. (2015) Histone variants and epigenetics. Cold Spring Harb. Perspect. Biol., 7, a019364.

16. Borg, M., Jiang, D. and Berger, F. (2021) Histone variants take center stage in shaping the epigenome. Curr. Opin. Plant Biol., 61, 101991.

17. Gandhivel, V.H.-S., Sotelo-Parrilla, P., Raju, S., Jha, S., Gireesh, A., Harshith, C.Y., Gut, F., Vinothkumar, K.R., Berger, F., Jeyaprakash, A.A., et al. (2025) An Oryza-specific histone H4 variant predisposes H4 lysine 5 acetylation to modulate salt stress responses. Nat. Plants, 11, 790–807.

18. Ravi, M. and Chan, S.W.L. (2010) Haploid plants produced by centromere-mediated genome elimination. Nature, 464, 615–618.

19. Lv, J., Yu, K., Wei, J., Gui, H., Liu, C., Liang, D., Wang, Y., Zhou, H., Carlin, R., Rich, R., et al. (2020) Generation of paternal haploids in wheat by genome editing of the centromeric histone CENH3. Nat. Biotechnol., 38, 1397–1401.

20. Marimuthu, M.P.A., Maruthachalam, R., Bondada, R., Kuppu, S., Tan, E.H., Britt, A., Chan, S.W.L. and Comai, L. (2021) Epigenetically mismatched parental centromeres trigger genome elimination in hybrids. Sci. Adv., 7, eabk1151.

21. Zhao, F., Zhang, H., Zhao, T., Li, Z. and Jiang, D. (2021) The histone variant H3.3 promotes the active chromatin state to repress flowering in Arabidopsis. Plant Physiol., 186, 2051–2063.

22. Long, X., Yang, W., Lv, Y., Zhong, X., Chen, L., Li, Q., Lv, Z., Li, Y., Cai, Y. and Yang, H. (2024) The histone variant H3.3 is required for plant growth and fertility in Arabidopsis. Int. J. Mol. Sci., 25, 2549.

23. Buttress, T., He, S., Wang, L., Zhou, S., Saalbach, G., Vickers, M., Li, G., Li, P. and Feng, X. (2022) Histone H2B.8 compacts flowering plant sperm through chromatin phase separation. Nature, 611, 614–622.

24. Yan, A., Borg, M., Berger, F. and Chen, Z. (2020) The atypical histone variant H3.15 promotes callus formation in Arabidopsis thaliana. Development, 147, dev184895.

25. Borg, M., Jacob, Y., Susaki, D., LeBlanc, C., Buendía, D., Axelsson, E., Kawashima, T., Voigt, P., Boavida, L., Becker, J., et al. (2020) Targeted reprogramming of H3K27me3 resets epigenetic memory in plant paternal chromatin. Nat. Cell Biol., 22, 621–629.

26. Nunez-Vazquez, R., Madeira, S., Rodríguez-Casillas, L., Gomez-Martinez, D., Desvoyes, B. and Gutierrez, C. (2025) The histone variant H3.14 is an early player in the abiotic stress response of Arabidopsis. Dev. Cell, 10.1016/j.devcel.2025.06.019.

27. Zhao, T., Lu, J., Zhang, H., Xue, M., Pan, J., Ma, L., Berger, F. and Jiang, D. (2022) Histone H3.3 deposition in seed is essential for the post-embryonic developmental competence in Arabidopsis. Nat. Commun., 13, 7728.

28. Wollmann, H., Stroud, H., Yelagandula, R., Tarutani, Y., Jiang, D., Jing, L., Jamge, B., Takeuchi, H., Holec, S., Nie, X., et al. (2017) The histone H3 variant H3.3 regulates gene body DNA methylation in Arabidopsis thaliana. Genome Biol., 18, 94.

29. Lorković, Z.J., Park, C., Goiser, M., Jiang, D., Kurzbauer, M.-T., Schlögelhofer, P. and Berger, F. (2017) Compartmentalization of DNA damage response between heterochromatin and euchromatin is mediated by distinct H2A histone variants. Curr. Biol., 27, 1192–1199.

30. Thatcher, T.H. and Gorovsky, M.A. (1994) Phylogenetic analysis of the core histones H2A, H2B, H3, and H4. Nucleic Acids Res., 22, 174–179.

31. Coleman-Derr, D. and Zilberman, D. (2012) Deposition of histone variant H2A.Z within gene bodies regulates responsive genes. PLoS Genet., 8, e1002988.

32. Zilberman, D., Coleman-Derr, D., Ballinger, T. and Henikoff, S. (2008) Histone H2A.Z and DNA methylation are mutually antagonistic chromatin marks. Nature, 456, 125–129.

33. Zemach, A., McDaniel, I.E., Silva, P. and Zilberman, D. (2010) Genome-wide evolutionary analysis of eukaryotic DNA methylation. Science, 328, 916–919.

34. Jarillo, J.A. and Piñeiro, M. (2015) H2A.Z mediates different aspects of chromatin function and modulates flowering responses in Arabidopsis. Plant J., 83, 96–109.

35. Kumar, S.V. and Wigge, P.A. (2010) H2A.Z-containing nucleosomes mediate the thermosensory response in Arabidopsis. Cell, 140, 136–147.

36. Zhang, H., Roberts, D.N. and Cairns, B.R. (2005) Genome-wide dynamics of Htz1, a histone H2A variant that poises repressed/basal promoters for activation through histone loss. Cell, 123, 219–231.

37. Sura, W., Kabza, M., Karlowski, W.M., Bieluszewski, T., Kus-Slowinska, M., Pawełoszek, Ł., Sadowski, J. and Ziolkowski, P.A. (2017) Dual role of the histone variant H2A.Z in transcriptional regulation of stress-response genes. Plant Cell, 29, 791–807.

38. Dai, X., Bai, Y., Zhao, L., Dou, X., Liu, Y., Wang, L., Li, Y., Li, W., Hui, Y., Huang, X., et al. (2018) H2A.Z represses gene expression by modulating promoter nucleosome structure and enhancer histone modifications in Arabidopsis. Mol. Plant, 11, 635.

39. March-Díaz, R., García-Domínguez, M., Lozano-Juste, J., León, J., Florencio, F.J. and Reyes, J.C. (2008) Histone H2A.Z and homologues of components of the SWR1 complex are required to control immunity in Arabidopsis. Plant J., 53, 475–487.

40. Deal, R.B., Kandasamy, M.K., McKinney, E.C. and Meagher, R.B. (2005) The nuclear actin-related protein ARP6 is a pleiotropic developmental regulator required for the maintenance of FLOWERING LOCUS C expression and repression of flowering in Arabidopsis. Plant Cell, 17, 2633–2646.

41. Deal, R.B., Topp, C.N., McKinney, E.C. and Meagher, R.B. (2007) Repression of flowering in Arabidopsis requires activation of FLOWERING LOCUS C expression by the histone variant H2A.Z. Plant Cell, 19, 74–83.

42. Xue, M., Zhang, H., Zhao, F., Zhao, T., Li, H. and Jiang, D. (2021) The INO80 chromatin remodeling complex promotes thermomorphogenesis by connecting H2A.Z eviction and active transcription in Arabidopsis. Mol. Plant, 14, 1799–1813.

43. Gómez-Zambrano, Á., Merini, W. and Calonje, M. (2019) The repressive role of Arabidopsis H2A.Z in transcriptional regulation depends on AtBMI1 activity. Nat. Commun., 10, 2828.

44. Choi, K., Park, C., Lee, J., Oh, M., Noh, B. and Lee, I. (2007) Arabidopsis homologs of components of the SWR1 complex regulate flowering and plant development. Development, 134, 1931–1941.

45. Yelagandula, R., Stroud, H., Holec, S., Zhou, K., Feng, S., Zhong, X., Muthurajan, U.M., Nie, X., Kawashima, T., Groth, M., et al. (2014) The histone variant H2A.W defines heterochromatin and promotes chromatin condensation in Arabidopsis. Cell, 158, 98–109.

46. Bourguet, P., Picard, C.L., Yelagandula, R., Pélissier, T., Lorković, Z.J., Feng, S., Pouch-Pélissier, M.-N., Schmücker, A., Jacobsen, S.E., Berger, F., et al. (2021) The histone variant H2A.W and linker histone H1 co-regulate heterochromatin accessibility and DNA methylation. Nat. Commun., 12, 2683.

47. Osakabe, A., Jamge, B., Axelsson, E., Montgomery, S.A., Akimcheva, S., Kuehn, A.L., Pisupati, R., Lorković, Z.J., Yelagandula, R., Kakutani, T., et al. (2021) The chromatin remodeler DDM1 prevents transposon mobility through deposition of histone variant H2A.W. Nat. Cell Biol., 23, 391–400.

48. Wang, Y., Wu, J., Yang, S., Li, X., Wang, J., Lv, Q., Zhu, X., Lu, G., Zhang, J., Shen, W.-H., et al. (2025) Structural and functional interrelationships of histone H2A with its variants H2A.Z and H2A.W in Arabidopsis. Structure, 33, 1240–1249.e5.

49. Zemach, A., Kim, M.Y., Hsieh, P.-H., Coleman-Derr, D., Eshed-Williams, L., Thao, K., Harmer, S.L. and Zilberman, D. (2013) The Arabidopsis nucleosome remodeler DDM1 allows DNA methyltransferases to access H1-containing heterochromatin. Cell, 153, 193–205.

50. Choi, J., Lyons, D.B. and Zilberman, D. (2021) Histone H1 prevents non-CG methylation-mediated small RNA biogenesis in Arabidopsis heterochromatin. Elife, 10.

51. Rogakou, E.P., Pilch, D.R., Orr, A.H., Ivanova, V.S. and Bonner, W.M. (1998) DNA double-stranded breaks induce histone H2AX phosphorylation on serine 139. J. Biol. Chem., 273, 5858–5868.

52. Turinetto, V. and Giachino, C. (2015) Multiple facets of histone variant H2AX: a DNA double-strand-break marker with several biological functions. Nucleic Acids Res., 43, 2489–2498.

53. Piquet, S., Le Parc, F., Bai, S.-K., Chevallier, O., Adam, S. and Polo, S.E. (2018) The histone chaperone FACT coordinates H2A.X-dependent signaling and repair of DNA damage. Mol. Cell, 72, 888–901.e7.

54. Heo, K., Kim, H., Choi, S.H., Choi, J., Kim, K., Gu, J., Lieber, M.R., Yang, A.S. and An, W. (2008) FACT-mediated exchange of histone variant H2AX regulated by phosphorylation of H2AX and ADP-ribosylation of Spt16. Mol. Cell, 30, 86–97.

55. Frost, J.M., Lee, J., Hsieh, P.-H., Lin, S.J.H., Min, Y., Bauer, M., Runkel, A.M., Cho, H.-T., Hsieh, T.-F., Fischer, R.L., et al. (2023) H2A.X promotes endosperm-specific DNA methylation in Arabidopsis thaliana. BMC Plant Biol., 23, 585.

56. Zhang, Z., Zhang, F. and Xiong, T. (2025) Evolution of the chromatin remodeling complex FACT: Functional analysis of SSRP1 and SPT16 in early anther development. Int. J. Biol. Macromol., 284, 138167.

57. Ikeda, Y., Kinoshita, Y., Susaki, D., Ikeda, Y., Iwano, M., Takayama, S., Higashiyama, T., Kakutani, T. and Kinoshita, T. (2011) HMG domain containing SSRP1 is required for DNA demethylation and genomic imprinting in Arabidopsis. Dev. Cell, 21, 589–596.

58. Duroux, M., Houben, A., Růzicka, K., Friml, J. and Grasser, K.D. (2004) The chromatin remodelling complex FACT associates with actively transcribed regions of the Arabidopsis genome. Plant J., 40, 660–671.

59. Lolas, I.B., Himanen, K., Grønlund, J.T., Lynggaard, C., Houben, A., Melzer, M., Van Lijsebettens, M. and Grasser, K.D. (2010) The transcript elongation factor FACT affects Arabidopsis vegetative and reproductive development and genetically interacts with HUB1/2. Plant J., 61, 686–697.

60. Orphanides, G., Wu, W.H., Lane, W.S., Hampsey, M. and Reinberg, D. (1999) The chromatin-specific transcription elongation factor FACT comprises human SPT16 and SSRP1 proteins. Nature, 400, 284–288.

61. Friesner, J.D., Liu, B., Culligan, K. and Britt, A.B. (2005) Ionizing radiation–dependent γ-H2AX focus formation requires ataxia telangiectasia mutated and ataxia telangiectasia mutated and Rad3-related. Mol. Biol. Cell, 16, 2566–2576.

62. Stucki, M., Clapperton, J.A., Mohammad, D., Yaffe, M.B., Smerdon, S.J. and Jackson, S.P. (2008) MDC1 directly binds phosphorylated histone H2AX to regulate cellular responses to DNA double-strand breaks. Cell, 133, 549.

63. Lou, Z., Minter-Dykhouse, K., Franco, S., Gostissa, M., Rivera, M.A., Celeste, A., Manis, J.P., van Deursen, J., Nussenzweig, A., Paull, T.T., et al. (2006) MDC1 maintains genomic stability by participating in the amplification of ATM-dependent DNA damage signals. Mol. Cell, 21, 187–200.

64. Lorković, Z.J., Klingenbrunner, M., Cho, C.H. and Berger, F. (2024) Identification of plants’ functional counterpart of the metazoan mediator of DNA Damage checkpoint 1. EMBO Rep., 25, 1936–1961.

65. Fan, T., Kang, H., Wu, D., Zhu, X., Huang, L., Wu, J. and Zhu, Y. (2022) Arabidopsis γ-H2A.X-INTERACTING PROTEIN participates in DNA damage response and safeguards chromatin stability. Nat. Commun., 13, 7942.

66. Waterworth, W.M., Wilson, M., Wang, D., Nuhse, T., Warward, S., Selley, J. and West, C.E. (2019) Phosphoproteomic analysis reveals plant DNA damage signalling pathways with a functional role for histone H2AX phosphorylation in plant growth under genotoxic stress. Plant J., 100, 1007–1021.

67. Guo, P., Wang, T.-J., Wang, S., Peng, X., Kim, D.H. and Liu, Y. (2024) Arabidopsis histone variant H2A.X functions in the DNA damage-coupling abscisic acid signaling pathway. Int. J. Mol. Sci., 25, 8940.

68. Dobersch, S., Rubio, K., Singh, I., Günther, S., Graumann, J., Cordero, J., Castillo-Negrete, R., Huynh, M.B., Mehta, A., Braubach, P., et al. (2021) Positioning of nucleosomes containing γ-H2AX precedes active DNA demethylation and transcription initiation. Nat. Commun., 12, 1072.

69. Xiao, S., Jiang, L., Wang, C. and Ow, D.W. (2021) Arabidopsis OXS3 family proteins repress ABA signaling through interactions with AFP1 in the regulation of ABI4 expression. J. Exp. Bot., 72, 5721–5734.

70. Wyrick, J.J. and Parra, M.A. (2009) The role of histone H2A and H2B post-translational modifications in transcription: a genomic perspective. Biochim. Biophys. Acta, 1789, 37–44.

71. Parra, M.A. and Wyrick, J.J. (2007) Regulation of gene transcription by the histone H2A N-terminal domain. Mol. Cell. Biol., 27, 7641–7648.

72. Narjala, A., Nair, A., Tirumalai, V., Hari Sundar, G.V. and Shivaprasad, P.V. (2020) A conserved sequence signature is essential for robust plant miRNA biogenesis. Nucleic Acids Res., 48, 3103–3118.

73. Anushree, N. and Shivaprasad, P.V. (2017) Regulation of Plant miRNA Biogenesis. Proc. Indian Natl. Sci. Acad. (A Phys. Sci.), 95.

74. Kawahara, Y., Oono, Y., Wakimoto, H., Ogata, J., Kanamori, H., Sasaki, H., Mori, S., Matsumoto, T. and Itoh, T. (2016) TENOR: Database for comprehensive mRNA-Seq experiments in rice. Plant Cell Physiol., 57, e7.

75. Frost, J.M., Lee, J., Hsieh, P.-H., Lin, S.J.H., Min, Y., Bauer, M., Runkel, A.M., Cho, H.-T., Hsieh, T.-F., Fischer, R.L., et al. (2023) H2A.X promotes endosperm-specific DNA methylation in Arabidopsis thaliana. Res. Sq., 10.21203/rs.3.rs-2974671/v1.

76. Hu, Y., Lai, Y., Chen, X., Zhou, D.-X. and Zhao, Y. (2020) Distribution pattern of histone marks potentially determines their roles in transcription and RNA processing in rice. J. Plant Physiol., 249, 153167.

77. Zhao, L., Xie, L., Zhang, Q., Ouyang, W., Deng, L., Guan, P., Ma, M., Li, Y., Zhang, Y., Xiao, Q., et al. (2020) Integrative analysis of reference epigenomes in 20 rice varieties. Nat. Commun., 11, 2658.

78. Bennetzen, J.L. and Wang, H. (2014) The contributions of transposable elements to the structure, function, and evolution of plant genomes. Annu. Rev. Plant Biol., 65, 505–530.

79. Espinas, N.A., Tu, L.N., Furci, L., Shimajiri, Y., Harukawa, Y., Miura, S., Takuno, S. and Saze, H. (2020) Transcriptional regulation of genes bearing intronic heterochromatin in the rice genome. PLoS Genet., 16, e1008637.

80. Lei, B. and Berger, F. (2020) H2A variants in Arabidopsis: Versatile regulators of genome activity. Plant Commun., 1, 100015.

81. VanDemark, A.P., Xin, H., McCullough, L., Rawlins, R., Bentley, S., Heroux, A., Stillman, D.J., Hill, C.P. and Formosa, T. (2008) Structural and functional analysis of the Spt16p N-terminal domain reveals overlapping roles of yFACT subunits. J. Biol. Chem., 283, 5058–5068.

82. Shalmani, A., Ullah, U., Tai, L., Zhang, R., Jing, X.-Q., Muhammad, I., Bhanbhro, N., Liu, W.-T., Li, W.-Q. and Chen, K.-M. (2023) OsBBX19-OsBTB97/OsBBX11 module regulates spikelet development and yield production in rice. Plant Sci., 334, 111779.

83. Niu, B., Xu, J., Zhiguo, Zhang, Z., Lu, X. and Chen, C. (2023) Ectopic expression of OsNF-YA8, an endosperm-specific nuclear factor Y transcription-factor gene, causes vegetative and reproductive development defects in rice. Crop J., 10.1016/j.cj.2023.07.001.

84. Shan, Q., Guan, J., Yang, Y., Chai, T., Gong, S., Wang, J. and Qiao, K. (2024) Cadmium-induced protein AS8: A protein to improve Cd accumulation and transport via Cd uptake in poplar. Plant Physiol. Biochem., 216, 109199.

85. Lee, S.-K., Eom, J.-S., Hwang, S.-K., Shin, D., An, G., Okita, T.W. and Jeon, J.-S. (2016) Plastidic phosphoglucomutase and ADP-glucose pyrophosphorylase mutants impair starch synthesis in rice pollen grains and cause male sterility. J. Exp. Bot., 67, 5557–5569.

86. Wang, J., Li, M., Nan, N., Ma, A., Ao, M., Yu, J., Wang, X., Han, K., Yun, D.-J., Liu, B., et al. (2024) OsGADD45a1: a multifaceted regulator of rice architecture, grain yield, and blast resistance. Plant Cell Rep., 43, 88.

87. Ma, B., Cao, X., Li, X., Bian, Z., Zhang, Q.-Q., Fang, Z., Liu, J., Li, Q., Liu, Q., Zhang, L., et al. (2024) Two ABCI family transporters, OsABCI15 and OsABCI16, are involved in grain-filling in rice. J. Genet. Genomics, 51, 492–506.

88. Le, H., Simmons, C.H. and Zhong, X. (2025) Functions and mechanisms of histone modifications in plants. Annu. Rev. Plant Biol., 76, 551–578.

89. Kawashima, T., Lorković, Z.J., Nishihama, R., Ishizaki, K., Axelsson, E., Yelagandula, R., Kohchi, T. and Berger, F. (2015) Diversification of histone H2A variants during plant evolution. Trends Plant Sci., 20, 419–425.

90. Shi, S., Wang, T., Chen, Z., Tang, Z., Wu, Z., Salt, D.E., Chao, D.-Y. and Zhao, F.-J. (2016) OsHAC1;1 and OsHAC1;2 function as arsenate reductases and regulate arsenic accumulation. Plant Physiol., 172, 1708–1719.

91. Li, X., Chen, R., Chu, Y., Huang, J., Jin, L., Wang, G. and Huang, J. (2018) Overexpression of RCc3 improves root system architecture and enhances salt tolerance in rice. Plant Physiol. Biochem., 130, 566–576.

92. Kang, Z., Qin, T. and Zhao, Z. (2019) Overexpression of the zinc finger protein gene OsZFP350 improves root development by increasing resistance to abiotic stress in rice. Acta Biochim. Pol., 66, 183–190.

93. Ding, N., Cai, J., Xiao, S. and Jiang, L. (2024) Heterologous expression of rice OsEXO70FX1 confers tolerance to cadmium in Arabidopsis thaliana and fission yeast. Plant Physiol. Biochem., 206, 108268.

94. Ye, N., Yang, G., Chen, Y., Zhang, C., Zhang, J. and Peng, X. (2014) Two hydroxypyruvate reductases encoded by OsHPR1 and OsHPR2 are involved in photorespiratory metabolism in rice. J. Integr. Plant Biol., 56, 170–180.

95. Broin, M., Cuiné, S., Peltier, G. and Rey, P. (2000) Involvement of CDSP 32, a drought-induced thioredoxin, in the response to oxidative stress in potato plants. FEBS Lett., 467, 245–248.

96. Wang, M., He, Y., Zhong, Z., Papikian, A., Wang, S., Gardiner, J., Ghoshal, B., Feng, S., Jami-Alahmadi, Y., Wohlschlegel, J.A., et al. (2025) Histone H3 lysine 4 methylation recruits DNA demethylases to enforce gene expression in Arabidopsis. Nat. Plants, 11, 206–217.

97. Zhu, K., Chen, J., Zhao, L., Lu, F., Deng, J., Lin, X., He, C., Wagner, D. and Xiao, J. (2025) Dynamic control of H2A.Zub and H3K27me3 by ambient temperature during cell fate determination in Arabidopsis. Dev. Cell, 60, 2192–2208.e5.

98. Osakabe, A., Lorkovic, Z.J., Kobayashi, W., Tachiwana, H., Yelagandula, R., Kurumizaka, H. and Berger, F. (2018) Histone H2A variants confer specific properties to nucleosomes and impact on chromatin accessibility. Nucleic Acids Res., 46, 7675–7685.

99. Choi, J., Hyun, Y., Kang, M.-J., In Yun, H., Yun, J.-Y., Lister, C., Dean, C., Amasino, R.M., Noh, B., Noh, Y.-S., et al. (2009) Resetting and regulation of Flowering Locus C expression during Arabidopsis reproductive development. Plant J., 57, 918–931.

100. Bratzel, F., López-Torrejón, G., Koch, M., Del Pozo, J.C. and Calonje, M. (2010) Keeping cell identity in Arabidopsis requires PRC1 RING-finger homologs that catalyze H2A monoubiquitination. Curr. Biol., 20, 1853–1859.

101. Vissers, J.H., Nicassio, F., van Lohuizen, M., Di Fiore, P.P. and Citterio, E. (2008) The many faces of ubiquitinated histone H2A: insights from the DUBs. Cell Div., 3, 8.

102. Su, X.-M., Yuan, D.-Y., Liu, N., Zhang, Z.-C., Yang, M., Li, L., Chen, S., Zhou, Y. and He, X.-J. (2025) ALFIN-like proteins link histone H3K4me3 to H2A ubiquitination and coordinate diverse chromatin modifications in Arabidopsis. Mol. Plant, 18, 130–150.

103. Xu, L., Wang, Y., Li, X., Hu, Q., Adamkova, V., Xu, J., Harris, C.J. and Ausin, I. (2025) H3K4me3 binding ALFIN-LIKE proteins recruit SWR1 for gene-body deposition of H2A.Z. Genome Biol., 26, 137.

104. Michl-Holzinger, P., Mortensen, S.A. and Grasser, K.D. (2019) The SSRP1 subunit of the histone chaperone FACT is required for seed dormancy in Arabidopsis. J. Plant Physiol., 236, 105–108.

105. Pfab, A., Breindl, M. and Grasser, K.D. (2018) The Arabidopsis histone chaperone FACT is required for stress-induced expression of anthocyanin biosynthetic genes. Plant Mol. Biol., 96, 367–374.

106. Formosa, T. (2013) The role of FACT in making and breaking nucleosomes. Biochim. Biophys. Acta, 1819, 247–255.

107. Nielsen, M., Ard, R., Leng, X., Ivanov, M., Kindgren, P., Pelechano, V. and Marquardt, S. (2019) Transcription-driven chromatin repression of Intragenic transcription start sites. PLoS Genet., 15, e1007969.

108. Zhang, W., Cheng, L., Li, K., Xie, L., Ji, J., Lei, X., Jiang, A., Chen, C., Li, H., Li, P., et al. (2024) Evolutional heterochromatin condensation delineates chromocenter formation and retrotransposon silencing in plants. Nat. Plants, 10, 1215–1230.

109. Quesneville, H. (2020) Twenty years of transposable element analysis in the Arabidopsis thaliana genome. Mob. DNA, 11, 28.

110. Harris, C.J., Zhong, Z., Ichino, L., Feng, S. and Jacobsen, S.E. (2024) H1 restricts euchromatin-associated methylation pathways from heterochromatic encroachment. Elife, 12.

111. Ma, X., Zhang, Q., Zhu, Q., Liu, W., Chen, Y., Qiu, R., Wang, B., Yang, Z., Li, H., Lin, Y., et al. (2015) A robust CRISPR/Cas9 system for convenient, high-efficiency multiplex genome editing in monocot and dicot plants. Mol. Plant, 8, 1274–1284.

112. Hiei, Y., Ohta, S., Komari, T. and Kumashiro, T. (1994) Efficient transformation of rice (Oryza sativa L.) mediated by Agrobacterium and sequence analysis of the boundaries of the T-DNA. Plant J., 6, 271–282.

113. Sridevi, G., Sabapathi, N., Meena, P., Nandakumar, R., Samiyappan, R., Muthukrishnan, S. and Veluthambi, K. (2003) Transgenic indica rice variety Pusa basmati 1 constitutively expressing a rice chitinase gene exhibits enhanced resistance to Rhizoctonia solani. J. Plant Biochem. Biotechnol., 12, 93–101.

114. Swetha, C., Basu, D., Pachamuthu, K., Tirumalai, V., Nair, A., Prasad, M. and Shivaprasad, P.V. (2018) Major domestication-related phenotypes in Indica rice are due to loss of miRNA-mediated laccase silencing. Plant Cell, 30, 2649–2662.

115. Tamura, K., Stecher, G. and Kumar, S. (2021) MEGA11: Molecular Evolutionary Genetics Analysis version 11. Mol. Biol. Evol., 38, 3022–3027.

116. Ramanathan, V. and Veluthambi, K. (1995) Transfer of non-T-DNA portions of the Agrobacterium tumefaciens Ti plasmid pTiA6 from the left terminus of TL-DNA. Plant Mol. Biol., 28, 1149–1154.

117. Sundar, H. and Shivaprasad, G.V. (2022) Investigation of transposon DNA methylation and copy number variation in plants using Southern hybridisation. Bio Protoc, 12.

118. Shivaprasad, P.V., Chen, H.-M., Patel, K., Bond, D.M., Santos, B.A.C.M. and Baulcombe, D.C. (2012) A microRNA superfamily regulates nucleotide binding site-leucine-rich repeats and other mRNAs. Plant Cell, 24, 859–874.

119. Tirumalai, V., Prasad, M. and Shivaprasad, P.V. (2020) RNA blot analysis for the detection and quantification of plant MicroRNAs. J. Vis. Exp., 10.3791/61394.

120. Methods for determining leaf chlorophyll content of rice: A reappraisal October 1996 (1996) Indian Journal of Experimental Biology, 34, 1030–1033.

121. Nakata, M. and Ohme-Takagi, M. (2014) Quantification of Anthocyanin Content. Bio Protoc., 4.

122. Tirumalai, V., Narjala, A., Swetha, C., Sundar, G.V.H., Sujith, T.N. and Shivaprasad, P.V. (2022) Cultivar-specific miRNA-mediated RNA silencing in grapes. Planta, 256, 17.

123. Ghandhivel, S. (2023) Plant polymerase IV sensitizes chromatin through histone modifications to preclude spread of silencing into protein-coding domains. Genome Res, 33, 715–728.

124. Harshith, C.Y., Pal, A., Chakraborty, M., Nair, A., Raju, S. and Shivaprasad, P.V. (2024) Wound-induced small-peptide-mediated signaling cascade, regulated by OsPSKR, dictates balance between growth and defense in rice. Cell Rep., 43, 114515.

125. Bolger, A.M., Lohse, M. and Usadel, B. (2014) Trimmomatic: a flexible trimmer for Illumina sequence data. Bioinformatics, 30, 2114–2120.

126. Kim, D., Langmead, B. and Salzberg, S.L. (2015) HISAT: a fast spliced aligner with low memory requirements. Nat. Methods, 12, 357–360.

127. Ge, S.X., Jung, D. and Yao, R. (2020) ShinyGO: a graphical gene-set enrichment tool for animals and plants. Bioinformatics, 36, 2628–2629.

128. Yao, L., Wang, H., Song, Y. and Sui, G. (2017) BioQueue: a novel pipeline framework to accelerate bioinformatics analysis. Bioinformatics, 33, 3286–3288.

129. Tian, T., Liu, Y., Yan, H., You, Q., Yi, X., Du, Z., Xu, W. and Su, Z. (2017) agriGO v2.0: a GO analysis toolkit for the agricultural community, 2017 update. Nucleic Acids Res., 45, W122–W129.

130. Pal, A.K., Gandhivel, V.H.-S., Nambiar, A.B. and Shivaprasad, P.V. (2024) Upstream regulator of genomic imprinting in rice endosperm is a small RNA-associated chromatin remodeler. Nat. Commun., 15, 7807.

131. Martin, M. (2011) Cutadapt removes adapter sequences from high-throughput sequencing reads. EMBnet J., 17, 10.

132. Krueger, F. and Andrews, S.R. (2011) Bismark: a flexible aligner and methylation caller for Bisulfite-Seq applications. Bioinformatics, 27, 1571–1572.

133. Huang, X., Zhang, S., Li, K., Thimmapuram, J. and Xie, S. (2018) ViewBS: a powerful toolkit for visualization of high-throughput bisulfite sequencing data. Bioinformatics, 34, 708–709.

134. Nair, A., Chatterjee, K.S., Jha, V., Das, R. and Shivaprasad, P.V. (2020) Stability of Begomoviral pathogenicity determinant βC1 is modulated by mutually antagonistic SUMOylation and SIM interactions. BMC Biol., 18, 110.

135. Langmead, B. and Salzberg, S.L. (2012) Fast gapped-read alignment with Bowtie 2. Nat. Methods, 9, 357–359.

136. Ramírez, F., Dündar, F., Diehl, S., Grüning, B.A. and Manke, T. (2014) deepTools: a flexible platform for exploring deep-sequencing data. Nucleic Acids Res., 42, W187–91.

137. Wickham, H. (2011) Ggplot2. Wiley Interdiscip. Rev. Comput. Stat., 3, 180–185.

